# Glial TGFβ activity promotes axon survival in peripheral nerves

**DOI:** 10.1101/2021.09.02.458753

**Authors:** Alexandria P. Lassetter, Megan M. Corty, Romina Barria, Amy E. Sheehan, Jo Hill, Sue A. Aicher, A. Nicole Fox, Marc R. Freeman

**Affiliations:** Vollum Institute, Oregon Health & Science University, Portland, OR, 97239, USA; Department of Chemical Physiology & Biochemistry, Oregon Health & Science University, Portland, OR, 97239, USA; University of Massachusetts Medical School, Worcester, MA, 01655, USA

**Keywords:** Axon, glia, peripheral nervous system, nerves, TGFβ, degeneration

## Abstract

Axons can represent the majority of the volume of a neuron and are energetically very demanding. Specialized glia ensheathe axons and are believed to support axon function and maintenance throughout life, but molecular details of glia-neuron support mechanisms remain poorly defined. Here we identify a collection of secreted and transmembrane genes that are required in glia for long-term axon survival *in vivo*. We show that key components of the TGFβ superfamily are required cell-autonomously in glia for peripheral nerve maintenance, although their loss does not disrupt glial morphology. We observe age-dependent neurodegeneration in the absence of glial TGFβ signaling that can be rescued by genetic blockade of Wallerian degeneration. Our data argue that glial TGFβ signaling normally acts to promote axon survival and suppress neurodegeneration.

**Significance Statement:** Axon maintenance is critical to preserving the functional integrity of the nervous system across animal lifespan. Glia contribute to axon maintenance, but their precise roles remain to be fully characterized. We identify glial genes that regulate axon support and provide new molecular insight into the means by which glia promote axon survival, which may help explain why neurodegeneration occurs when glia are lost in disease. We show that TGFβ signaling in mature glia is essential for long-term maintenance of axons, and that loss of TGFβ signaling activates an axon death signaling pathway.

## Introduction

Axons connect distant structures in the nervous system and maintaining axon integrity is crucial to preserve neural circuit function. Axons can traverse great distances across the body and have complex morphologies, such that the majority of a neuron’s volume is contained within the axon. Axons in the human sciatic nerve can be a meter long (Jessen & Mirsky, 2005), and axons of single dopaminergic neurons in the substantia nigra pars compacta branch so profusely that their estimated cumulative linear length can be up to nearly 800 mm (Matsuda et al., 2009). Sustaining axons presents a significant challenge for the neuron both in terms of their size, and the substantial metabolic demand required for signal transduction (Harris et al., 2012).

Glial cells that ensheathe axons are believed to serve as a local, external sources of support to axons, particularly at great distances from the cell body. Evidence from human diseases such as multiple sclerosis (MS) or Charcot-Marie-Tooth (CMT), where loss of glia or glial molecules causes neurodegeneration, further implicate glia in the support of axon maintenance (Brennan et al., 2015; Kornek et al., 2000; Kuhlmann et al., 2002; Trapp et al., 1998). Disrupting a single glial gene causes neurodegeneration in the absence of glial cell loss (Griffiths et al., 1998; Lappe-Siefke et al., 2003). A growing body of work has established that glia provide significant metabolic support to axons (Fünfschilling et al., 2012; Volkenhoff et al., 2015) and more recently, defense against iron-mediated toxicity (Mukherjee et al., 2020). However, a comprehensive mechanistic understanding of glial support of axons *in vivo* remains elusive.

To identify mechanisms by which glia support axon maintenance *in vivo*, we conducted a screen in *Drosophila* to identify glial molecules required to sustain long-term axon survival. We identified over 200 genes that—when depleted from glia—lead to axon degeneration or animal lethality. Interestingly, we found that glial loss of the majority of TGFβ superfamily molecules, a multi-functional family of molecules that regulate diverse aspects of metazoan growth, differentiation and homeostasis (Upadhyay et al., 2017), resulted in age-dependent axon loss and cell body death in sensory neurons in the peripheral nervous system. We provide evidence that glial TGFβ signaling is required to support axons directly (versus supporting the cell body), since all neurodegeneration after glial TGFβ depletion can be suppressed by supplying the axon survival factor Wallerian degeneration Slow (Wld^S^) to neurons. These data identify the TGFβ superfamily as a key regulator of *in vivo* glial functions in supporting long-term maintenance of axons.

## Results

### Ablation of wrapping glia leads to axon degeneration

We aimed to explore the mechanisms by which glia support axon function and survival. We used the adult L1 wing nerve of *Drosophila melanogaster* as a model, where it is possible to independently manipulate neurons and glia and examine their morphology with single cell/axon resolution *in vivo* (Fig. 1A-B). This sensory nerve contains roughly 280 sensory neurons (Fig. 1C). Their cell bodies are positioned along the anterior wing margin, and they project their axons into the thorax (Fig. 1A; (Palka et al., 1983). These are among the longest axons in *Drosophila* (Fig. 1A; (Palka et al., 1983) and each is individually ensheathed by glia along their length (Fig. 1B & C): wrapping glia (WG) cover the entire nerve and interdigitate into the axon bundle separating axons from one another (Fig. 1C; (Neukomm et al., 2014). Given their length, and their relatively unusually extensive ensheathment by *Drosophila* glia, we speculated that the glia surrounding these neurons would be a good model to explore how glia provide essential support for neuronal function and maintenance.

**Fig. 1:**
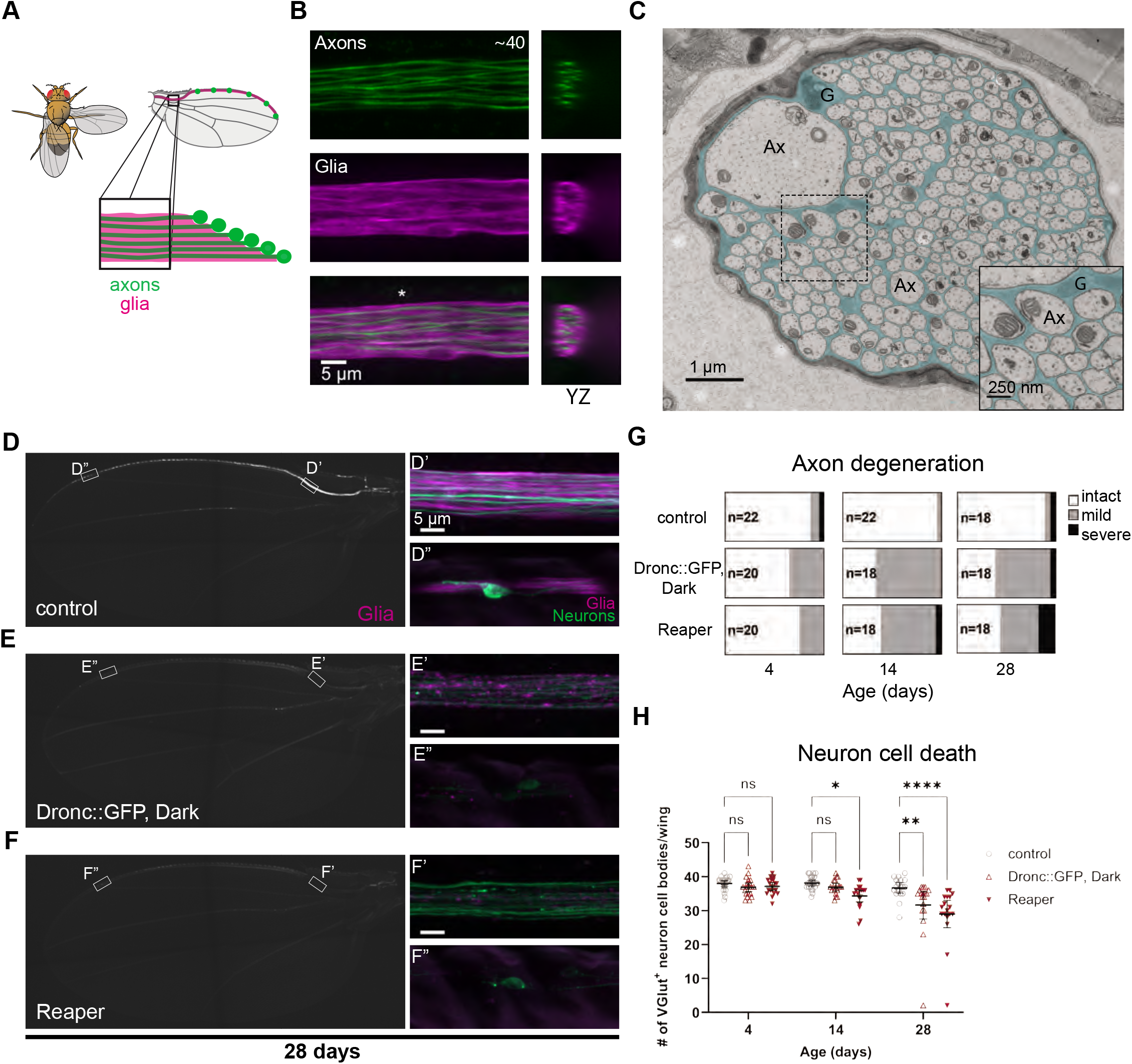
Ablating wrapping glia results in neurodegeneration in the peripheral nerve of the wing with age. A) Diagram of the sensory nerve in the wing of *Drosophila* B) Images from the area depicted in the box in A. A subset of glutamatergic neurons are genetically labeled with GFP and glia are labelled with tdTomato. The orthogonal fluorescent image corresponds to the location at the asterisk. C) Electron micrograph of a cross section of the nerve in the wing from the same region as in A. Wrapping glial membrane is psuedocolored in cyan. Example glia (G) and axons (Ax) are labeled. D-F) Representative images of control and glial-ablated wings at 28 days of age with subset of wrapping glia labeled with tdTomato. D-F’ & D-F”) Higher magnification images from ROIs in D-F. G) Classification of axon phenotype for each condition categorized into intact, mild, or severe. n=number of wings. H) Quantification of the number of intact neuron cell bodies at each time point. Images are maximum intensity projections derived from z-stack images. Data are represented as mean ± 95% CI. Statistics: Two-way ANOVA with Dunnett’s multiple comparisons test. Significance: *= p < 0.05, **= p < 0.01, ***= p < 0.001, ****= p < 0.0001. (See also Figure S1 & S2).

To assess whether glia were required for maintenance of axons in this model, we selectively ablated WG and measured neuronal integrity as the animals aged. We used the Gal4/UAS binary expression system (Brand & Dormand, 1995) to overexpress the cell death molecules – Dronc & Dark, or Reaper (Dorstyn et al., 1999; White et al., 1994; Zhou et al., 1999) – and tdTomato in a subset of the WG by using a split Gal4 construct (Luan et al., 2006). This split Gal4 was exclusively expressed in WG and is henceforth referred to as *WG split-Gal4*. *WG split-Gal4* labeled 87% of WG in the wing as determined by nuclear reporter expression as compared to *nrv2*-Gal4, which is expressed in all the WG in the wing but is also widely expressed in the central nervous system (Figure S1; (Neukomm et al., 2014). We combined the *WG split-Gal4* with the independent binary expression system QF2/QUAS (Potter et al., 2010; Riabinina et al., 2015) to fluorescently label a subset of *VGlut*^+^ neurons in the wing (∼40) in order to evaluate the effect of ablating WG on neurons in this nerve (Fig. 1D-F). Interestingly, ablating WG in these animals caused increased age-dependent degeneration of axons within this peripheral sensory nerve in the wing (Fig. D-G). A larger proportion of nerves from aged, ablated animals exhibited mild or severe degeneration compared with the control group (control: 2/18 animals or 11%, Dronc::GFP, Dark: 11/18 or 61%, Reaper: 10/18 or 56%; Fig. 1G & Figure S2A).

In addition to the effect that ablating WG had on axons, to our surprise, ablating WG also caused cell death of sensory neurons (Fig. 1H & Figure S2B). At baseline (4 days post eclosion) there were no significant differences in the average number of intact GFP^+^ neuron cell bodies per wing between genotypes (control: 38.0±2.1 n=21, Dronc::GFP, Dark: 36.9±2.7 n=20, Reaper: 37.2±2.4 n=20; Fig. 1H). However, at 28 days of age, cell death had increased significantly in WG-ablated animals compared to controls (control: 36.7±3.1 n=18, Dronc::GFP, Dark: 31.7±8.3 n=18, Reaper: 29.0±8.1 n=18; Fig. 1H). Together, these data indicate that WG are required for axon and neuronal maintenance in the sensory nerve of the wing.

### Identification of glial genes required for axon maintenance

Our results demonstrated that WG are required for axon maintenance in the L1 nerve, and we sought to use this system to identify glial genes required for supporting axon survival. We accomplished this by severing the axon from its soma thereby creating a situation where all that remains are the axons and the glia that ensheathe them (Fig. 2A). However, when wild type axons are severed, the portion of the axon distal to the injury site undergoes Wallerian degeneration (WD) (MacDonald et al., 2006). This can be genetically blocked by overexpressing Wld^S^ in neurons, which suppresses WD and allows severed axons to remain intact for weeks after axotomy (Fig. 2A; (Glass & Griffin, 1991). The ability of Wld^S^ to protect axons *in vivo* in intact nerves far exceeds its ability to do so in purified neurons *in vitro* (Adalbert et al., 2005; Buckmaster et al., 1995; Conforti et al., 2006; Lunn et al., 1989; Wang et al., 2005). We hypothesized that this greater protection was due to glial support of axon survival *in vivo* in intact nerves. Consistent with this notion, we found that when we ablated WG, Wld^S^-expressing axons exhibited increased axon degeneration compared to controls (degeneration phenotype: control=7/24 (29%), WG-ablated=18/21 (86%); Fig. 2C).

**Fig. 2:**
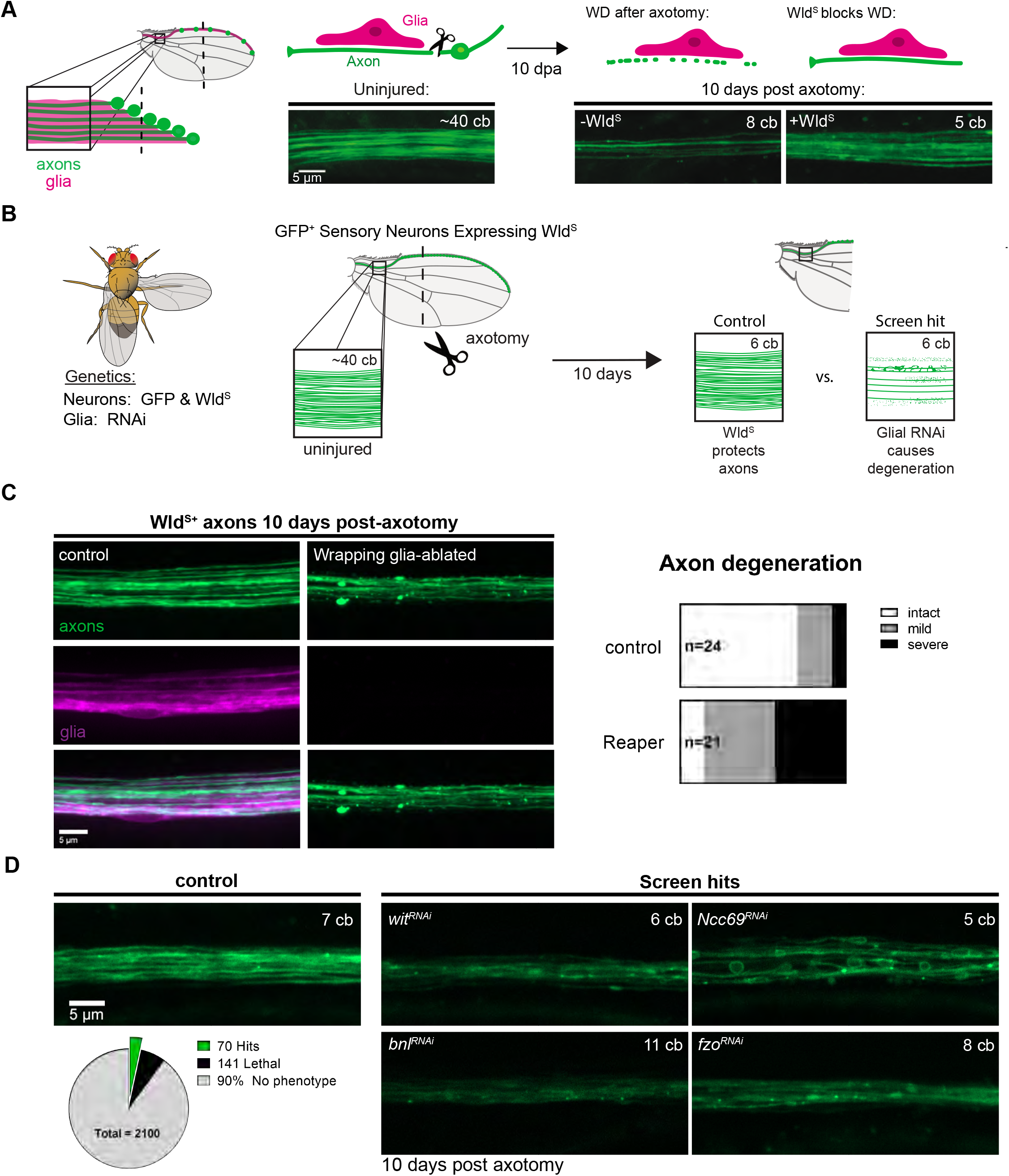
A screen for glial genes required for axon maintenance. A) Diagram illustrating the peripheral sensory nerve in the wing (left) with neurons labeled in green and glia in magenta. Expression of Wld^S^ prevents axon degeneration 10 days post axotomy (dpa) in the wing (right). Images are maximum intensity projections of z-stack images. B) Diagram of screen workflow. Flies expressing GFP and Wld^S^ in neurons and GAL4 in glia were crossed to UAS-RNAi flies and resulting progeny were injured and assessed for axon integrity at 10-14 dpa. C) Axon integrity in injured Wld^S^ expressing neurons in control and WG-ablated animals at 10 dpa. D) Summary of screen results with examples of individual screen hits: *wit*, *Ncc69*, *bnl* and *fzo*. Images are of single slices from middle of the axon bundle of z-stack images. (See also **<u>REF-Sup. file</u>**)

To identify glial genes that promote long term axon survival, we designed a genetic screen where we could sever Wld^S+^ axons, which allowed axons to survive for weeks after axotomy (and without a cell body), then systematically knock down genes selectively in glia and assay for axon loss compared to controls. Using *VGlut-QF2* (Diao et al., 2015), we expressed a membrane-tethered GFP and Wld^S^ in glutamatergic neurons in the L1 nerve of the wing to both visualize axons and block Wallerian degeneration (WD), respectively (Fig. 2B). In these animals, we removed the cell bodies from most of the neurons by cutting off the distal portion of the wing, leaving behind the WD-resistant axons and glia that surround them (Fig. 2A). We used *GAL4/UAS* to express RNA interference (RNAi) (Perrimon et al., 2010) constructs in all glia with the pan-glial driver *repo-Gal4* (Sepp et al., 2001). We then screened >2,000 publicly available *UAS-RNAi* lines targeting a panel of genes enriched for those encoding proteins containing predicted transmembrane domains or signal peptides (Fig. 2B).

In control animals, Wld^S^ prevented WD and axons remained intact at 10 days post axotomy (dpa) (Fig. 2C). In our screen, we sought to identify genes that, when knocked down in glia, resulted in axon degeneration (Fig. 2B). When we knocked down single genes in glia using this assay, we identified 70 candidate genes whose loss in glia resulted in axon degeneration or defects in axon morphology. For instance, depletion of a TGFβ receptor (*wit*), a fibroblast growth factor (FGF) (*bnl*), or a mitofusin (*fzo*) led to robust axon loss (Fig. 2D). Glial loss of the sodium-chloride co-transporter *Ncc69*, led to an axon blebbing phenotype similar to the neuronal activity-dependent axon disruption observed in zebrafish *slc12a2b* (NKCC1b) mutants (Marshall-Phelps et al., 2020). In addition, and consistent with previous work (Mukherjee et al., 2020), we also identified 141 genes that caused lethality—defined by absence of viable adult progeny—when selectively knocked down in glia (Fig. 2D).

### Knocking down glial TGFβ superfamily genes results in degeneration of severed Wld^S^-expressing axons

The TGFβ receptor *wit* was one of several members of the TGFβ superfamily identified in our screen. The TGFβ superfamily is made up of two major branches (TGFβ and BMP)(Upadhyay et al., 2017). Our initial screening panel did not include RNAis targeting all members of the TGFβ superfamily. We therefore obtained and tested additional RNAi lines to test all genes in this pathway using our sensitized screening approach (Fig. 3B). We found that most RNAi constructs targeting components of this superfamily caused axon degeneration or lethality when expressed in glia (Fig. 3B & C), suggesting a role for TGFβ signaling in glial support of axons. Surprisingly, knocking down of both ligands and their receptors—selectively in glia—caused axon defects, suggesting a potential glial autocrine signaling mechanism.

**Fig. 3:**
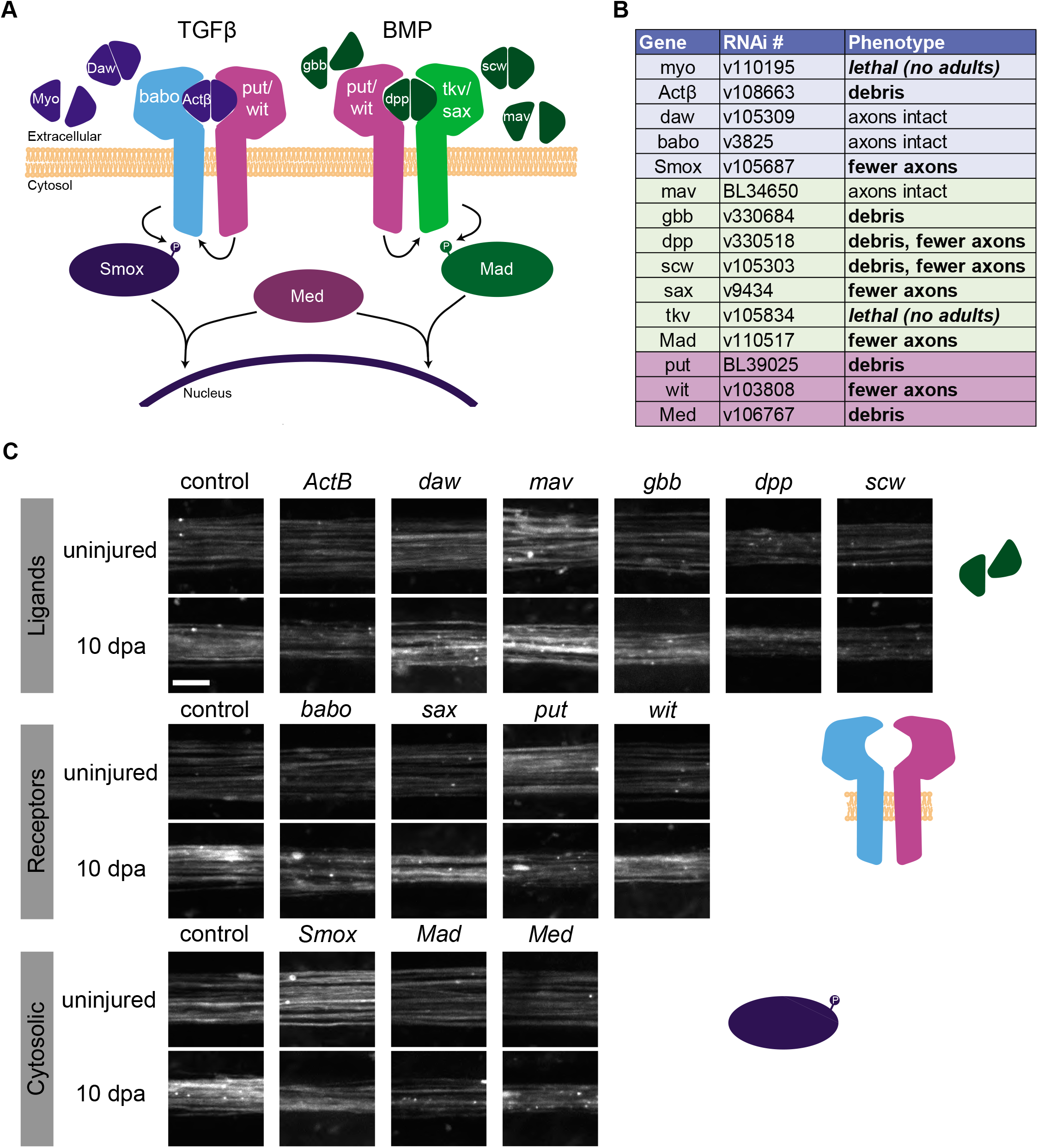
Glial-specific knockdown of TGFβ and BMP pathway components causes axon degeneration in the sensory nerve of the wing. A) Diagram of the TGFβ superfamily members in the *Drosophila* genome. B) Table summarizing the phenotypes for the corresponding RNAis targeting the TGFβ superfamily genes. VDRC - ‘v#’, Bloominton - ‘BL#’. C) Images of Wld^S+^ axons in the sensory nerve of the wing 10 days post-axotomy from control and TGFβ knockdown animals uninjured wing (top) axotomized wing (bottom). Scale bar 5 μm. Pan-glial knockdown of the ligand *myo* or the receptor *tkv* were lethal (not shown).

### TGFβ receptor *babo* is expressed in adult WG in the wing

To examine whether TGFβ proteins were expressed in nerve glia, we utilized a transgenic fly line where the *Gal4* sequence was inserted into the *babo* gene (Lee et al., 2018). We examined co-localization of a nuclear reporter (*UAS-lamin::GFP*) driven by *babo-Gal4* expression with an antibody that specifically labels WG nuclei within the larval peripheral nerves (Oaz) as well as the pan-glial protein Repo (Figure S3). All Oaz^+^ nuclei within larval nerves were GFP^+^/Repo^+^ (n=18 Oaz^+^ nuclei from n=3 larvae, Figure S3) indicating that *babo* was expressed in WG in peripheral nerves during development. Additionally, all Repo^+^/Oaz^-^ nuclei were also GFP^+^, indicating that *babo* was also expressed in other nerve glia as well as WG (n=123 Repo^+^/Oaz^-^ nuclei form n=3 larvae, Figure S3). To test whether *babo* was also expressed in adults, we crossed the *babo-Gal4* animals to a nuclear reporter (*UAS-mCherry.NLS*) in a genetic background where WG were labeled independently with GFP (a *nrv2-GFP* protein trap which labels all WG membranes (Stork et al., 2008)). As expected, our positive control (*nrv2-Gal4*) had nuclear reporter expression in WG nuclei within the GFP^+^ WG membrane at all timepoints tested (n=57 wings, Fig. 4). The experimental *babo-Gal4* wings also labeled nuclei within the WG membrane at 4 (n=20/20 wings, Fig. 4A), 14 (n=15/15 wings, Fig. 4B), and 28 (n=17/17 wings, Fig. 4C) days of age (and Figure S4). Importantly, the only nuclei present within the nerve in this region are glial nuclei (Neukomm et al., 2014), indicating that *babo* is expressed in mature WG within the peripheral sensory nerve in the wing. The *babo-Gal4* also appeared to label a subset of neurons within the wing at all timepoints tested (Figure S5). The expression of *babo* in adult WG suggests that the Babo receptor is present in adult glia and therefore could contribute to axon maintenance.

**Fig. 4:**
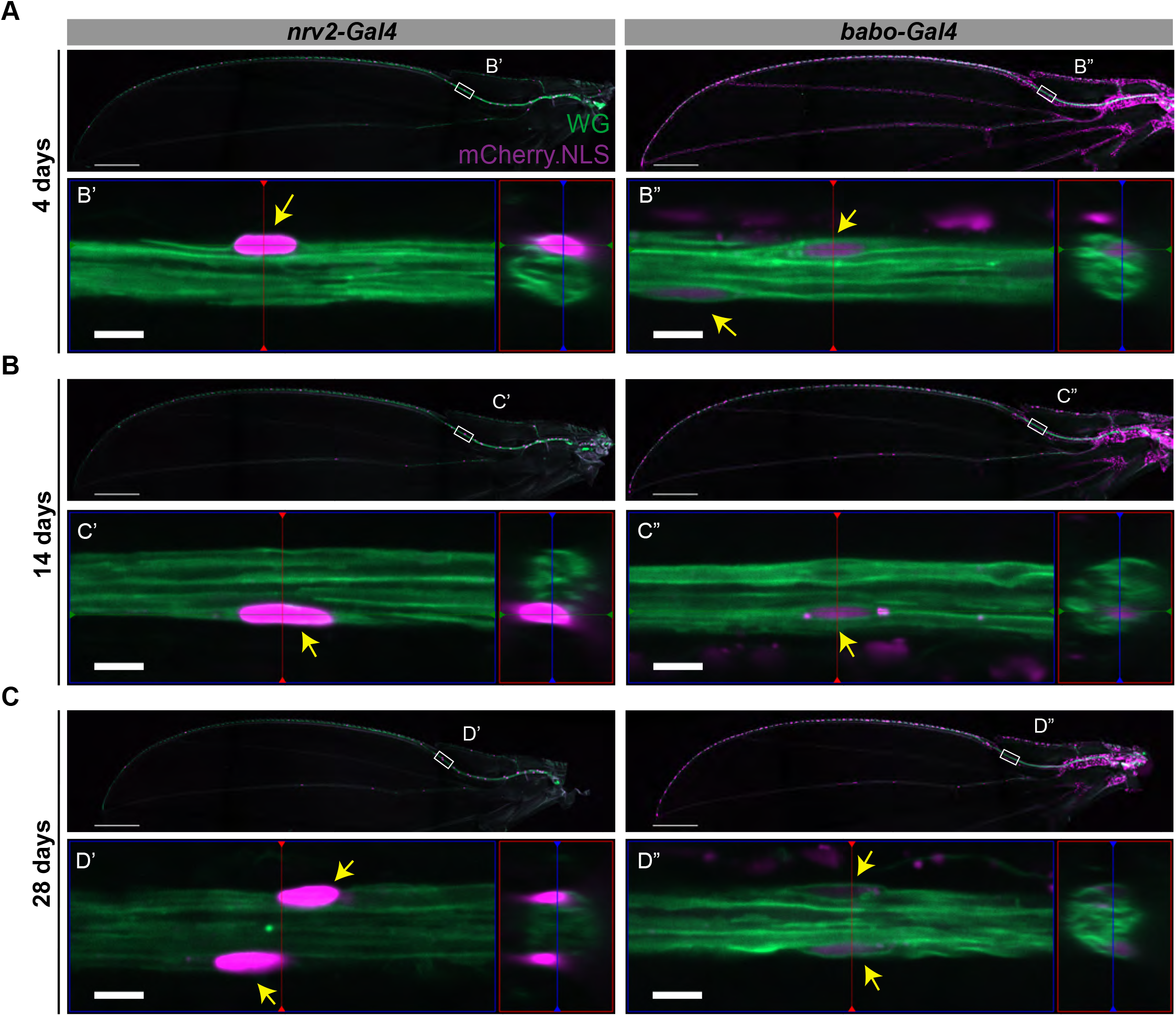
TGFβ receptor *babo* is expressed in adult WG. A) Nuclear mCherry reporter expression under *nrv2* (’) or *babo* (”) transcriptional control in a background where membrane-tethered GFP is expressed in WG at 4 (A), 14 (B), and 28 (C) days. Scale bar 200 μm. Higher magnification image of regions of interest in A-C showing mCherry^+^ nuclei within the GFP^+^ WG membrane (yellow arrows). Scale bar 5 μm. (See also Figure S3–S5)

### Loss of TGFβ does not alter ensheathment of axons

TGFβ molecules play crucial roles in tissue development and morphogenesis, so we next examined whether loss of this pathway would affect glial development and morphology. We evaluated the overall coverage of the L1 nerve by labeling the glia using a genetically encoded fluorescent reporter (tdTomato) and imaged the nerve at 4 and 28 days. When we examined nerves where TGFβ genes were knocked down in all glia, there were no obvious defects in glial morphology or coverage of the nerve in knockdown animals compared to controls at either timepoint (4 days: control: n=24 animals, *babo*: n=23, *Smox*: n=21; 28 days: control: n=21, *babo*: n=22, *Smox*: n=16) (Fig. 5A & Figure S6A). Even in nerves that had axonal debris present, glial ensheathment appeared normal (Fig. 5A). Since WG directly ensheathe axons in this nerve, we also assessed WG morphology specifically, by examining reporter expression in WG-specific knockdown conditions compared to controls. Again, we found no changes in morphology or coverage of the nerve by WG despite the presence of neurodegeneration (4 days: control: n=23 animals, *babo*: n=23, *Smox*: n=23; 28 days: control: n=22, *babo*: n=24, *Smox*: n=23) (Fig. 5B & Figure S6B).

**Fig. 5:**
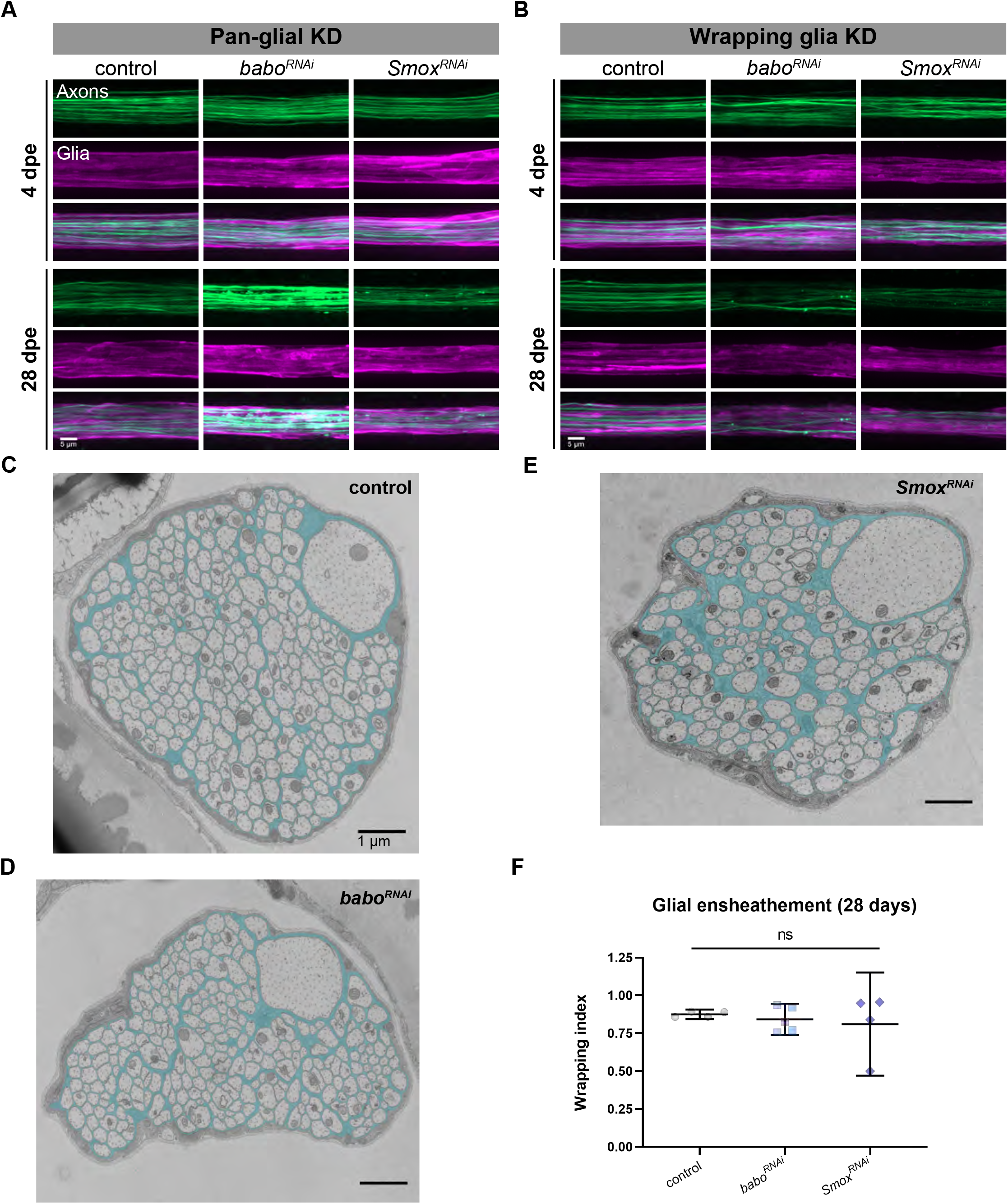
Glial morphology appears intact in TGFβ knockdown animals. A) Axons (green) and glia (magenta) in control and pan-glial TGFβ knockdown animals from 4 (top) and 28 (bottom) day old L1 nerves. B) Nerves from WG-specific TGFβ knockdown animals from 4 (top) and 28 (bottom) day old L1 nerves. C-E) Electron micrographs of a cross section of the L1 nerve at 28 days from control (C), *babo* (D), and *Smox* (E) knockdown animals. Wrapping glia are highlighted in cyan. F) Quantification of the wrapping index [(individually wrapped axons + bundles of axons) / total axons]. Data are represented as mean ± 95% CI. Statistics: One-way ANOVA with Dunnett’s multiple comparisons test. Significance: ns = p > 0.05. (See also Figure S6)

While these results suggested that gross morphology of WG were unaltered upon knockdown of TGFβ signaling, to examine whether axonal ensheathment was truly unperturbed required us to assess the ultrastructure of this nerve using transmission electron microscopy. We examined nerves from control, *babo,* and *Smox* pan-glial knockdown animals and found no evidence of defects in glial ensheathment at 28 days (control: n=3 animals, *babo*: n=5, *Smox*: n=4; Fig. 5C-E & Figure S6 C-E). To quantify this form of multi-axonal ensheathment, we measured the wrapping index for each nerve (Matzat et al., 2015). When we calculated the wrapping index for each condition, there was no significant difference between the conditions, although one nerve from the *Smox* knockdown condition did appear to have reduced ensheathment (control: 0.88±0.020 n=4, *babo*: 0.84±0.083 n=5, *Smox*: 0.81±0.21 n=4; Fig 5F). Together, our data indicate that inhibiting TGFβ in glia does not cause defects in glial morphology and therefore disrupted glial ensheathment would not explain the degeneration observed in TGFβ knockdown animals.

Although WG ensheathment was not disrupted, it remained possible that the number of WG could be impacted by inhibiting TGFβ. We quantified the number of WG present in the nerve in control and knockdown animals by using a genetically encoded nuclear reporter (*UAS-lamin::GFP*) (Figure S7). Knock down of *babo* or *Smox* in WG caused a slight increase in the number of WG in the nerve at 4 days (*nrv2*>: 24.9±3.52, *nrv2>lacZ*: 26.9±3.37, *nrv2>babo^RNAi^*: 30.1±3.54, *nrv2>Smox^RNAi^*: 29.0±3.76; Figure S7). At 28 days, the number of WG nuclei in *babo* knockdown animals remained elevated compared to controls, but the number of WG nuclei in *Smox* knockdown animals was not significantly different from controls (*nrv2*>: 24.5±3.40, *nrv2>lacZ*: 26.2±3.39, *nrv2>babo^RNAi^*: 29.5±3.38, *nrv2>Smox^RNAi^*: 25.4±6.76; Figure S7).

In summary, while we find a slight increase in the total number of WG in the adult nerve when TGFβ signaling is inhibited, overall glial morphology and axonal ensheathment appeared normal.

### TGFβ superfamily is required in WG for long-term neuron maintenance

Our finding that ablating WG caused spontaneous, age-dependent degeneration supports a role for WG in suppressing neurodegeneration as animals age. The TGFβ signaling receptor Babo appears to be expressed for at least 28 days into adulthood, but is glial TGFβ signaling involved in this glia→neuron pro-survival mechanism? To explore this possibility, we targeted each of the TGFβ superfamily genes using RNAi and evaluated axon integrity and neuron survival during normal aging at three timepoints: 4, 14, and 28 days. Interestingly, we found robust age-dependent neurodegeneration in several TGFβ knockdown conditions compared to controls, with the strongest phenotype elicited by knock down of the TGFβ receptor *babo* (Fig. 6A-C & Figure S8). Additionally, we used an RNAi-independent method to inhibit TGFβ signaling by overexpressing a dominant negative form of the receptor (*babo^DN^*) in glia that lacked the kinase domain (Brummel et al., 1999). Glial expression of *babo^DN^* phenocopied the RNAi-mediated knockdown (Fig. 6D), indicating that disruption of Babo in glia is sufficient to induce neurodegeneration. We confirmed the specificity of the RNAi targeting *babo* and its downstream target *Smox* with additional non-overlapping RNAi constructs (Fig. 6E). From these data, we conclude that the TGFβ pathway is required in glia for long-term neuron survival.

**Fig. 6:**
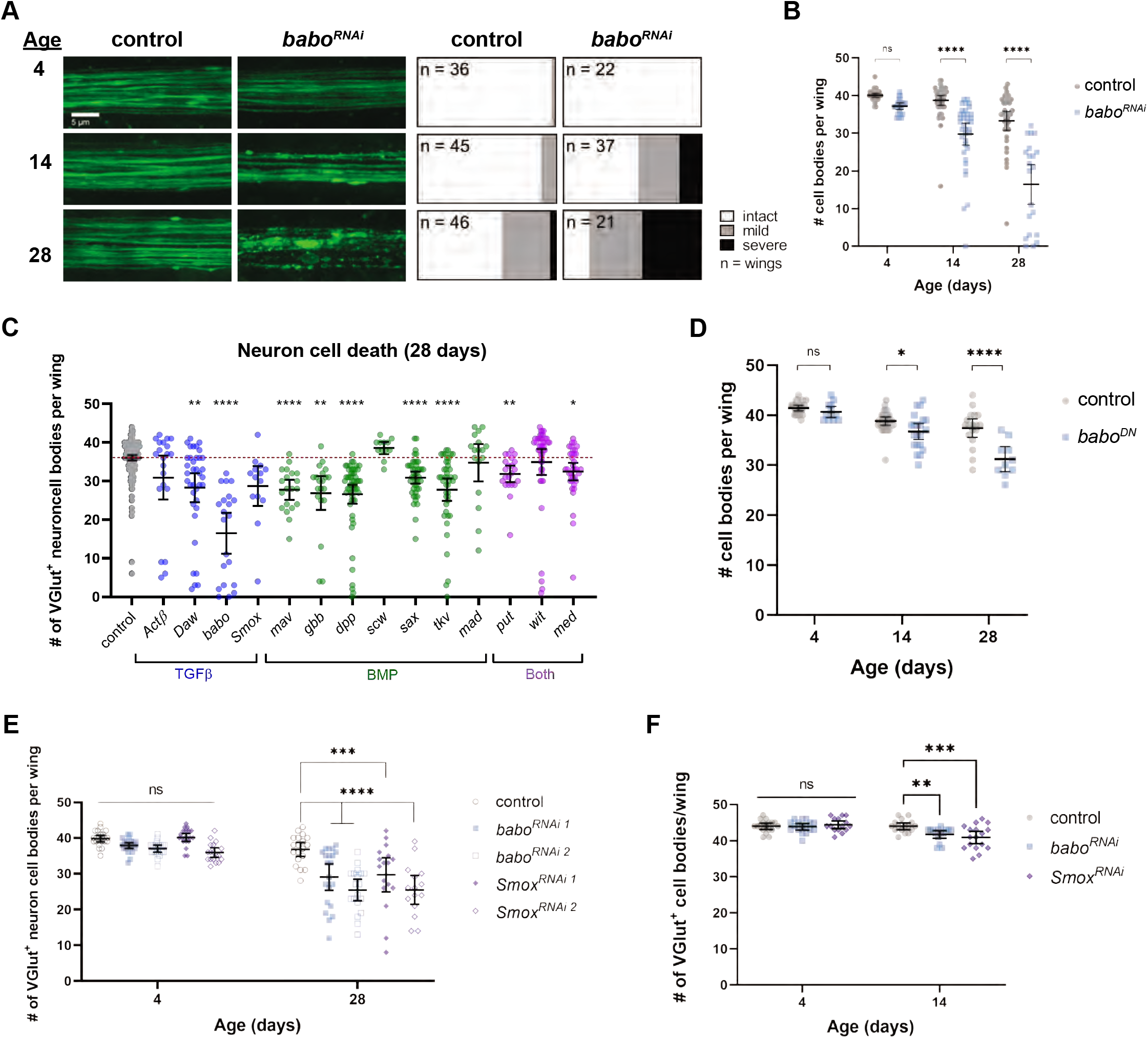
Glial-specific knockdown of TGFβ superfamily members results in age-dependent neurodegeneration in the sensory nerve in the adult wing. A) Images from control and *babo*-knockdown animals of the axon bundle in the wing at each time point (left) and the proportion of wings in each classification (right) for all conditions. B) Quantification of the number of intact neuron cell bodies per wing at each time point in control and *babo*-knockdown animals. C) Comparison of the number of intact neuron cell bodies per wing at 28 days for control and TGFβ superfamily knockdown conditions. Note: repeated data shown in B and C for control and *babo*-knockdown at 28 days. D) Quantification of the number of intact neuron cell bodies at each time point in control animals and animals with over-expression of a dominant negative form of the *babo* receptor (*babo^DN^*). E) Quantification of the number of intact neuron cell bodies in control, *babo*, and *Smox* knockdown animals using two non-overlapping RNAis each. F) Quantification of the number of intact neuron cell bodies in adult-specific knockdown animals. Data are represented as mean ± 95% CI. Statistics: (B, D, E, F) Two-way ANOVA with Sidak’s multiple comparisons test; (C) One-way ANOVA with Dunnett’s T3 multiple comparisons test. Significance: ns= p > 0.05, *= p < 0.05, **= p < 0.01, ***= p < 0.001, ****= p < 0.0001. (See also Figure S8)

It is possible that the effects we observe on neuronal survival could be due to developmental defects or represent a role for TGFβ signaling in adults. To determine whether there is a requirement for TGFβ signaling in adults, we combined glial-specific knockdown with a temperature-sensitive Gal80 construct to temporally control RNAi expression (McGuire et al., 2003). Using this tool, we inhibited RNAi expression during development by rearing animals at 18°C until they eclosed as adults. We then transferred adult animals to 31°C to allow glia-specific RNAi expression to knock down *babo* or *Smox*. At 4 days post temperature shift, there was no significant difference in the number of neuron cell bodies between control and TGFβ knockdown animals (control: 44.0±1.65, *babo*: 43.9±1.65, *Smox*: 44.4±1.88) (Fig. 6F). However, after 14 days of RNAi-mediated knockdown, both *babo* and *Smox* knockdown animals had fewer intact neuron cell bodies (control: 44.0±1.71, *babo*: 41.8±1.94, *Smox*: 40.9±3.31) (Fig. 6F). These data indicate that *babo* and *Smox* are required in glia in the mature nerve to promote neuron survival and suppress neurodegeneration.

### Neuronal Wld^S^ expression rescues age-dependent neurodegeneration induced by glial *babo* knockdown

In aging animals both the axons and cell body of the neurons were affected by *babo* knockdown in glia. This could result from a lack of glial support of neuronal cell bodies or axons. The neuroprotective effects of Wld^S^ are known to specifically mediate injury-induced axon degeneration while it fails to block apoptosis in several contexts (Beirowski et al., 2008; Deckwerth & Johnson, 1994; Hoopfer et al., 2006). We tested whether promoting axon survival with Wld^S^ could save axons, and if so, we could use this to test whether the cell body loss observed in *babo* glial-knockdown animals was dependent on axon loss. We overexpressed Wld^S^, in the glutamatergic neurons to prevent WD in aged control and *babo* knockdown animals and measured its effect on both axon and cell body integrity. Blocking WD rescued the axon degeneration observed in *babo* knockdown animals at 28 days (control-Wld^S^ n=17, control +Wld^S^ n=19, *babo^RNAi^* -Wld^S^ n=27, *babo^RNAi^* +Wld^S^ n=20) (Fig. 7A & B) indicating that loss of *babo* in glia leads to activation of a Wld^S^-sensitive axon degeneration pathway in axons. Moreover, we found that suppressing axon degeneration with Wld^S^ also completely rescued neuron cell death (Fig. 7A & C). These data are consistent with a similar finding that suppressing axon degeneration with Wld^S^ in a model of motoneuron disease reduced subsequent cell death of neurons (Ferri et al., 2003). Our findings suggest that glial loss of Babo leads to activation of an axon degeneration pathway in neurons that can ultimately result in neuronal cell death.

**Fig. 7:**
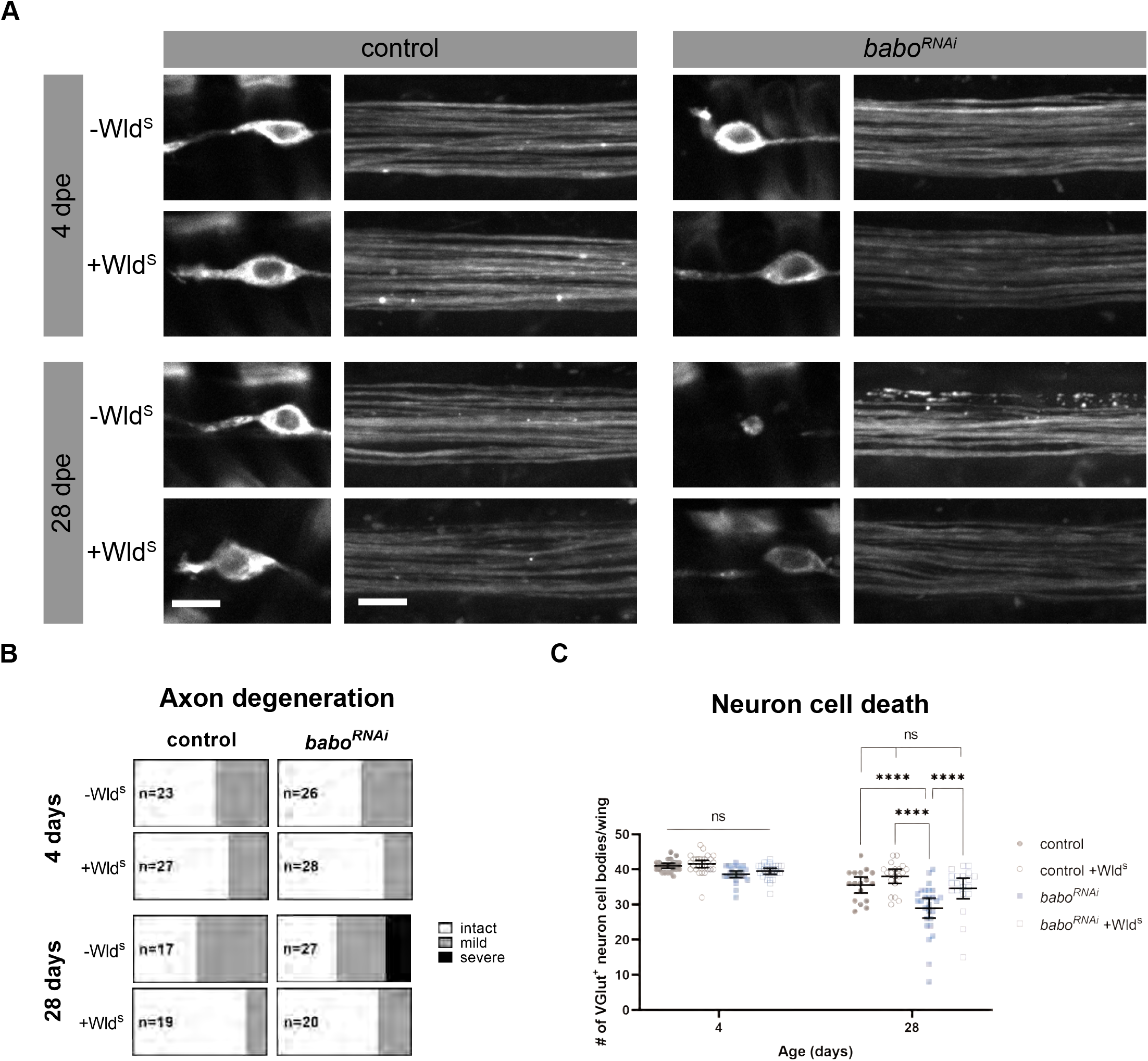
Wld^S^ overexpression in neurons rescued neurodegeneration in *babo* knockdown animals. A) Representative images of neuron cell bodies and axons within the wing in control and *babo* knockdown animals at 4 (top) and 28 (bottom) days of age with and without Wld^S^ expressed in neurons. Scale bar 5 μm. B) Classification of axon integrity in control and *babo* knockdown animals at 4 and 28 days with and without Wld^S^. C) Quantification of the number of VGlut^+^ neuron cell bodies in the wing from control and *babo* knockdown animals with and without Wld^S^. Data are represented as mean ± 95% CI. Statistics: 2-Way ANOVA with Tukey’s multiple comparisons test. Significance: ns= not significant, ****= p <0.0001.

## Discussion

In this study, we sought to identify genetic pathways that modulate glial support of axons *in vivo*. We focused on the L1 wing nerve of adult *Drosophila*, as these are among the longest axons in the fly, and the wrapping glia that ensheathe them separate axons at the single axon level. Through a genetic screen to identify genes with roles in promoting axon survival, we assayed a library consisting of most of the secreted and transmembrane proteins encoded in the *Drosophila* genome. We identified an array of molecules that, when depleted selectively from glia, lead to axon degeneration. Components of the TGFβ superfamily were over-represented in our candidates from this screen, and here we have shown that loss of glial TGFβ signaling— ligands, receptors, or downstream signaling molecules—leads to age-dependent axon degeneration and neuronal loss. Surprisingly, we found that providing Wld^S^ to neurons was sufficient to overcome neurodegeneration caused by reduced TGFβ signaling in glia. We propose that TGFβ signaling in glia normally promotes axon maintenance, and this support is required to sustain neuronal survival over the animal’s lifespan.

The intense ensheathment of axons by wrapping glia in the adult L1 wing nerve suggests that isolating individual axons in this tissue is critical for neuronal function, maintenance, or both. It is also worth noting that this ensheathment creates a physical barrier that likely prevents axons from directly accessing metabolites outside the nerve needed to support their activity. This anatomical arrangement suggests that glia act as the go-between and provide support to the axons they enwrap. Using a new, highly specific split Gal4 line, we showed that ablating most of the wrapping glia in the adult peripheral nerve in the wing caused robust age-dependent neurodegeneration. These results are remarkably similar to what has been seen in vertebrate models of CMT where loss of myelinating glia results in neurodegeneration in peripheral nerves (Adlkofer et al., 1995; Verhamme et al., 2011). Whether the loss of wrapping glia in the fly leads to axon degeneration because of a loss of metabolic support, as is observed in mammals (Fünfschilling et al., 2012; Y. Lee et al., 2012), or is a direct result of a lack of survival cues through TGFβ signaling remains unclear. Nevertheless, our study supports the notion that the L1 wing nerve of *Drosophila* should be an excellent model to explore how glia support axon maintenance and function.

One could imagine that disruptions in the physical association between axons and glia could indirectly disrupt signaling between them, for instance by impinging on the ability of glia to provide metabolites or survival cues to axons. Conversely, reduced ensheathment could presumably impair the ability of glia to maintain proper extracellular ion concentrations causing disruptions to axon integrity like is seen in our *Ncc69^RNAi^* condition (Figure 1; Leiserson et al., 2011) and the homologous NKCC1b mutant zebrafish (Marshall-Phelps et al., 2020). However, reduced glial ensheathment does not appear to explain the phenotypes reported here in *babo* and *Smox* knockdown animals. Our live imaging and ultrastructural analyses both indicate that glia remain tightly associated with axons in these knockdown animals. Additionally, knockdown of either *babo* or *Smox* did not cause a decrease in WG numbers, but rather an increase. This result is consistent with previous literature on the role of TGFβ in developmental apoptosis (D’Antonio et al., 2006; Parkinson et al., 2001). Indeed, inhibiting apoptosis in WG phenocopied the increase in WG numbers observed in *babo^RNAi^* indicating that this phenotype was likely caused by TGFβ’s role in developmental apoptosis. However, we do not believe that the increase in WG is responsible for the age-dependent neurodegeneration present in *babo^RNAi^* conditions since adult-specific knockdown was sufficient to induce neurodegeneration. We propose that our data point instead to a direct signaling role for glial TGFβ in promoting neuron maintenance.

The precise role of the TGFβ pathway in supporting axons remains unclear. Inhibiting members of both branches of the TGFβ superfamily (TGFβ and BMP) in glia caused neurodegeneration in aged animals. The TGFβ receptor Babo phosphorylates and activates the Smad transcription factor Smox, allowing it to enter the nucleus (Upadhyay et al., 2017). Smox has the potential to modify many genetic pathways simultaneously, and given that loss of Smox phenocopies Babo depletion, our data argue that at least part of the ability of TGFβ signaling to support axon maintenance involves gene regulation. Identifying the key transcriptional targets for Smox that promote this pro-survival function in axons will be an exciting next step in understanding the full array of molecules involved in glial support of axon maintenance.

Is TGFβ signaling required during development or in the adult to support axon survival? Using a genetically encoded reporter, we found that *babo* (encoding the TGFβ receptor) is expressed in glia in the adult wing, supporting a role for this receptor in mature glia. We further showed that mature glia utilize the TGFβ pathway to promote axon maintenance, as conditional knockdown of *babo* or *Smox* specifically in mature glia was sufficient to induce neurodegeneration. The adult-specific knockdown was modest compared to the constitutive knockdown conditions, and this could indicate that TGFβ signaling is required both during development and in the mature nerve. However, there were crucial differences between these experimental approaches that make direct comparison challenging. Nevertheless, together our data support a role for the TGFβ pathway in mature tissues to promote axon survival.

The role of the axon in survival of the cell body appears complex, and context dependent. For instance, during development, a lack of trophic support in growing axons leads to death of dorsal root ganglion (DRG) cell bodies (Deckwerth & Johnson, 1994). This allows sensory ganglia to scale DRG numbers according to segment specific changes in the size of the sensory field. Once mature, however, the majority of sensory and peripheral motor axons exhibit robust regenerative capacity when axotomized *in vivo* (Cajal et al., 1991) indicating the requirement for trophic molecules from distal axons in promoting cell body survival is transient. In the context of most neurodegenerative diseases, it remains unclear which part of the cell becomes sick first. Does the axon degenerate, fail to support the neuronal soma, and results in cell death? Alternatively, does the cell body die and axon degeneration follows as a result? One can explore this through examining the timing of these events. Dying back neuropathies, where axons are lost first, suggest that axon loss may drive neuronal death (Fischer et al., 2004; Monte et al., 1988; Schaumburg et al., 1974; Sima et al., 1983). However, neuronal drop out in many neurodegenerative diseases is often sparse and occurs over a protracted phase and high-resolution imaging is not possible in human patients. Unless one can assay both parts of the very same neuron, it is impossible to know which came first. Our system in the L1 wing nerve allows us to assay these events with single cell precision in many cases. That said, we found that axon degeneration and neuronal cell loss was coordinately timed in both our glial-ablated and TGFβ knockdown animals. As such, we are unable to determine which compartment of the cell (i.e., axons versus cell body) began to degenerate first.

An alternative approach to determine where neurodegenerative mechanisms impinge on a healthy neuron (axon versus soma) is to block key genetic pathways that drive axonal degeneration versus cell death. Driving expression of the Wld^S^ molecule in neurons is an effective way to suppress axon degeneration without altering apoptotic signaling pathways (Adalbert et al., 2006; Beirowski et al., 2008; Deckwerth & Johnson, 1994) and can be used to probe whether neurodegeneration is driven by primarily the axon versus the cell body. In the case of the *progressive motorneuronopathy* (*pmn*) mouse, which exhibits age dependent motor axon degeneration followed by motoneuron cell death, simply supplying Wld^S^ was sufficient to block axon loss and suppress motoneuron death (Ferri et al., 2003). These data strongly suggested that axon loss was the primary driver of motoneuron loss in this model. Since this initial exciting discovery similar approaches have been used with Wld^S^ and other components of the axon death signaling cascade like Sarm1 (Osterloh et al., 2012), to probe the molecular programs driving neurodegeneration in a number of disease models (reviewed in (Coleman & Höke, 2020)). In some cases, blocking axon degeneration can profoundly block neurodegeneration (Ferri et al., 2003; Henninger et al., 2016; Samsam et al., 2003; Turkiew et al., 2017) while in others, it has surprisingly little effect (Adalbert et al., 2006; Beirowski et al., 2008; Fernandes et al., 2018; Fischer et al., 2005; Peters et al., 2018). The latter observation does not rule out a role for Wld^S^ or Sarm1 in these diseases, it only reveals that blockade of the axon death pathway alone is not sufficient to alleviate neurodegeneration in these models of human disease.

Our findings that preventing Wallerian degeneration of uninjured, normally aging axons in animals lacking glial TGFβ signaling rescues neurodegeneration argues that loss of glial TGFβ signaling activates a Wld^S^-sensitive axon death signaling cascade to drive axon loss and ultimately cell body death. It is notable that Wld^S^ was able to rescue age-dependent degeneration in TGFβ knockdown conditions, but not in our sensitized injury paradigm where TGFβ genes were knocked down in glia and axons were severed. One could imagine that an aged axon that still has a cell body is in a better position than an axon that has been severed and no longer has support from its own cell body. Therefore, the difference in these two sensitizing conditions likely explain the difference in the ability (or lack thereof) of Wld^S^ to rescue axon degeneration when TGFβ genes are also knocked down. Furthermore, the conditions where Wld^S^ was unable to rescue injury-induced degeneration when TGFβ genes were knocked down in glia, implicate this pathway in glial support of axon maintenance. We would speculate that loss of *dsarm* in axons would result in a similar phenotype (i.e., rescue of axons), but technical limitations precluded our doing this experiment. While it remains possible that Wld^S^ directly protects the neuron cell bodies, this would depart from the body of data supporting an axon-specific role for Wld^S^ in suppressing neurodegeneration. Therefore, the simplest interpretation of our data is that TGFβ signaling activity is required in glia to support axon maintenance and its loss results in progressive axon degeneration culminating in neuron cell death.

A yet-to-be explained phenomenon in the study of axon degeneration is the observation that Wld^S^ is a much more potent suppressor of Wallerian degeneration *in vivo* as compared to *in vitro*. Severed Wld^S+^ axons survive for many weeks *in vivo* (Adalbert et al., 2005; Lunn et al., 1989), yet in purified neuron cultures, protection persists for several days (Buckmaster et al., 1995; Wang et al., 2005). While there are many differences between these environments, one striking difference is the presence or absence of a glial sheath around axons. Our findings that Wld^S^ protection of injured axons *in vivo* is significantly impaired when glia are absent argues that when axons lack a neuronal cell body their survival is highly dependent upon glial support.

In summary, our work provides direct *in vivo* evidence that glia are crucial for supporting long axons. When these glia are eliminated, axons show increased age-dependent degeneration and even Wld^S^ is unable to protect axons without glial support. We have identified several candidate glial genes required for this glial support of axon integrity and neuronal survival *in vivo*. Among these genes, our data show that the TGFβ signaling pathway plays a crucial role in suppressing axon degeneration, even in uninjured nerves, and its loss leads to activation of the axon death program that is sensitive to Wld^S^.

## Materials and methods

### Fly husbandry

Flies were grown on standard molasses cornmeal agar with extra yeast and maintained at 25°C. The following fly (*Drosophila melanogaster*) stocks used in this study were obtained from the following sources. Bloomington: *OK371-QF2* (66473), *10xQUAS-6xGFP*, *UAS-mtdTomato-3xHA* (66479), *Repo-GAL4* (7415), *UAS*-*Reaper*^14^ (5824), *UAS*-*dronc::GFP* (56759), *UAS-lacZ.NZ^312^* (3956), *UAS-lacZ.NZ^20b^* (3955), *VGlut-QF2* (60315), *QUAS*-*mCD8::GFP* (30002), *UAS-lamin::GFP* (BL7376), *nrv2-GAL4* (6799), *UAS*-*babo^DN^* (64423), *babo^RNAi2^* (40866), *Smox^RNAi2^* (41670), *tkv^RNAi^* (40937), *mav^RNAi^* (34650), *put^RNAi^* (39025), *UAS-mCherry.NLS^3^* (38424), *babo*-*Gal4*^CRIMIC00274^ (83164), *tubP-Gal80^ts-20^* (7019). Vienna *Drosophila* Resource Center RNAi lines are listed in the supplementary excel file (<u>Supp File 1</u>). Additional RNAi lines including *tkv^RNAi^, put^RNAi^,* and *sax^RNAi^* were generously provided by Dr. Michael O’Connor. *UAS*-*dark* was kindly provided by Dr. John M. Abrams (Akdemir et al., 2006). The protein trap *nrv2-GFP* published in Stork et al., 2008. The *WG split-Gal4* line was established using the *nrv2*-DNA binding domain construct previously reported in (Coutinho-Budd et al., 2017) combined with a VP16 activation domain converted from the *IT.0117-Gal4* (BL62647) using methods described in (Gohl et al., 2011). Additional lines that we generated to complete this work were *QUAS*-*Wld^s^*(III).

### Sensitized RNAi screen

RNAi lines were crossed to the *w*; VGlut-QF2, QUAS-mCD8::GFP/*CyO *; QUAS-Wld^s^, Repo-Gal4/TM3* driver line. After 7 days, parents were discarded and progeny returned to 25°C. Progeny were then anesthetized on CO_2_ fly pads, sorted for genotype using visible markers, aged 4 days at 25°C, anesthetized on CO_2_ and one wing was cut between the two cross veins of the wing using spring scissors (F.S.T #15002-08), while the other wing served as an uninjured control. Injured flies were transferred to fresh vials every 3-7 days and then imaged 10 or 14-days post axotomy (see imaging). For each RNAi line at least 5 wings were evaluated, results are reported in the supplemental excel file (<u>Supp File 1</u>). RNAi lines were scored as lethal if no viable adult flies of the correct genotype emerged or if adults died before the imaging timepoint. Both female and male progeny were used except where genetics prohibited use of males.

### Aging assay

Animals of the appropriate genotypes were crossed, as described above, selected for markers at eclosion, and adults were aged for the indicated time windows at 25°C. Aging flies were transferred into fresh vials every 3-7 days. The number of dead flies in each vial was recorded during each transfer and these tallies can be found in Supplementary Fig. 6. Subsets of wings from each cohort were imaged at 4, 14, and 28 days after progeny were originally collected. All wings were inspected at 63x for injuries and were not evaluated if they had any visible tears or scars in the L1 wing vein containing the nerve.

### Adult-specific knockdown

Crosses were performed at 18°C and the progeny were allowed to develop at 18°C. Adults of the correct genotype were collected into fresh cornmeal agar vials and transferred to 31°C. Flies were maintained at 31°C and transferred to fresh vials every 3-5 days until imaging.

### Imaging

Imaging of the wing nerve was done as previously described in (Neukomm et al., 2014). Briefly, flies were anesthetized using CO_2_ and their wings were removed using spring scissors, mounted on a slide in Halocarbon oil 27 (Sigma #H8773), covered with #1.5 cover glass, and imaged within 15 minutes of mounting. Z-stack images were taken of the nerve on a Zeiss Axio Examiner with a Yokogawa spinning disk and Hamamatsu camera using a 63×1.4NA oil-immersion objective. The same acquisition settings were used across samples for each of the experiments and control samples were imaged in the same imaging session as experimental samples. *VGlut*^+^ neuron cell bodies in the L1 vein were counted under 63x magnification. Cells were counted as intact if they had a clear nucleus and dendrite or were considered dead if they were shrunken and the dendrite or nucleus were not clearly visible (Supplemental Fig. 2B).

### Quantification of axon degeneration

Images were classified into phenotypic categories (intact, mild, or severe degeneration) with the conditions blinded to the scorer (Figure S2A). Images were given randomized numerical names and all genotypes and ages for a given experiment were scored together in one session and later decoded. For experiments in which the wrapping glia were ablated, the channel containing the axons was first extracted from the two-color images before blinding and scoring so that the scorer remained blind to the presence or absence of glia.

### Immunofluorescence

Wandering third instar larvae were dissected and pinned open as filets in cold PBS and fixed in 4% paraformaldehyde in PBS for 15 minutes at room temperature. Larvae were then permeabilized in 0.3% PBST (PBS + TritonX-100) for 15 minutes at room temperature with agitation and remaining wash and antibody solutions were made in 0.3% PBST. Antibodies used were: (1°) anti-Repo (Mouse anti-Repo, DSHB #8D12), Alexa Fluor® 647 anti-HRP (Goat anti-HRP, Jackson Labs #123-605-021), anti-oaz (Rabbit anti-oaz, this paper), anti-GFP (Chicken anti-GFP abcam #ab13970); (2°) DyLight™ 405 Donkey anti-Mouse (Jackson Labs #715-475-150), Alexa Fluor® 488 Donkey anti-Chicken (Jackson Labs #703-545-155), Rhodamine Red™-X Donkey anti-Rabbit (Jackson Labs #711-295-152). After staining, larva filets were mounted in Vectashield (Vector Labs #H-1000) and covered with #1.5 cover glass (Globe scientific #1404-15) and stored at 4°C.

### Electron microscopy

Aged flies were maintained as described above. For EM procedures we used a modified microwave protocol from (Cunningham & Monk, 2018; Czopka & Lyons, 2011). Flies were anesthetized with CO_2_ and their wings were removed with spring scissors and immediately put into freshly made fix solution (2% glutaraldehyde, 4% paraformaldehyde, 0.1M sodium cacodylate buffer). Forceps were used to gently submerge the tissue in a microcentrifuge tube and microwaved using the following settings: 2x (100W for 1min, OFF for 1 min), then immediately followed by 5x (450W for 20s, OFF for 20s) before storing the tissue at 4°C overnight in fix solution. The following day samples were washed in 0.1M sodium cacodylate buffer followed by secondary fixation in 2% osmium tetroxide, 0.1M sodium cacodylate buffer and 0.1M imidazole pH 7.5 and microwaved 2x (100W for 1min, OFF for 1 min), 5x (450W for 20s, OFF for 20s). Following osmium fixation, samples were rinsed in distilled water 3 x 10-minute washes. Next, samples were stained in saturated uranyl acetate (UA) ∼8% in water and microwaved 2x (450W for 1 min, OFF for 1 min). This was followed by dehydration steps with an escalating ethanol series with each step microwaved at 250W for 45s. The final 100% EtOH step was repeated 3 times and each step was microwaved for 2x (250W for 1 min, OFF for 1 min). Following EtOH dehydration, samples were dehydrated in 100% acetone and microwaved 2x (250W for 1 min, OFF for 1 min) and repeated 3 times. Next, samples were transferred to a 50:50 resin:acetone solution and agitated overnight at room temperature. Final resin infiltration was done in 100% resin and agitated at room temperature for at least 1 hour. Tissues were embedded in Embed 812 resin (EMS #14120) and cured in a 60°C oven overnight. Ultrathin 70 nm sections were cut on a Leica ultramicrotome and transferred to 100mesh Formvar grids (EMS #FCF100-Cu). Grids were counter stained for 20 minutes in 5% uranyl acetate followed by 7 minutes in Reynold’s lead citrate. Micrographs were acquired on a FEI Tecnai T12 interfaced to Advanced Microscopy Techniques (AMT) CCD camera.

### Statistical analysis

Statistical analyses were done in GraphPad Prism 8. When analyzing the effect of two variables (genotype and age) two-way ANOVA was used with Sidak’s multiple comparisons test to analyze the effect of genotype at each age. When comparing multiple experimental groups to the same control group Welch’s ANOVA was used with Dunnett’s T3 multiple comparison test to compare experimental groups to the control. When comparing one experimental group to a control a one-tailed Welch’s t test was used. Significance was determined using an α of 0.05. In figures, p-values are represented as follows: * p<0.05, ** p<0.01, *** p<0.001, **** p<0.0001.

## Acknowledgements

We thank all the Freeman Lab members for their discussion and feedback. We would like to acknowledge the technical support and expertise of the staff in the electron microscopy core facility, particularly Dr. Robert Kayton as well as Dr. Deborah Hegarty from the Aicher Lab. We would also like to acknowledge Dr. Kelly Monk and members of the Monk Lab for sharing their expertise and equipment in regards to the electron microscopy experiments. We thank Dr. Kevin Wright and Dr. Ben Emery for their critical feedback throughout the development of this work. We also acknowledge Dr. Rachel Dresbeck critical reading of the manuscript. The study was supported by funding from NIH grant from NINDS P30 NS061800 (S.A.A.) and NIH RO1 grants NS059991 and NS112215 (M.R.F.) and OHSU.

## Author Contributions

Conceptualization, A.P.L and M.R.F.; Methodology A.P.L, M.M.C., J.H., A.N.F. and M.R.F.; Investigation, A.P.L., M.M.C., R.B., A.E.S., J.H., A.N.F.; Resources, S.A.A. and M.R.F., Writing – Original Draft, A.P.L. and M.R.F. Writing – Review & Editing, A.P.L, M.M.C., S.A.A. and M.R.F.; Supervision, M.R.F.; Funding Acquisition, S.A.A and M.R.F.

---

**Supplemental Figure 1:**
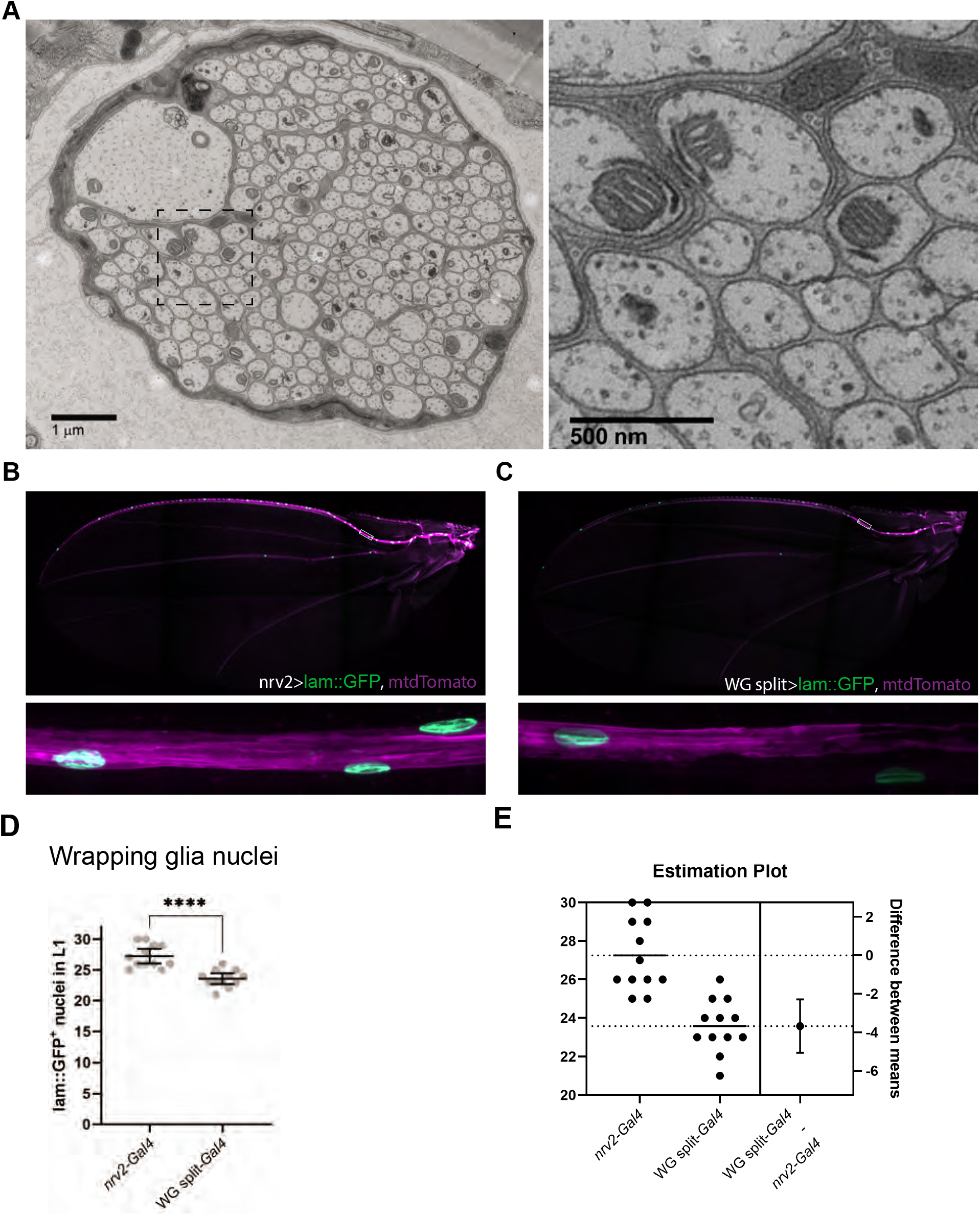
A) Raw electron micrographs from Fig. 1. B) Expression pattern of *nrv2* in the adult wing. C) Expression pattern of the wrapping glia split-*Gal4* in the adult wing. D) Quantification of the number of wrapping glia nuclei genetically labelled by *nrv2* and the WG split-*Gal4*. E) Estimation plot of the difference in labelling efficiency of wrapping glia nuclei for each *Gal4* construct within the wing. Statistics: Unpaired t test. ****= p < 0.0001.

**Supplemental Figure 2:**
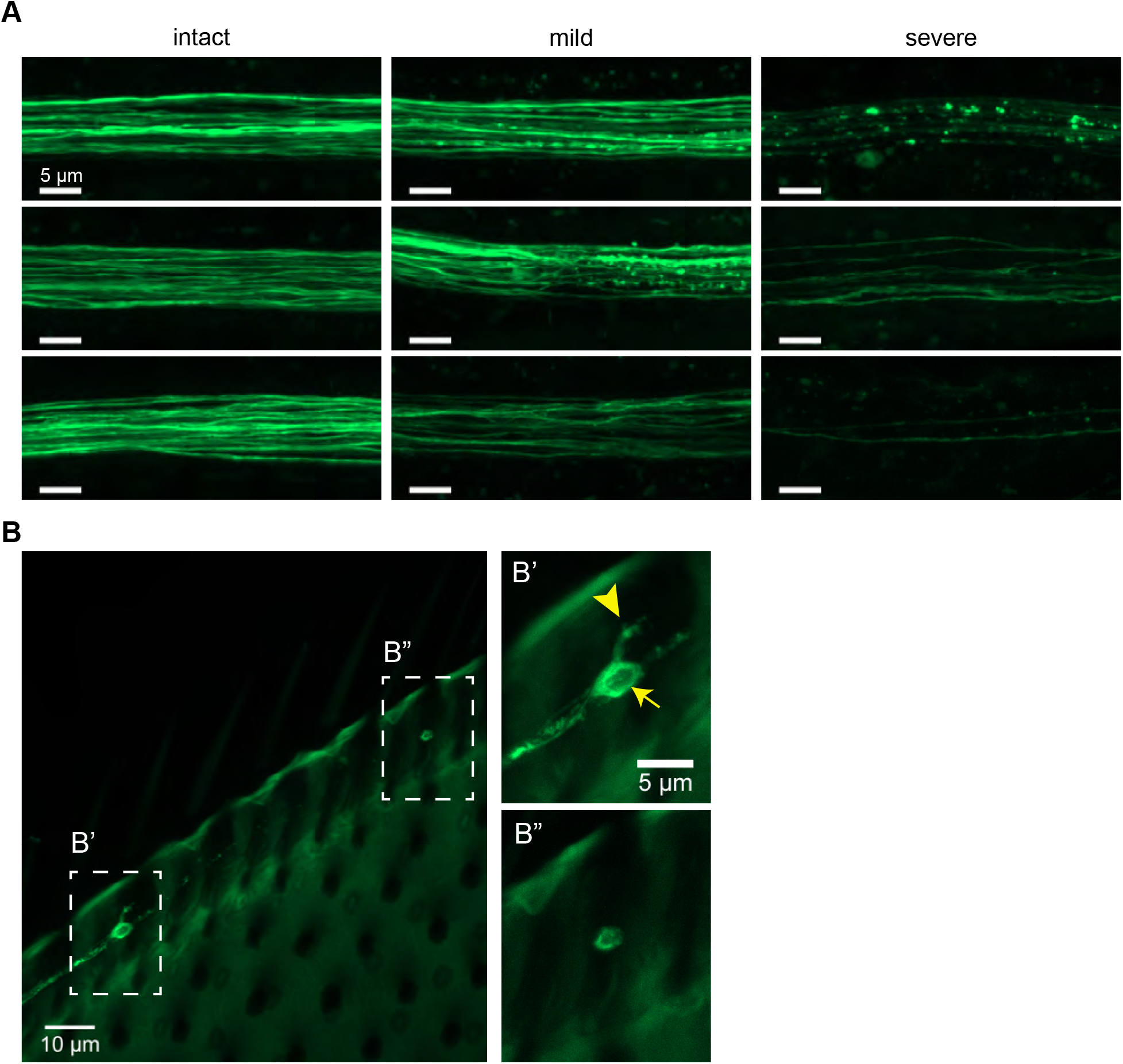
A) Examples of orthogonal projections of axons classified as intact, mild, or severe degeneration phenotype. All examples are from wrapping glia-ablated nerves, only the GFP channel is shown. B) Examples of an intact (B’) and neuron corpse (B”) within the same nerve. B’) Arrow indicates the nucleus, arrowhead indicates dendrite.

**Supplemental figure 3:**
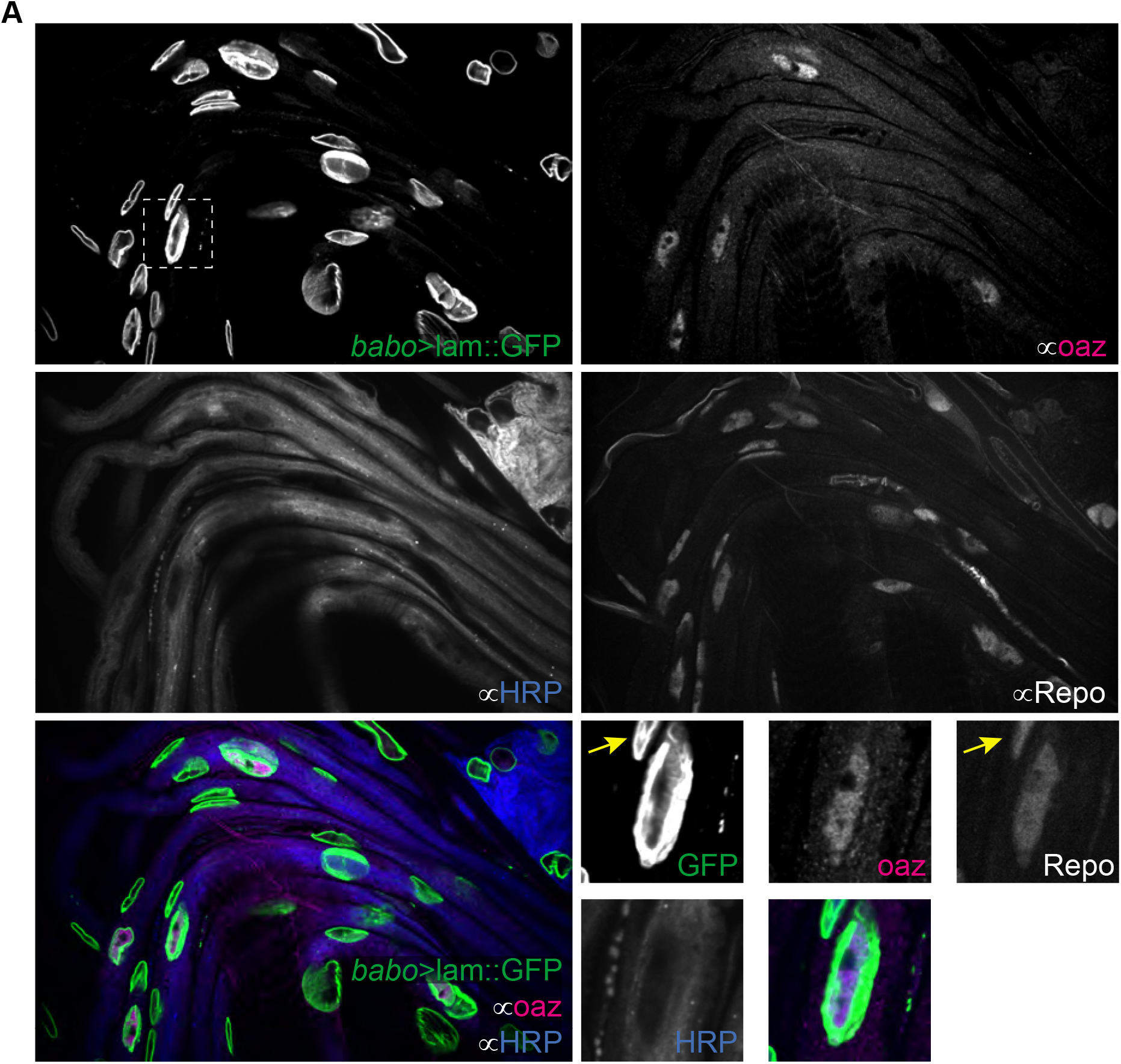
A) nerves from wandering L3 animals expressing a nuclear membrane teathered GFP in *babo^+^* cells. Nerves were co-stained for oaz (WG nuclei), HRP (neurons), and Repo (all glial nuclei). The region of interest indicated by the box is shown at higher magnification (bottom right panel). Yellow arrow indicates a GFP^+^/oaz^-^/Repo^+^ nucleus.

**Supplemental Figure 4:**
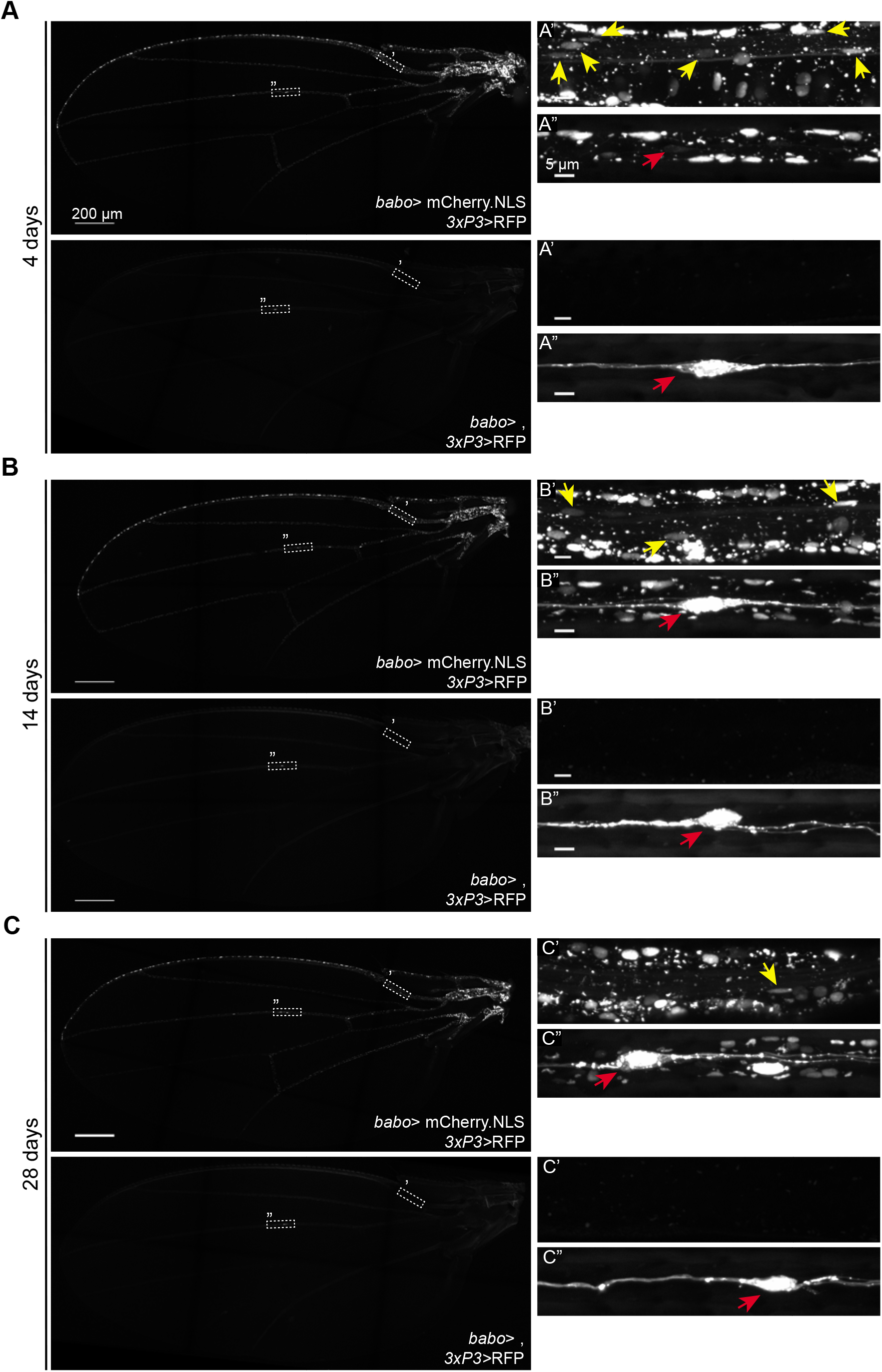
*babo-Gal4* expression is distinct from *3xP3>*RFP expression. A-C) Expression patterns of *babo*-driven nuclear mCherry versus *3xP3*-driven RFP within the wing at 4 (A), 14 (B), and 28 (C) days. Nuclear mCherry is present throughout the wing (top) whereas RFP is expressed within one cell in the L3 vein (bottom). Corresponding ROIs (’&”) are shown to the right. At all timepoints mCherry^+^ nuclei were present in the nerve region (yellow arrows) and was distinct from RFP^+^ cells in the L3 vein (red arrows).

**Supplemental Figure 5:**
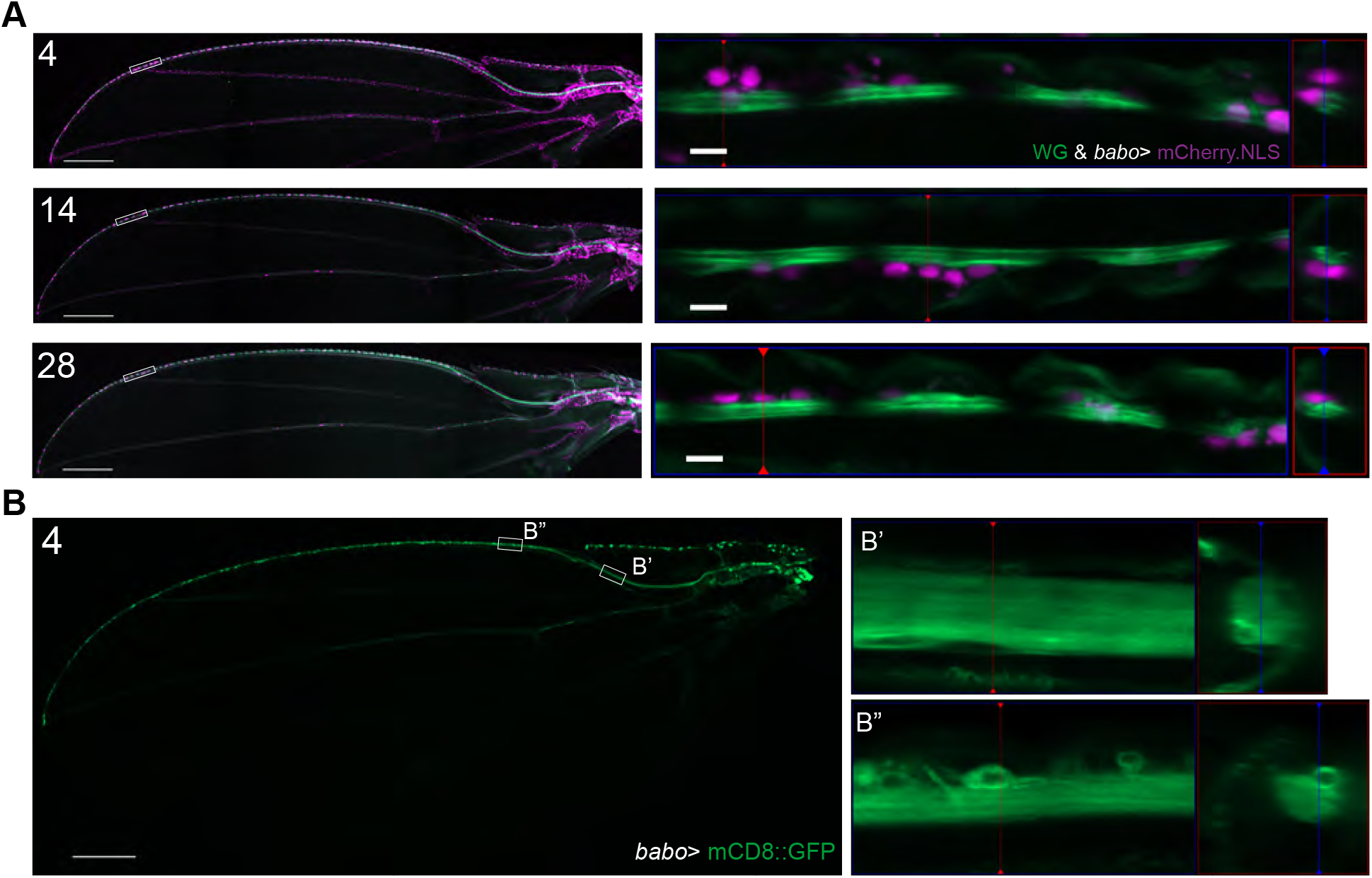
A) *babo* nuclear reporter expression in the wing at 4 (top), 14 (middle), and 28 (bottom) days. ROIs from boxes shown to the right. mCherry^+^ nuclei residing outside of WG GFP^+^ membrane resemble neuronal nuclei. B) *babo* membrane reporter expression in the adult wing at 4 days. ROIs from boxes shown to the right showing dense labeling throughout the nerve bundle (B’) and GFP^+^ cells that resemble neurons (B”).

**Supplemental Figure 6:**
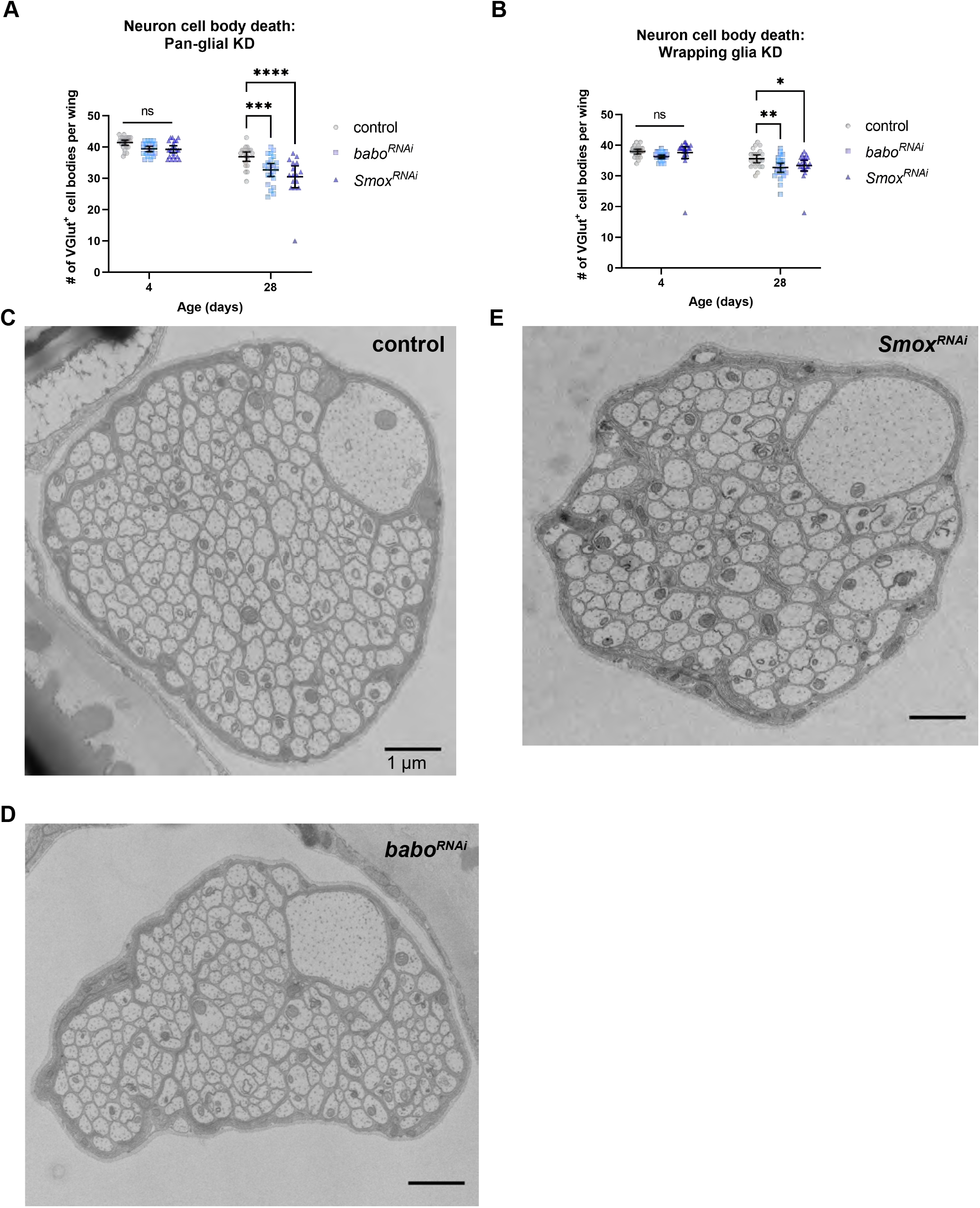
Glial ensheathement is unchanged in TGFβ knockdown conditions. A) Quantification of the number of intact VGlut+ neuron cell bodies in the pan-glial (A) and wrapping glia (B) knockdown conditions. C-D) Raw electron microcgraphs corresponding to pseudo-colored micrographs in Figure 6. Control (C) *babo^RNAi^* (D) *Smox^RNAi^* (E) Scale bar 1µm.

**Supplemental Figure 7:**
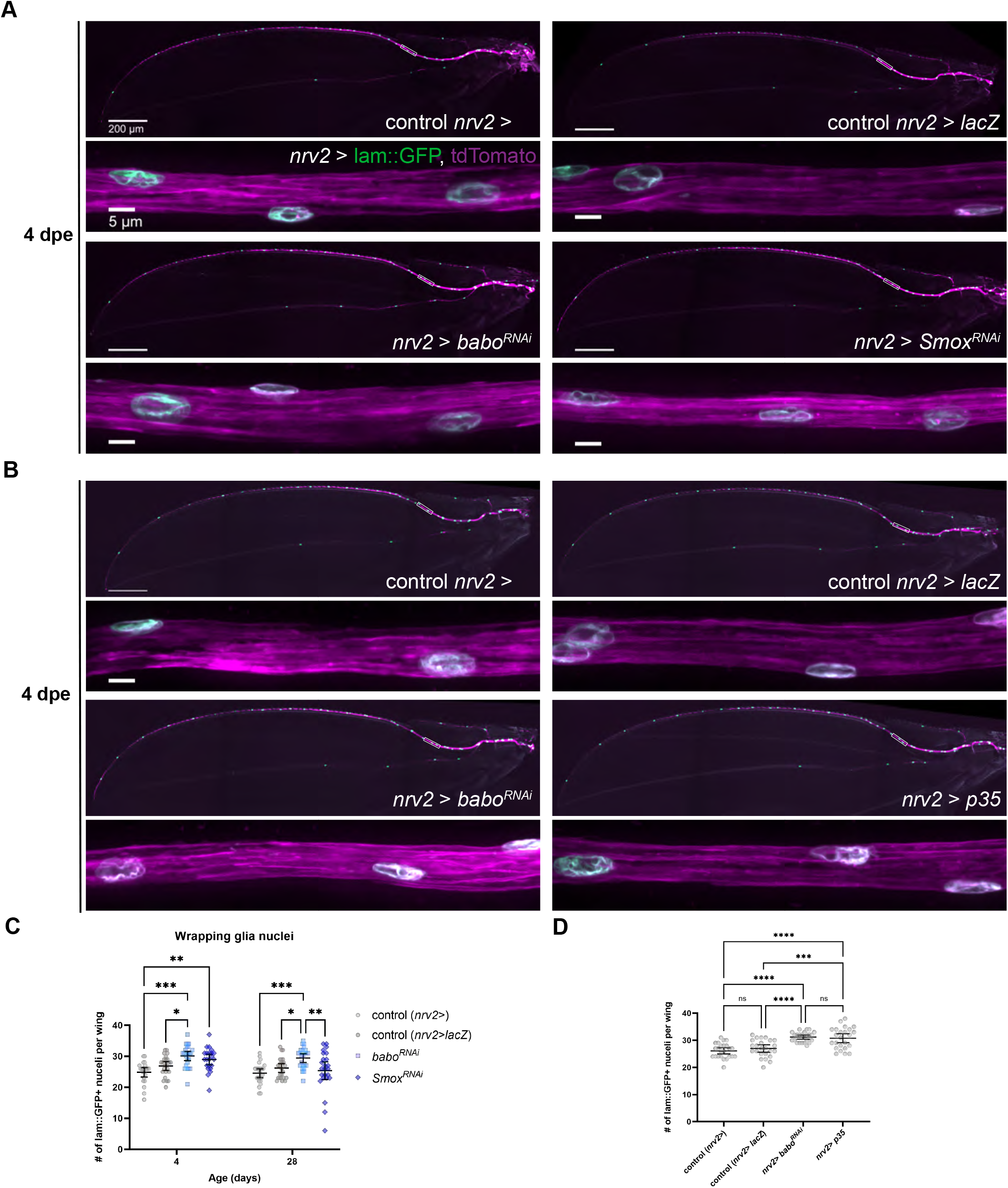
Greater number of wrapping glia in TGFβ knockdown animals. A) Images from 4 dpe wings expressing nuclear GFP and membrane-tethered tdTomato in wrapping glia in control (n/a & lacZ) and TGFβ knockdown animals. Higher magnification images from the boxed area are shown below each wing. B) Images from 4 dpe wings in control and *babo^RNAi^* or *p35* conditions. C) Quantification of the number of GFP^+^ wrapping glia nuclei per wing. D) Quantification of the number of wrapping glia in control, *babo^RNAi^*, and *p35* conditions at 4 dpe. Graph: mean ± 95% CI. Statistics: 2-way ANOVA with Dunnett’s (C) or Tukey’s (D) multiple comparisons test. Significance: *= p < 0.05, **= p < 0.01, ***= p < 0.001, ****= p < 0.0001.

**Supplemental Figure 8:**
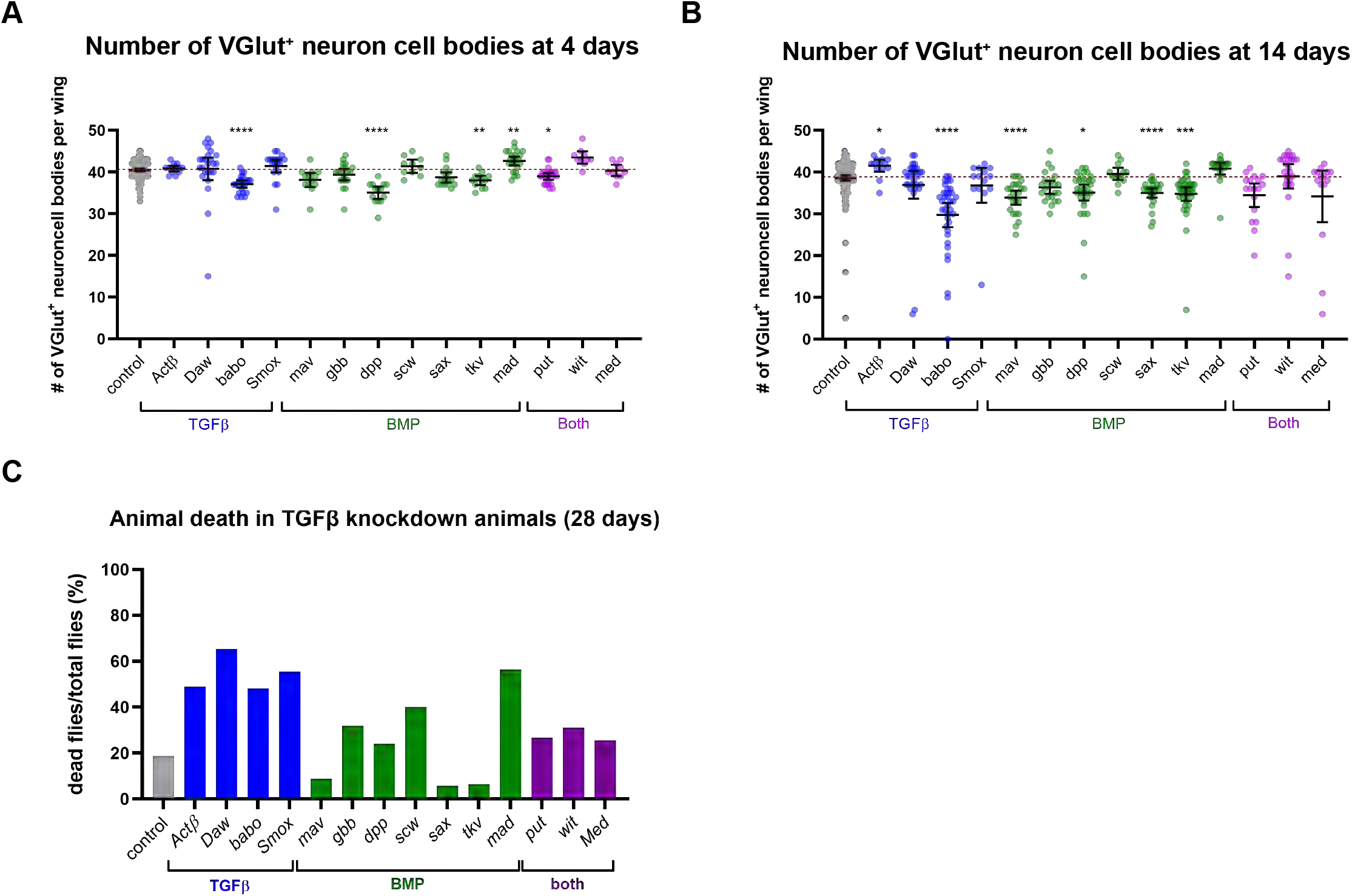
A-B) Quantification of the number of intact *VGlut*^+^ neuron cell bodies present per wing of 4 (A) and 14 (B) dpe nerves in control and pan-glial RNAi conditions. Data are represented as mean ± 95% CI. C) The percent of flies that died by 28 days of age for each control and RNAi-mediated knockdown condition.

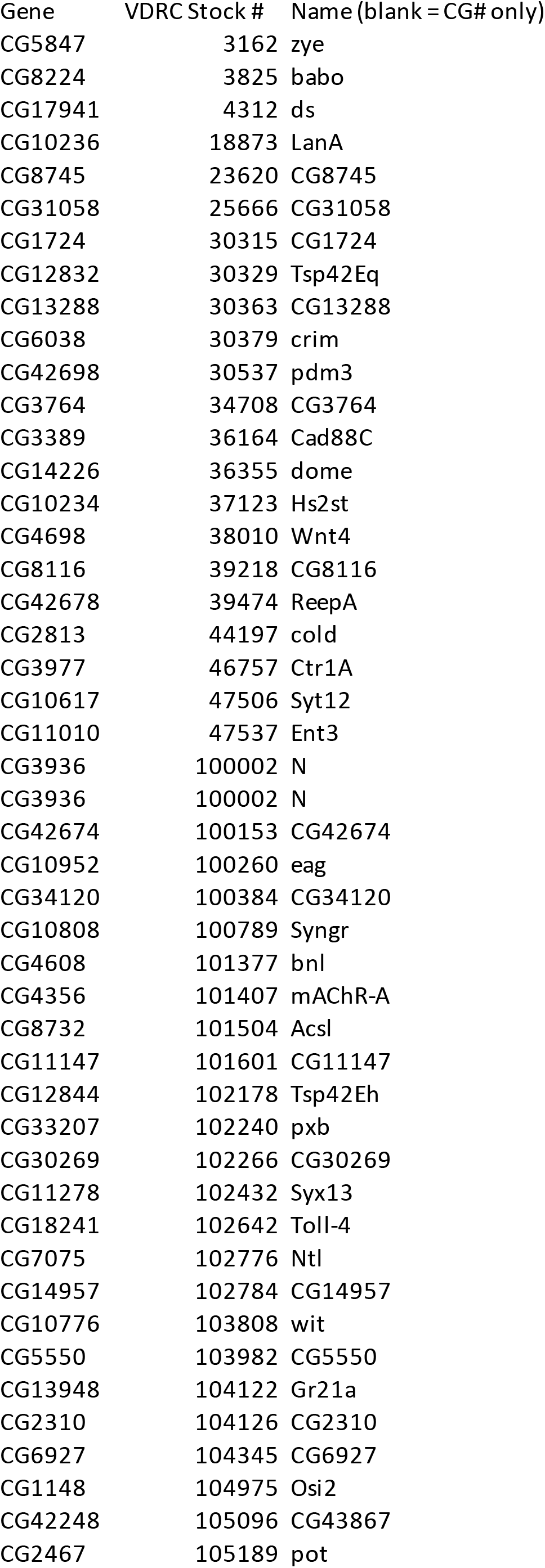
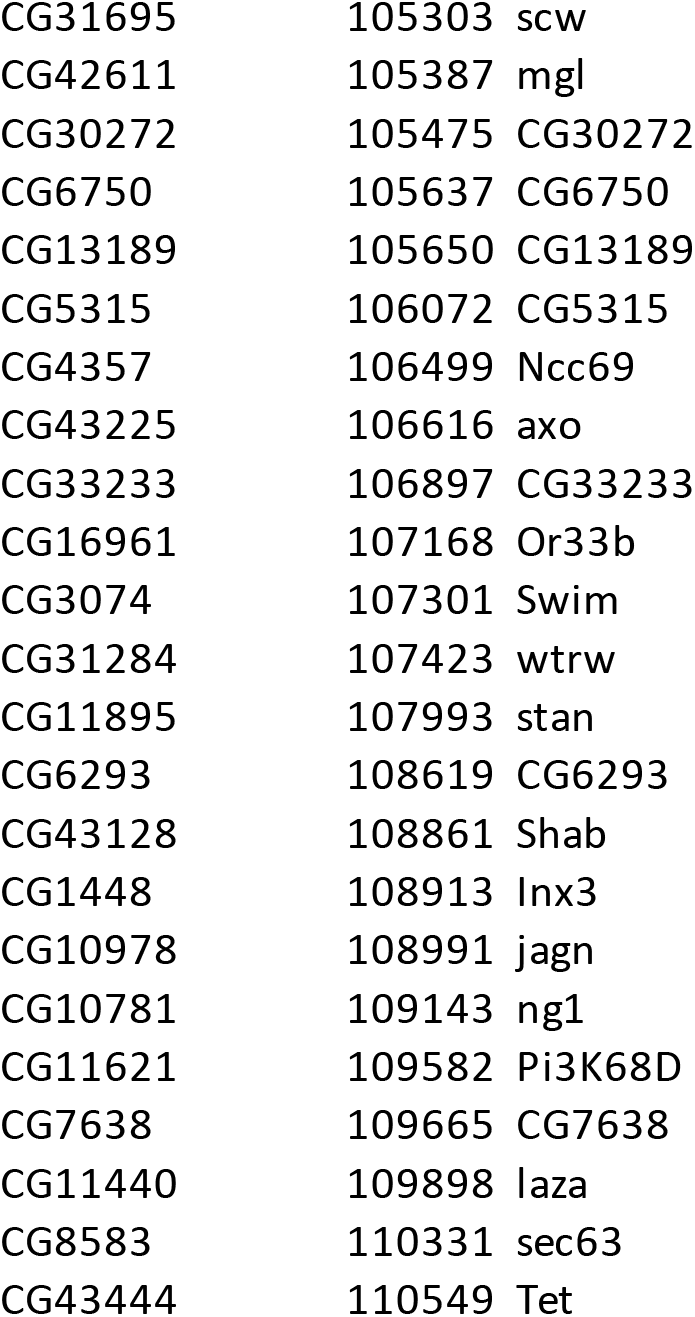

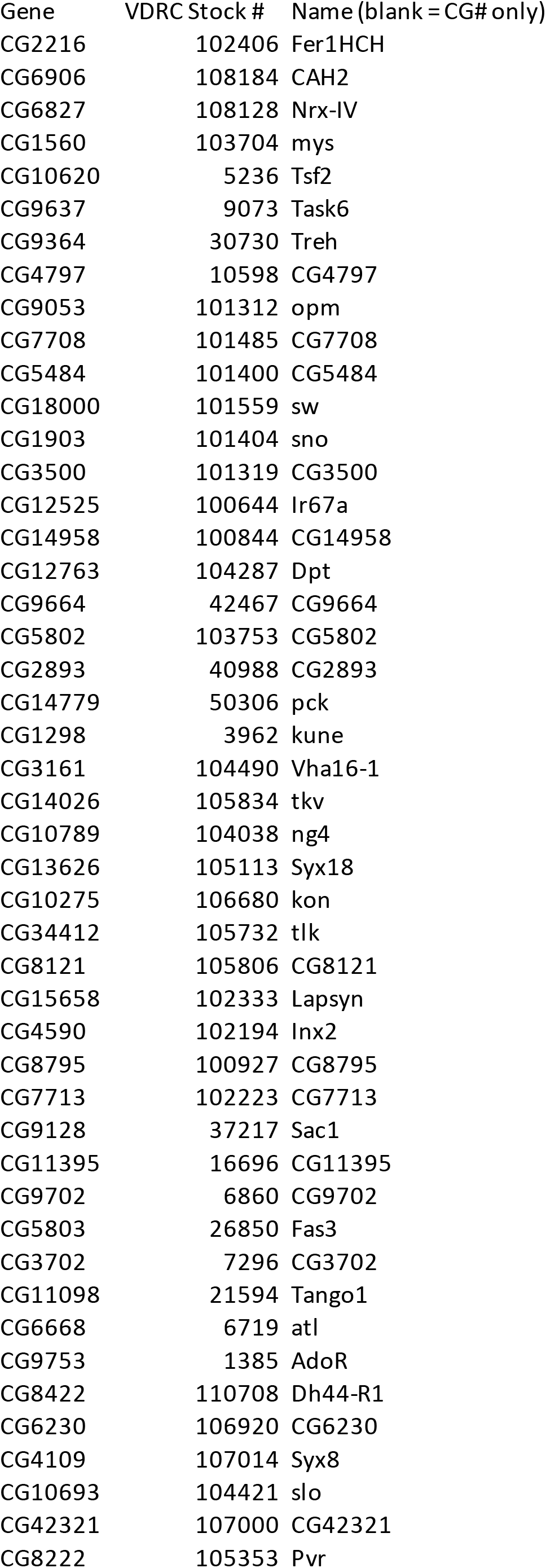
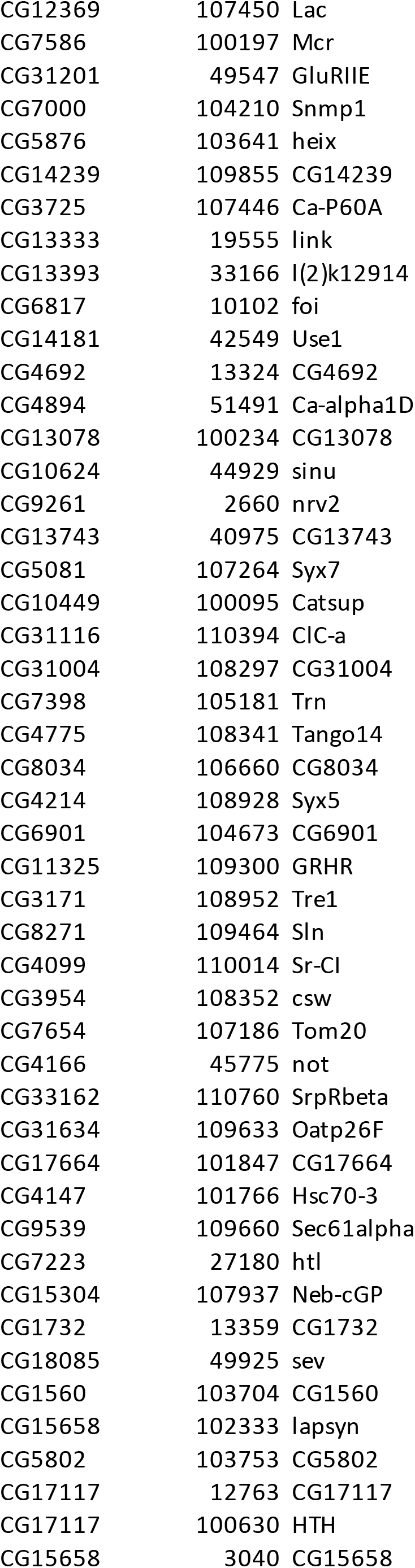
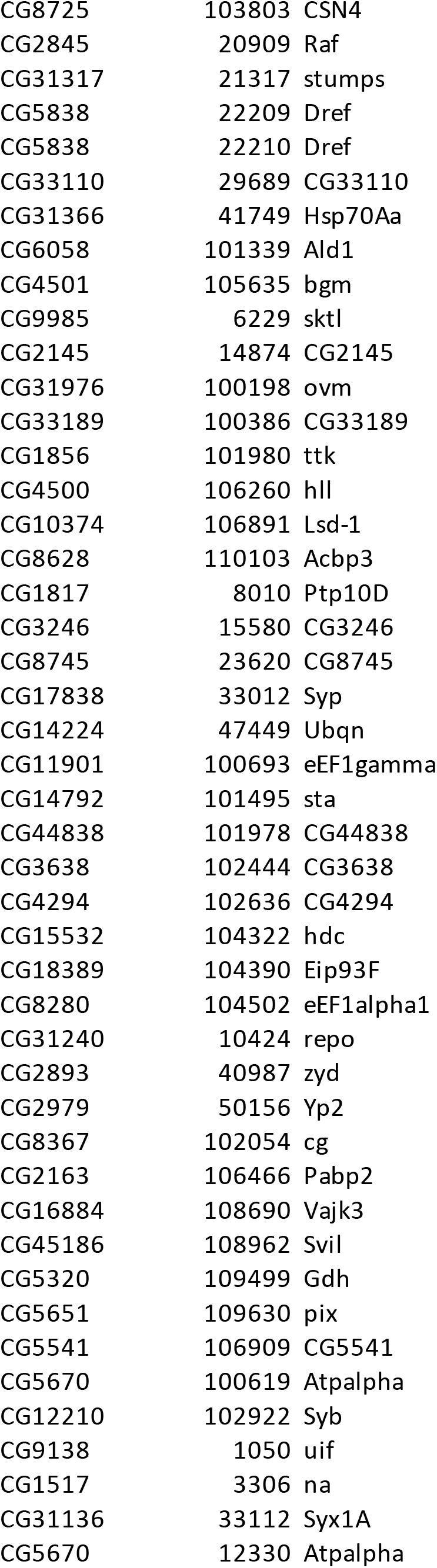

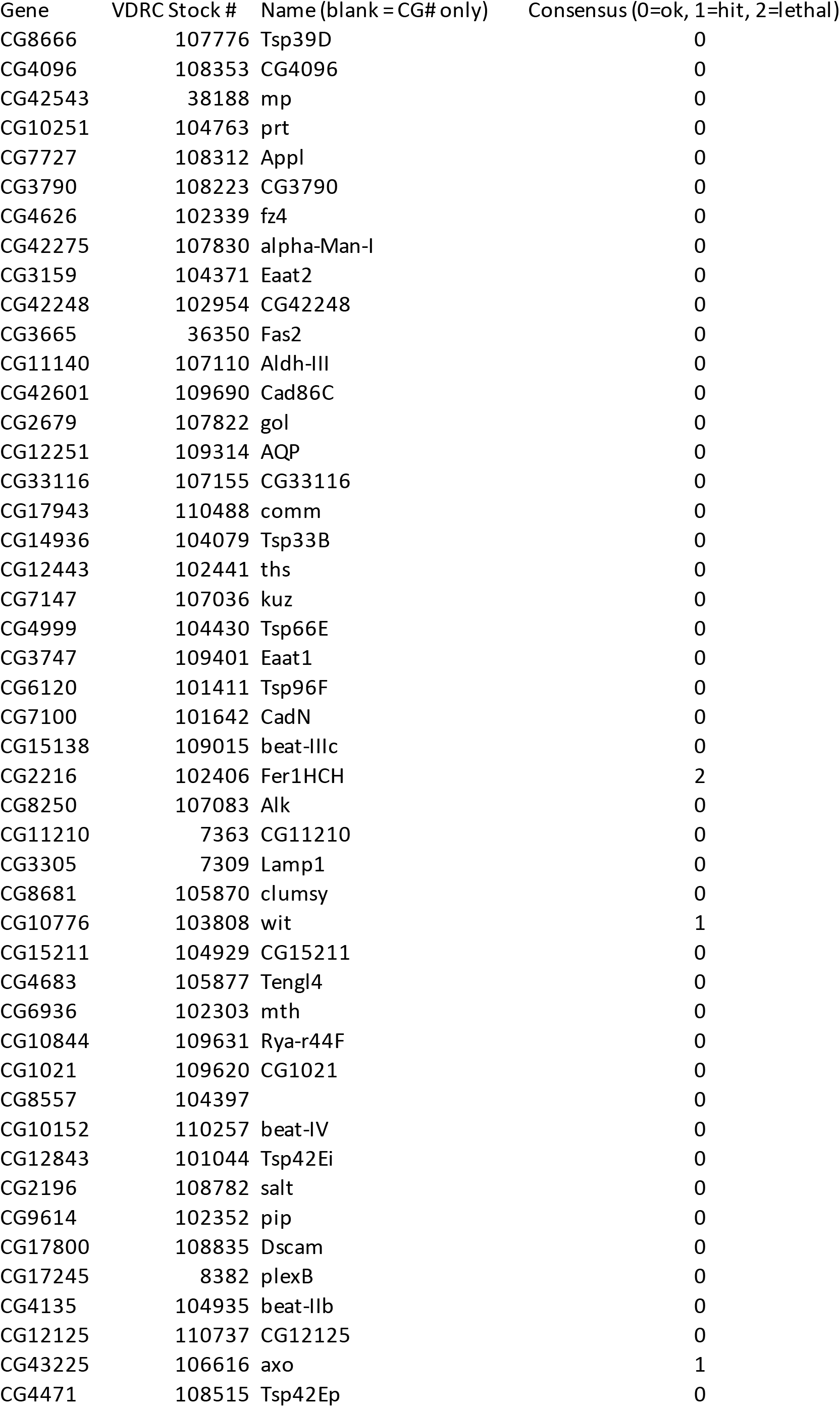
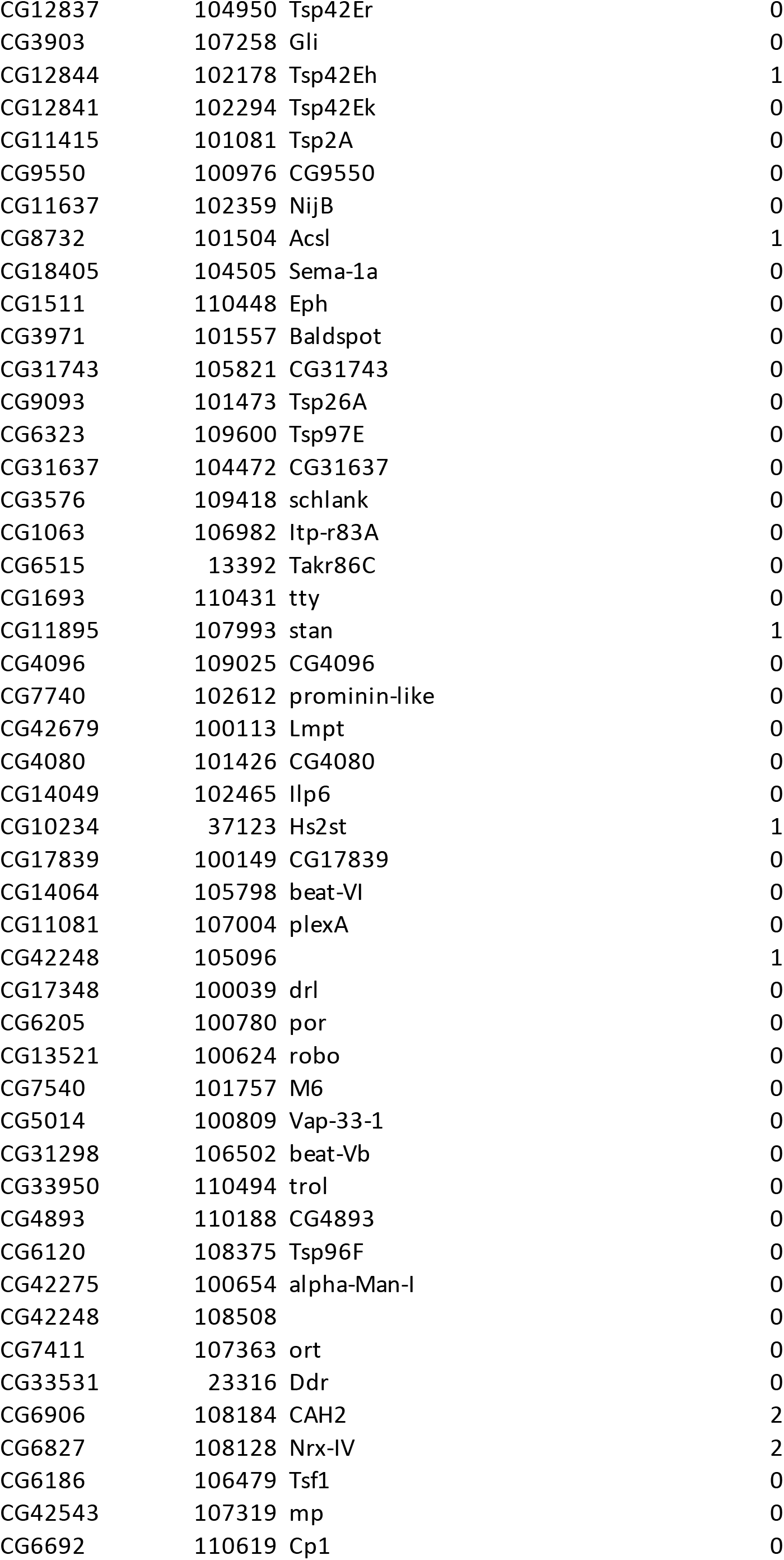
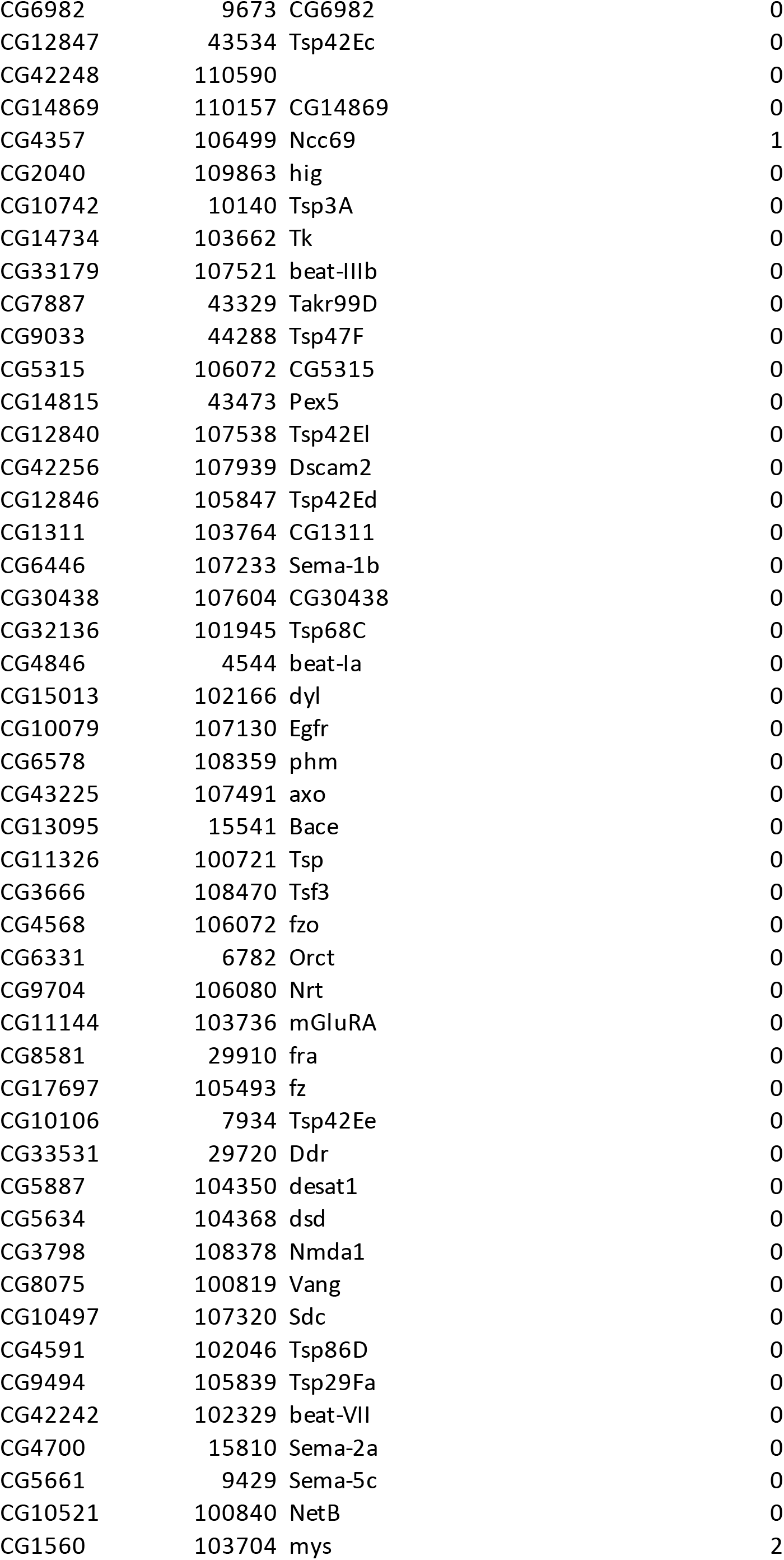
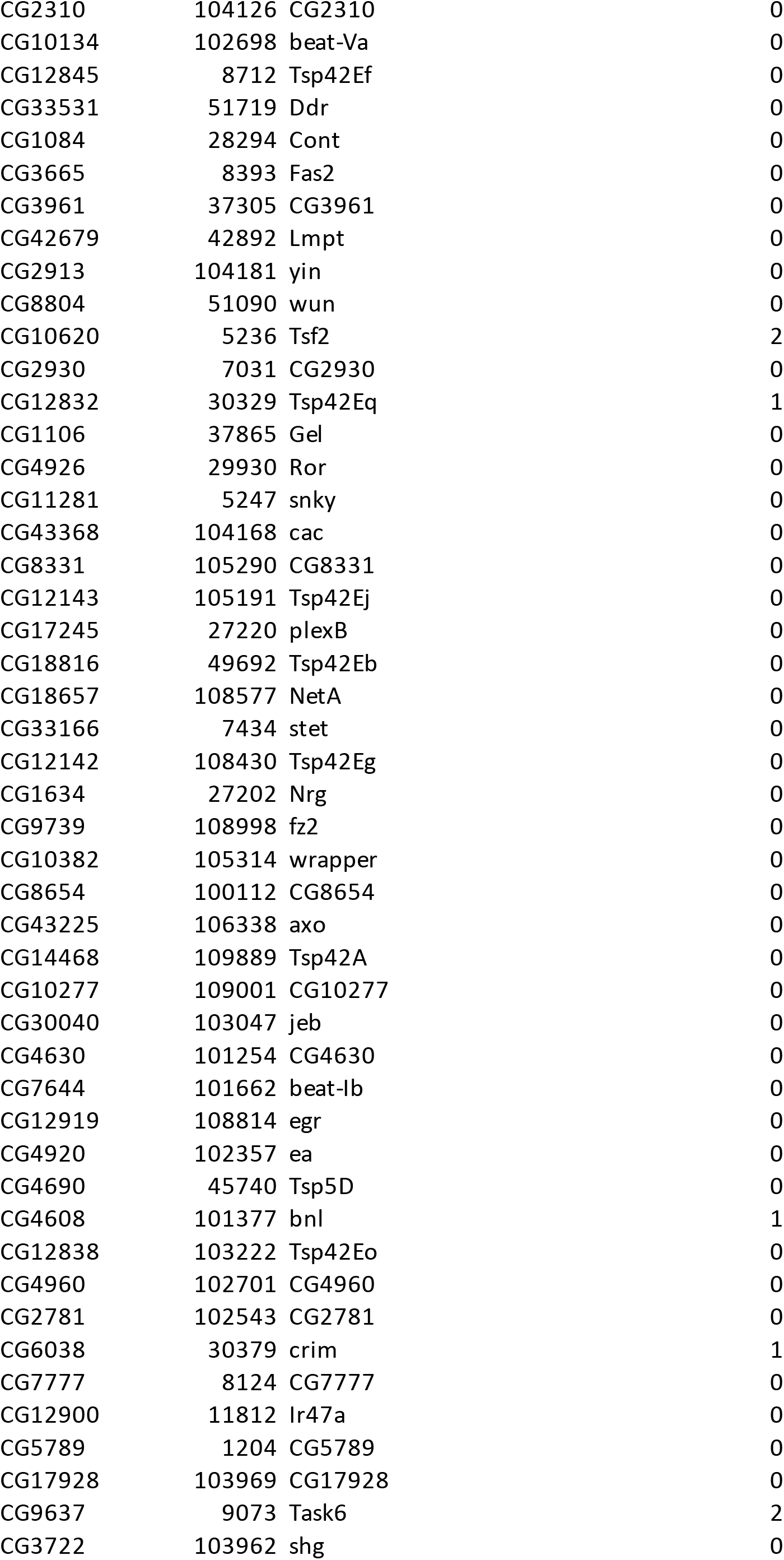
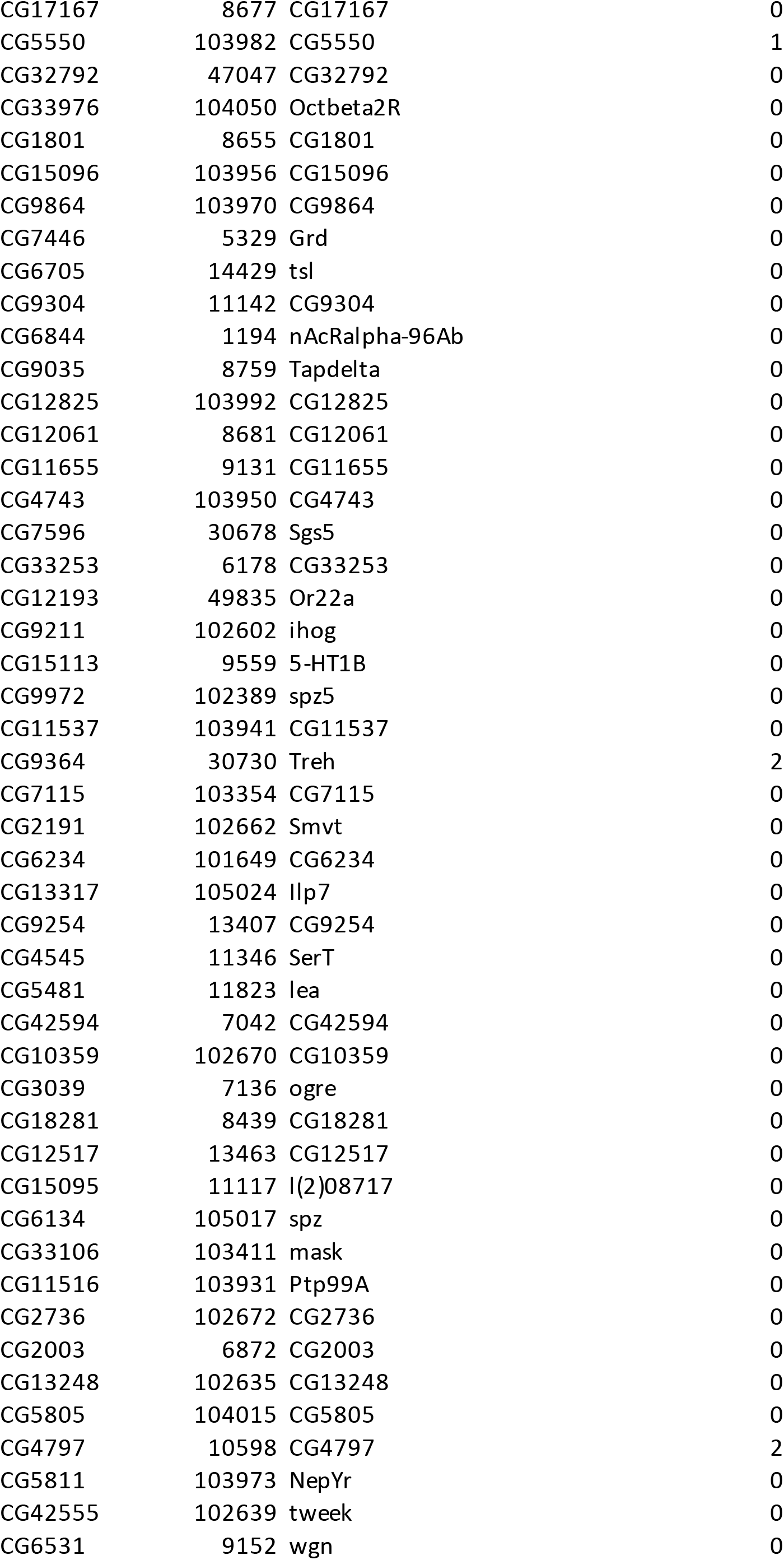
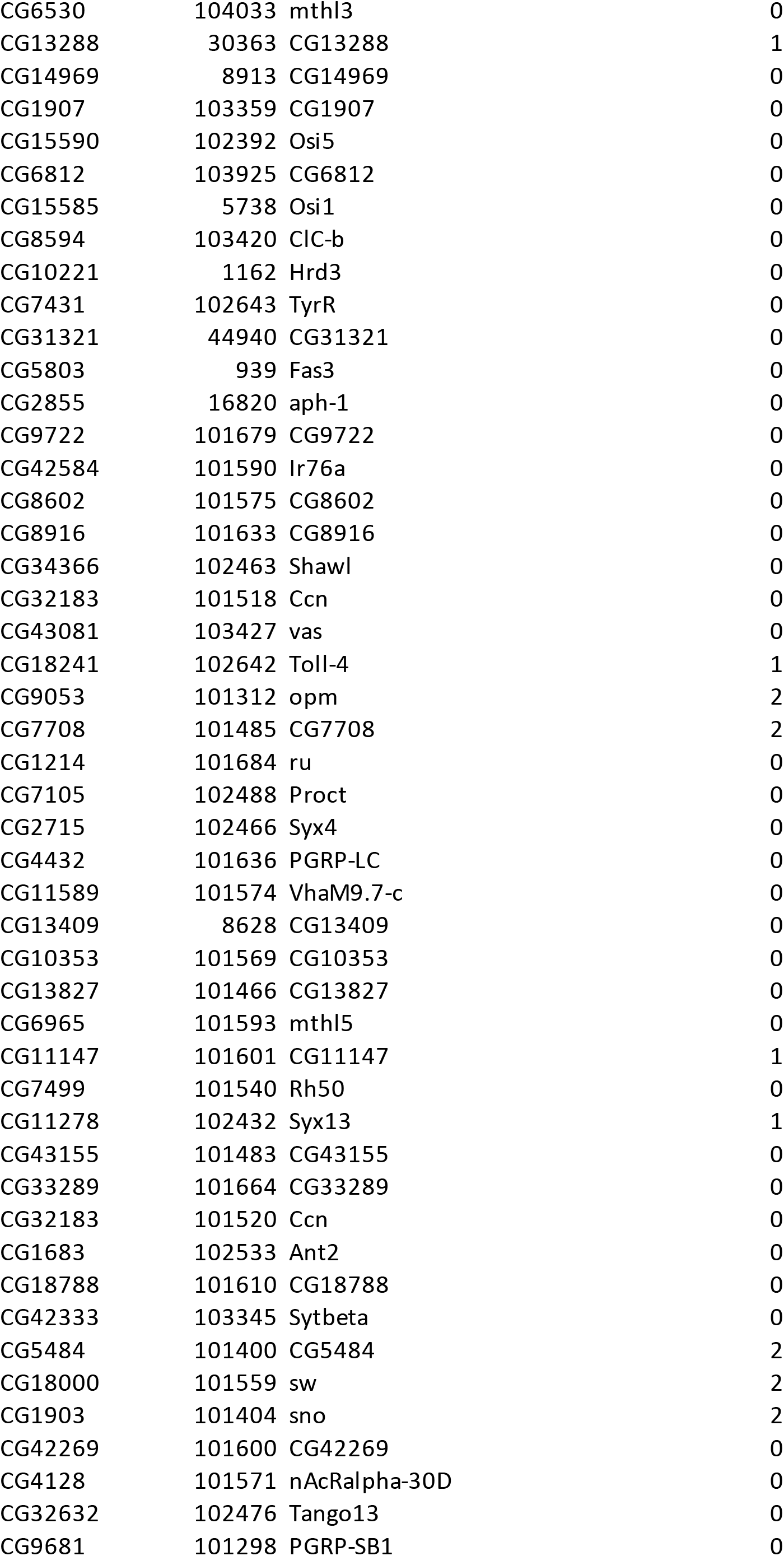
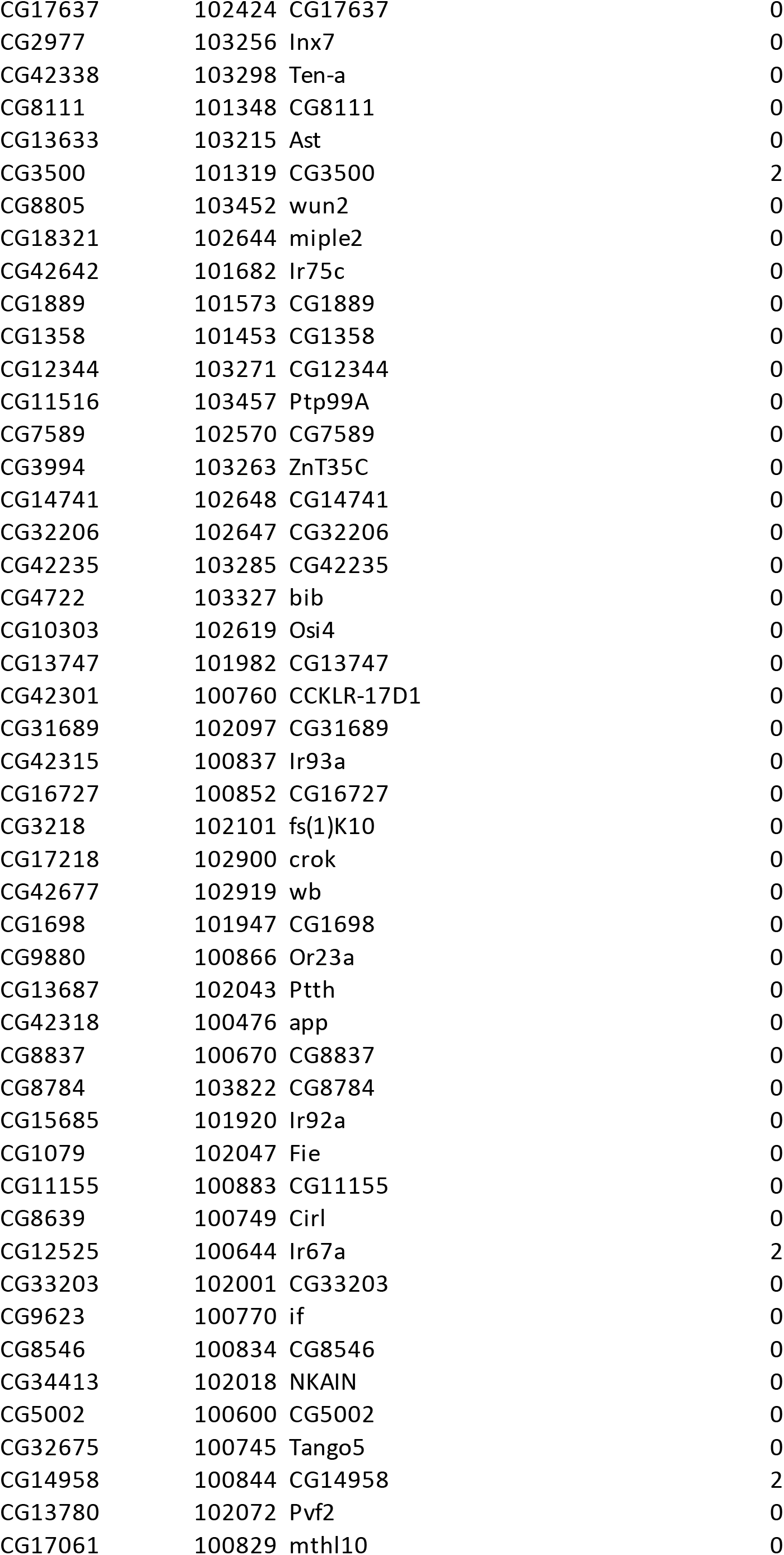
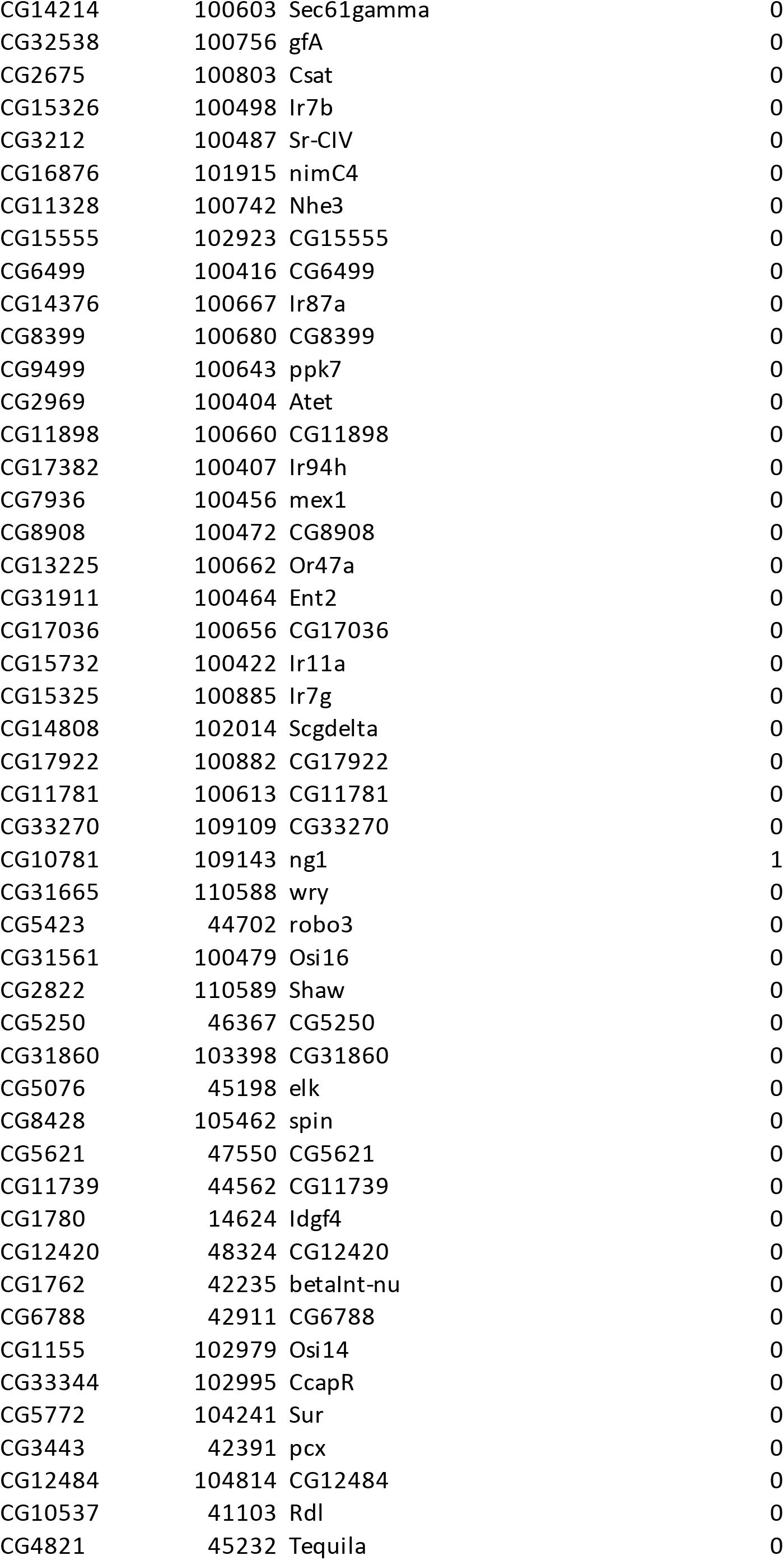
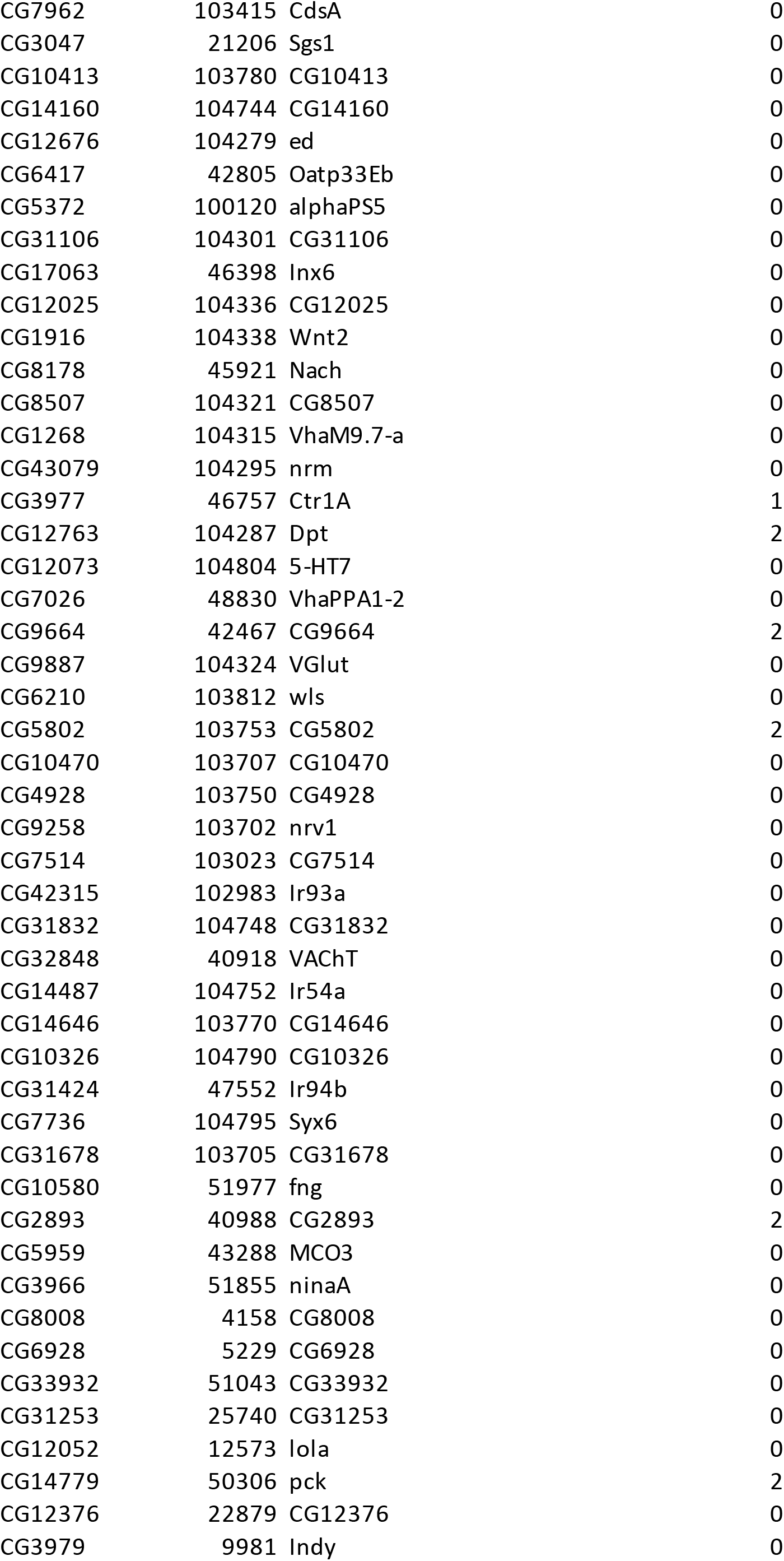
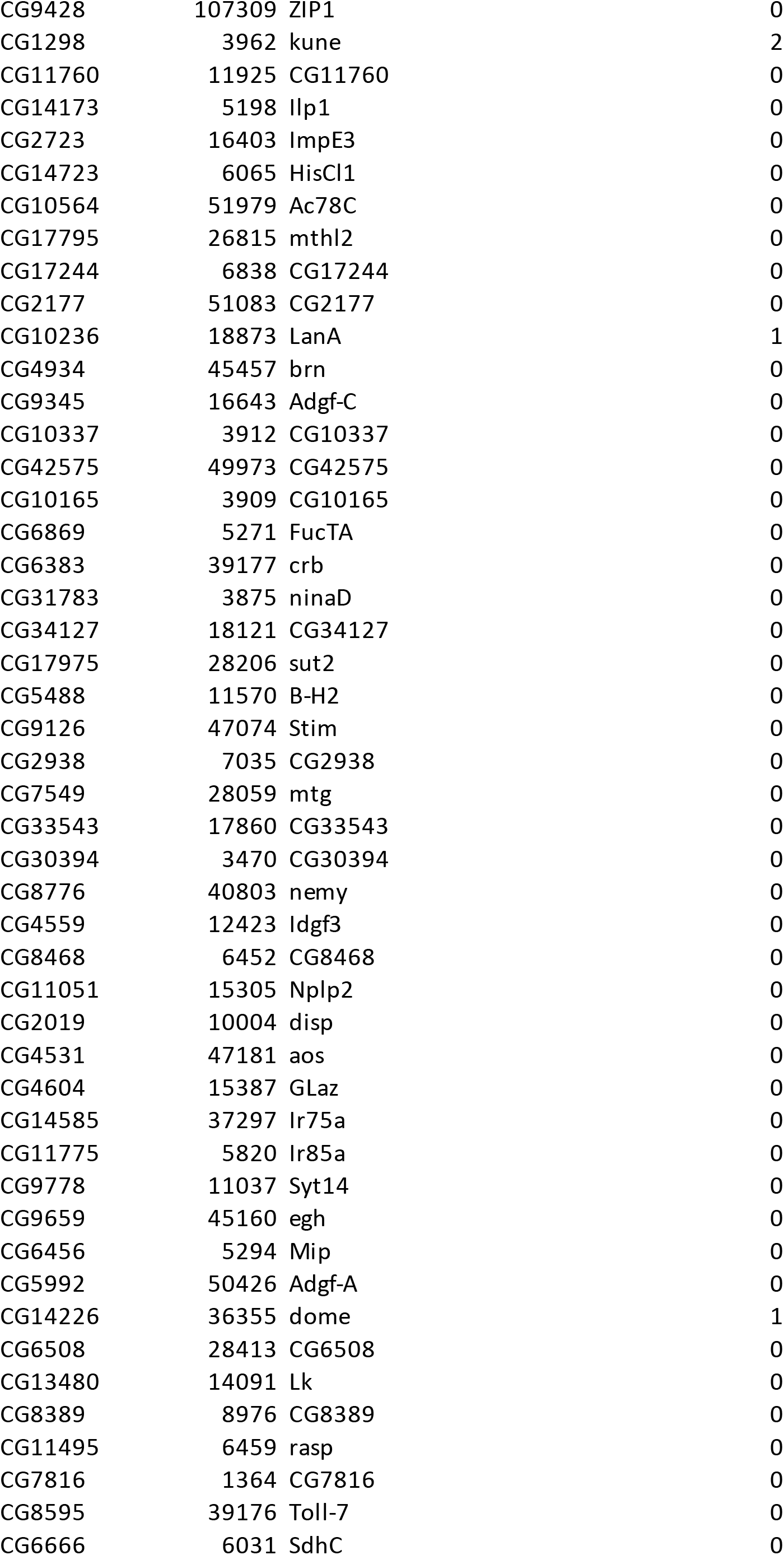
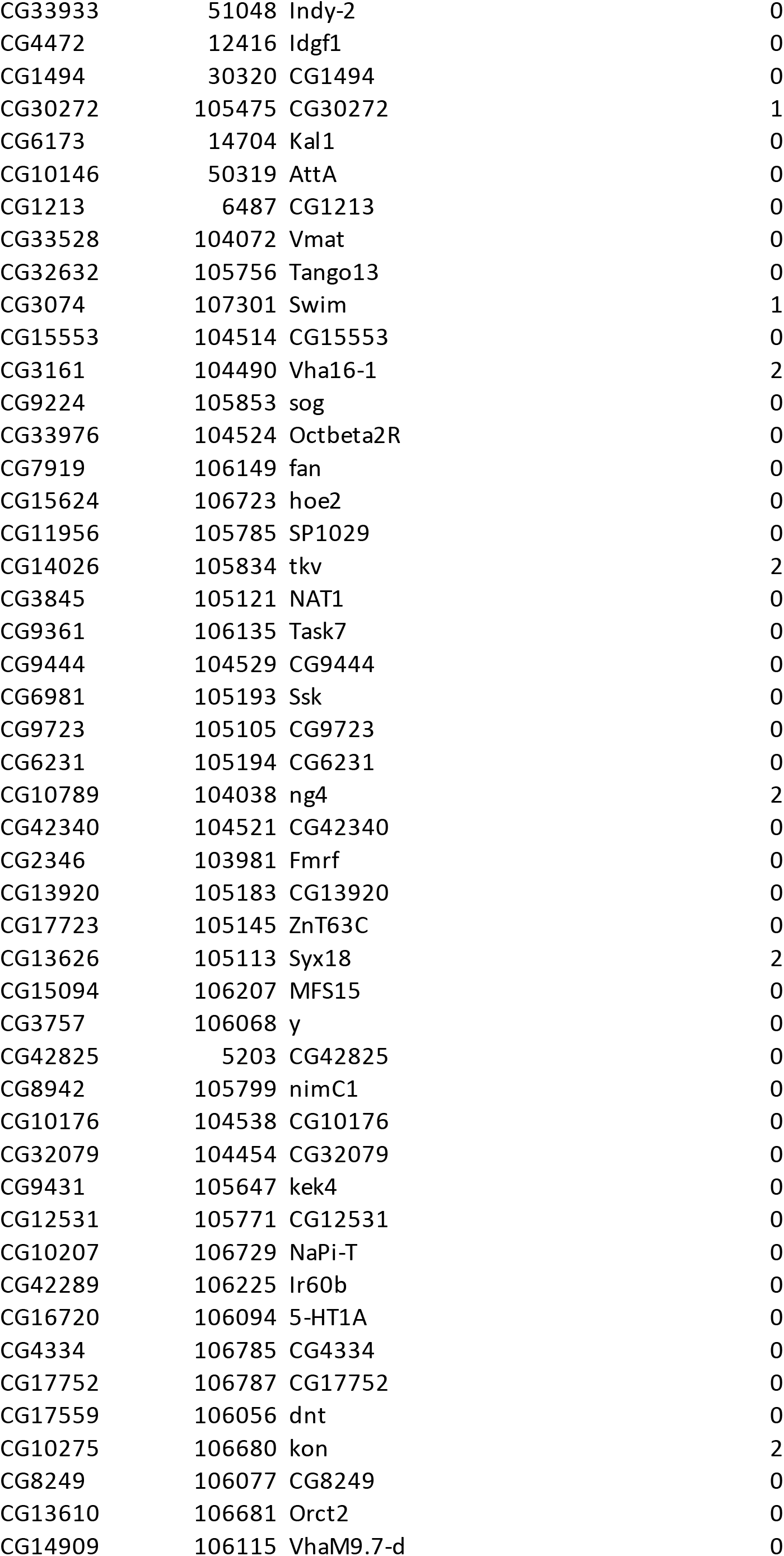
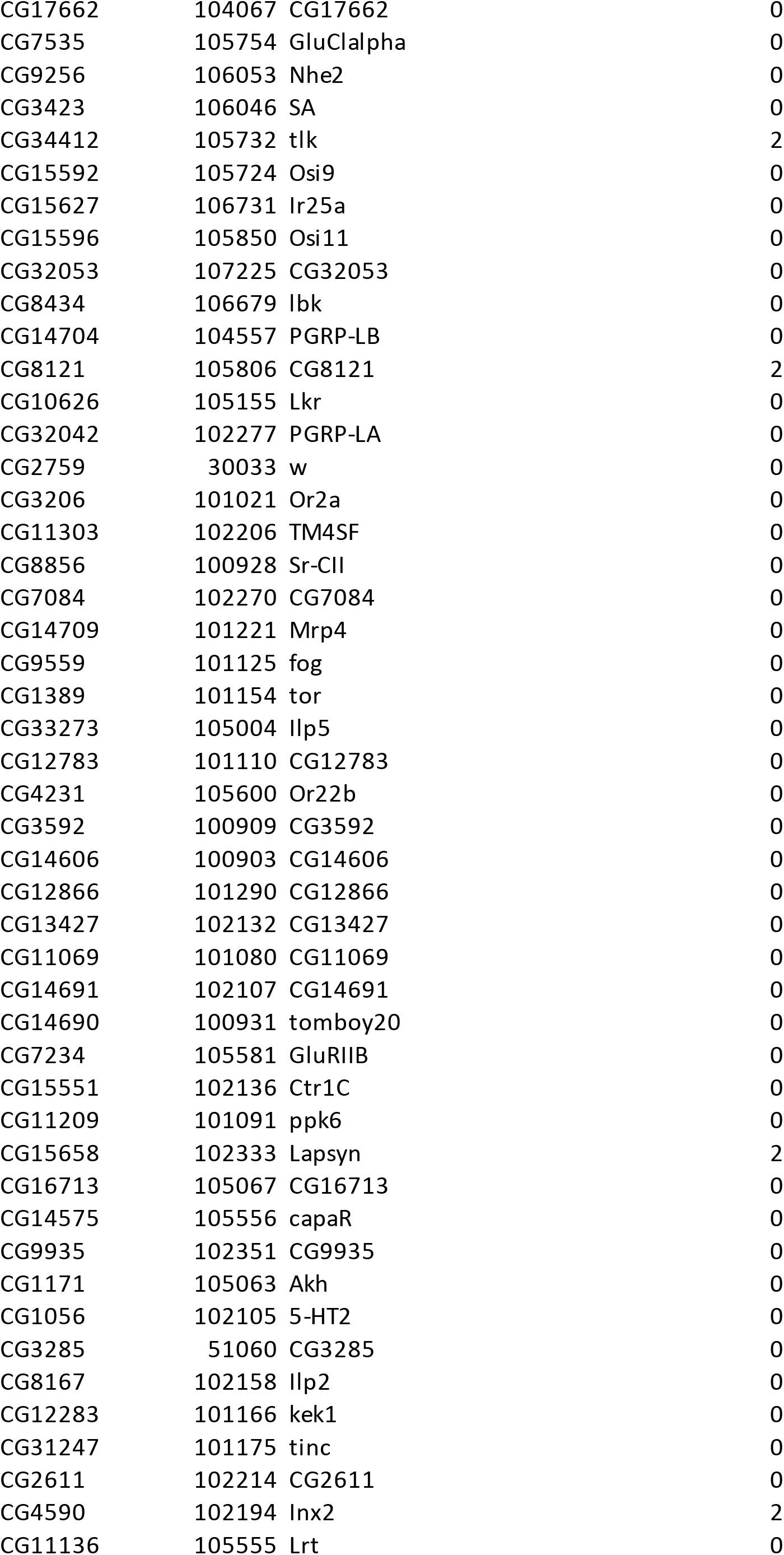
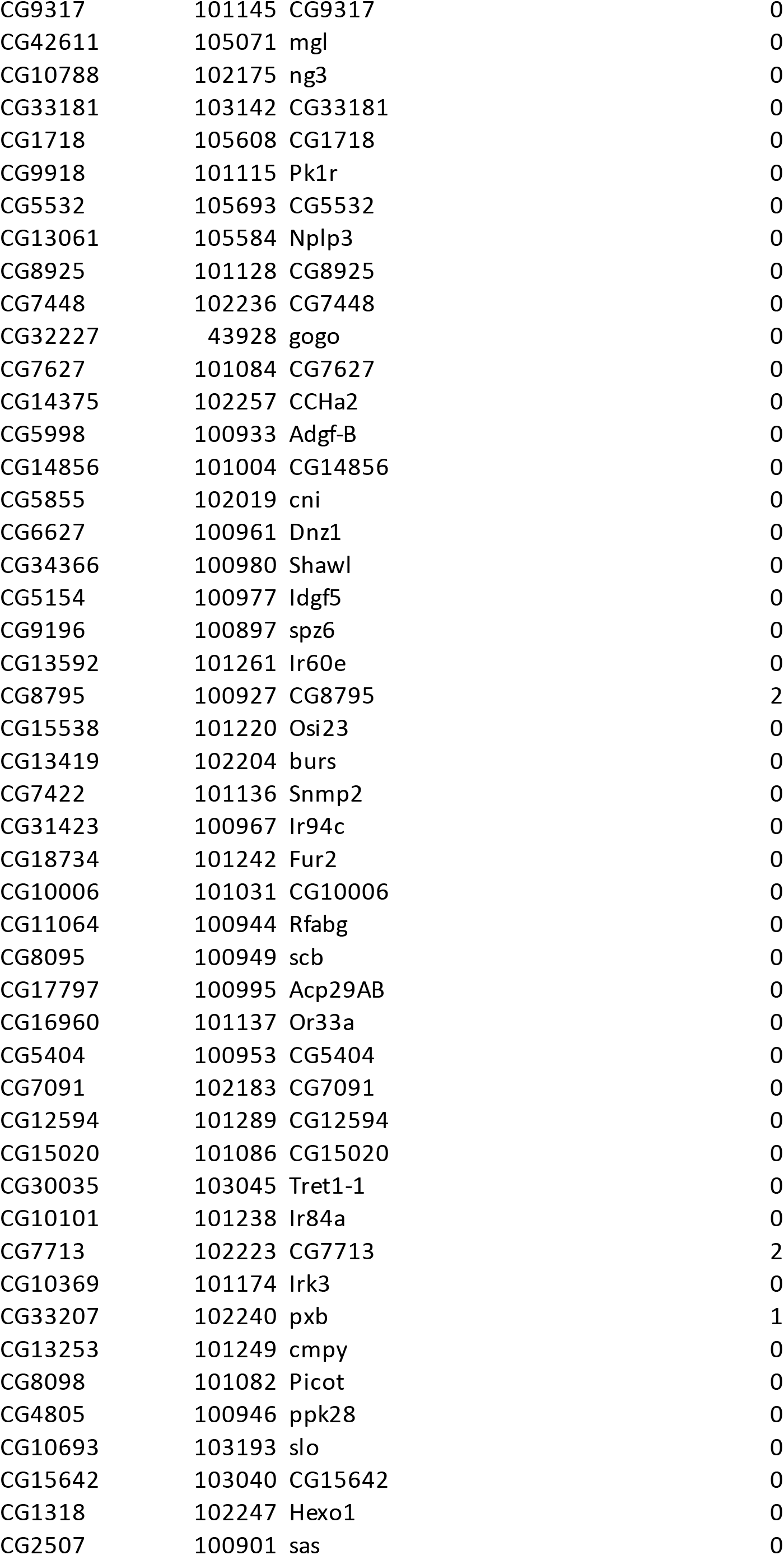
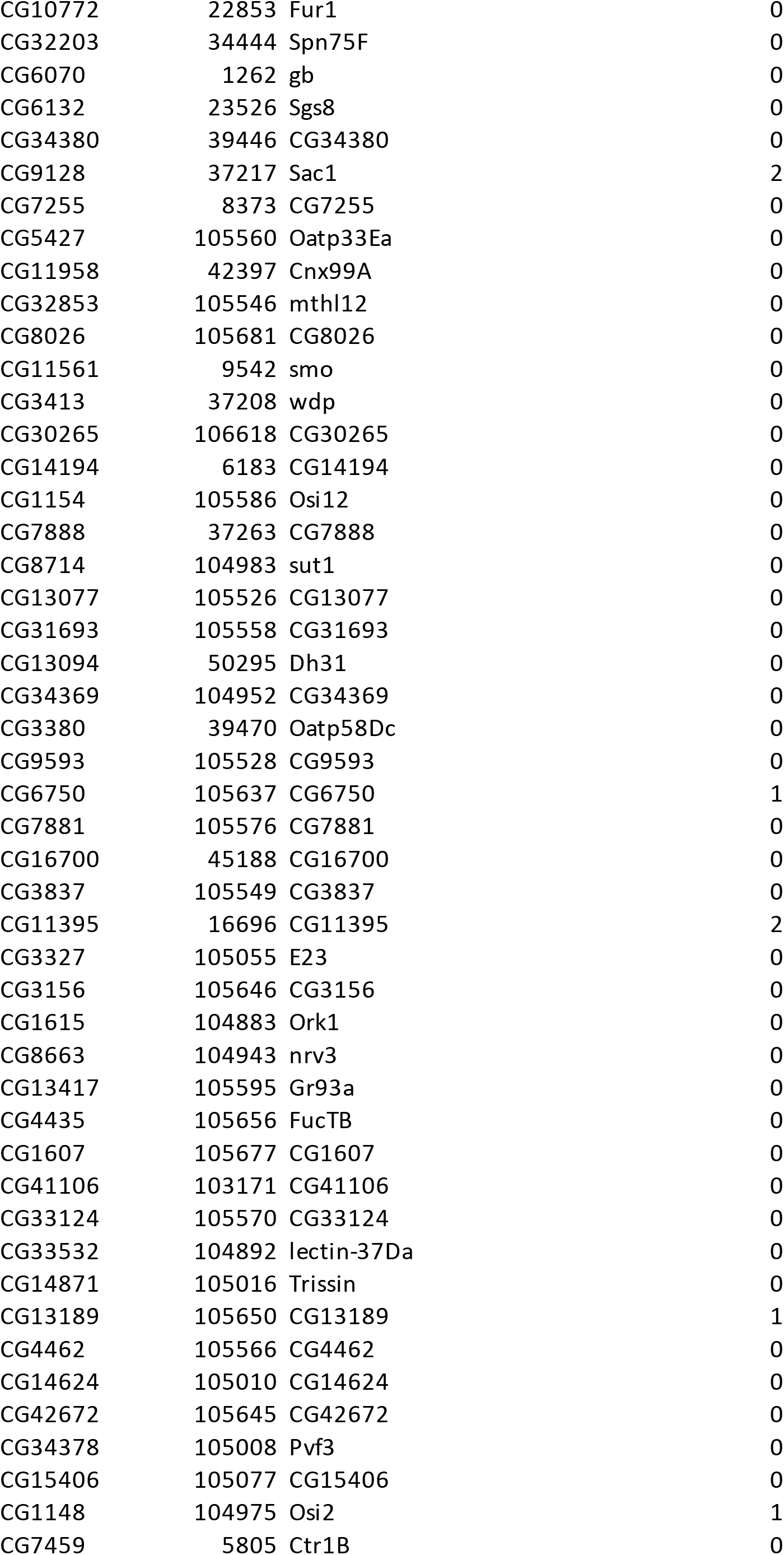
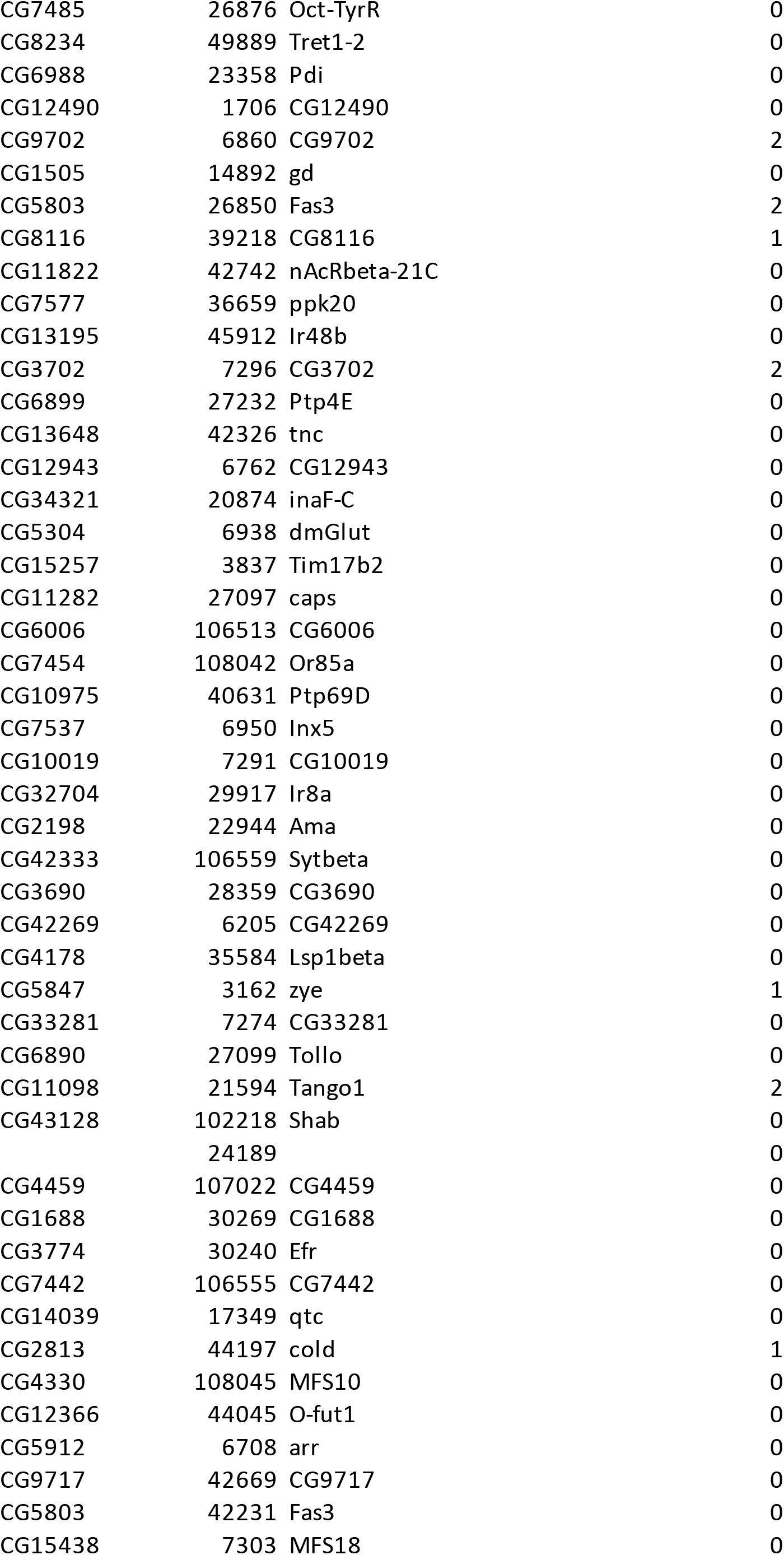
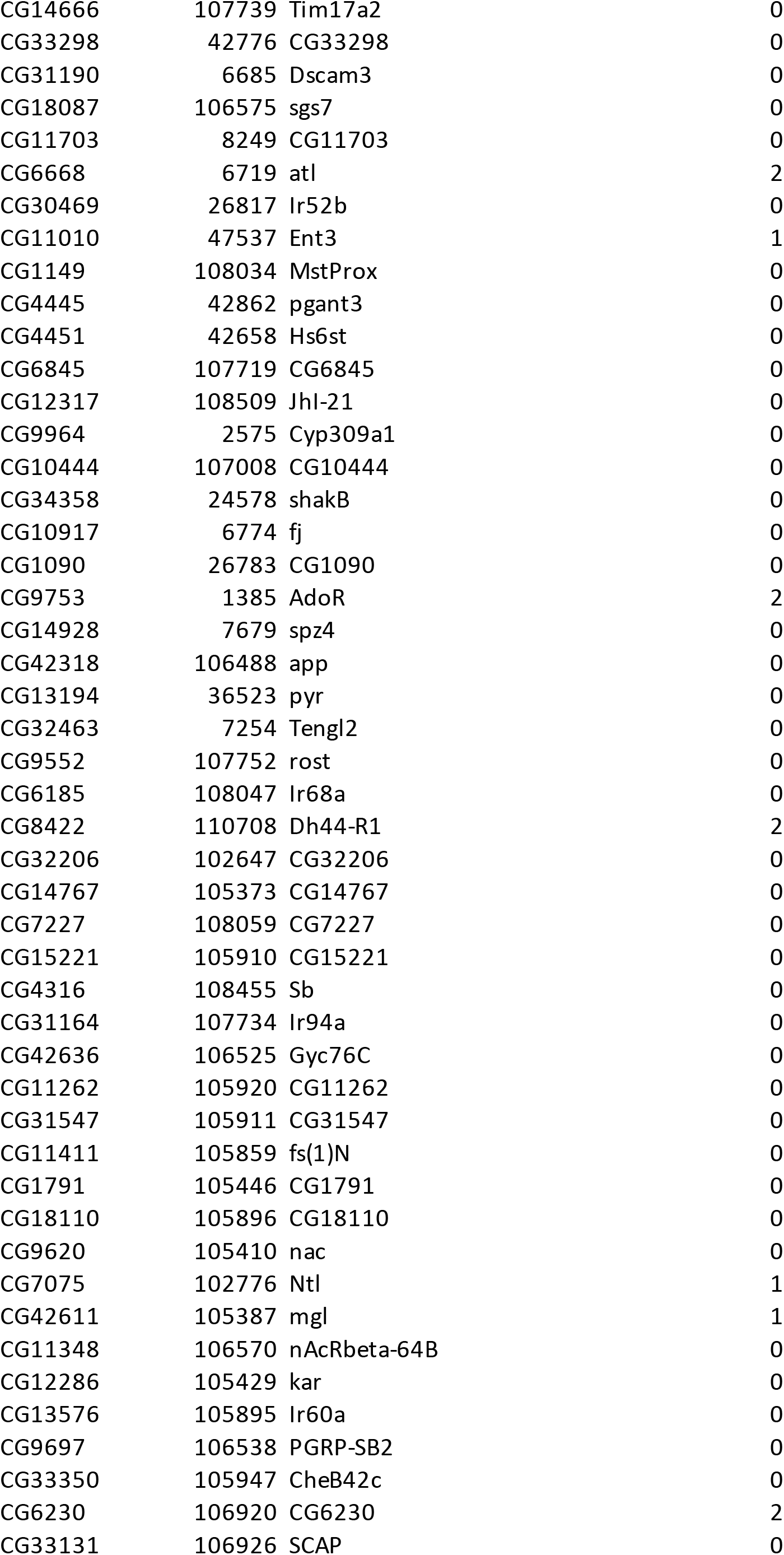
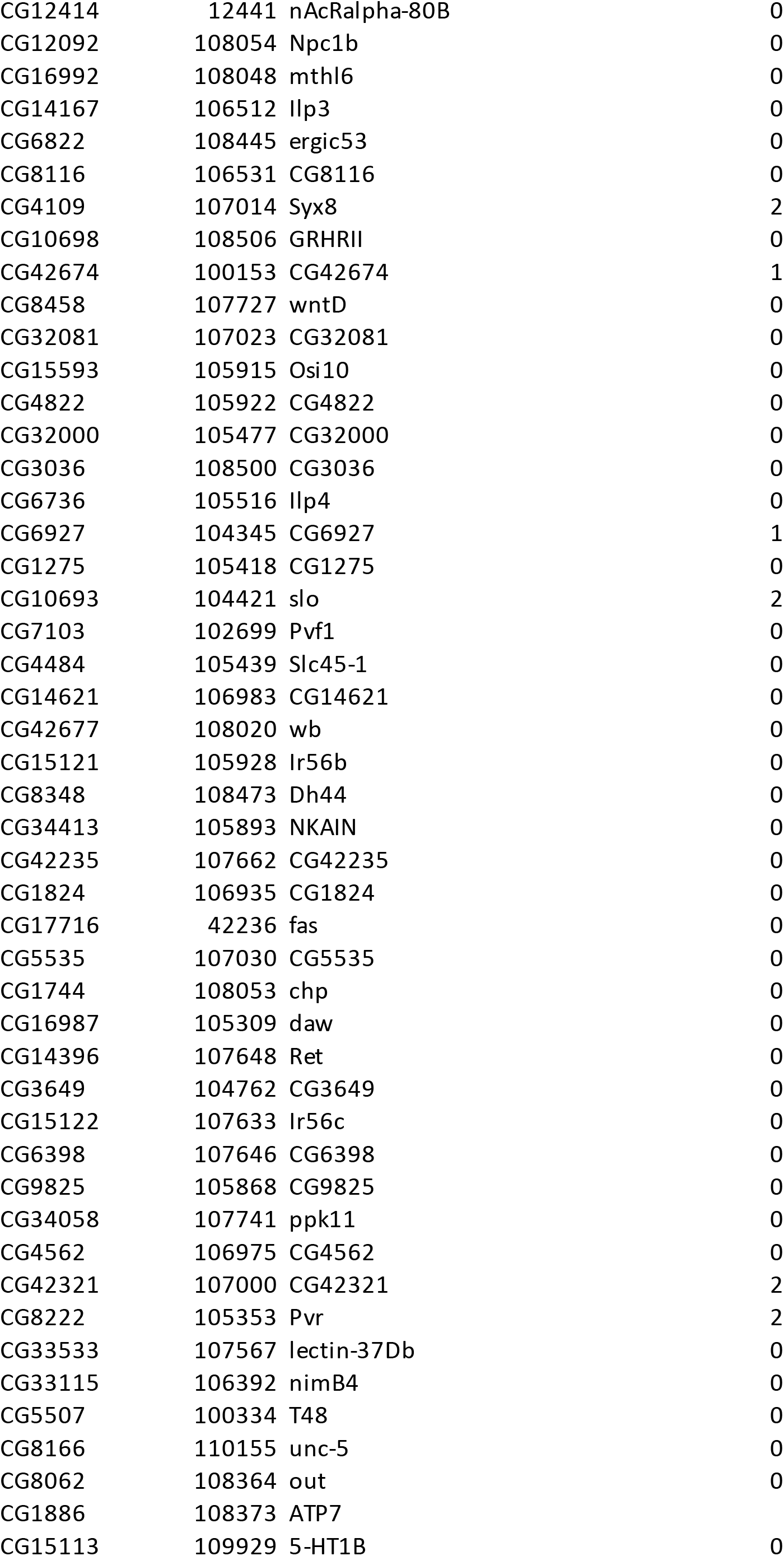
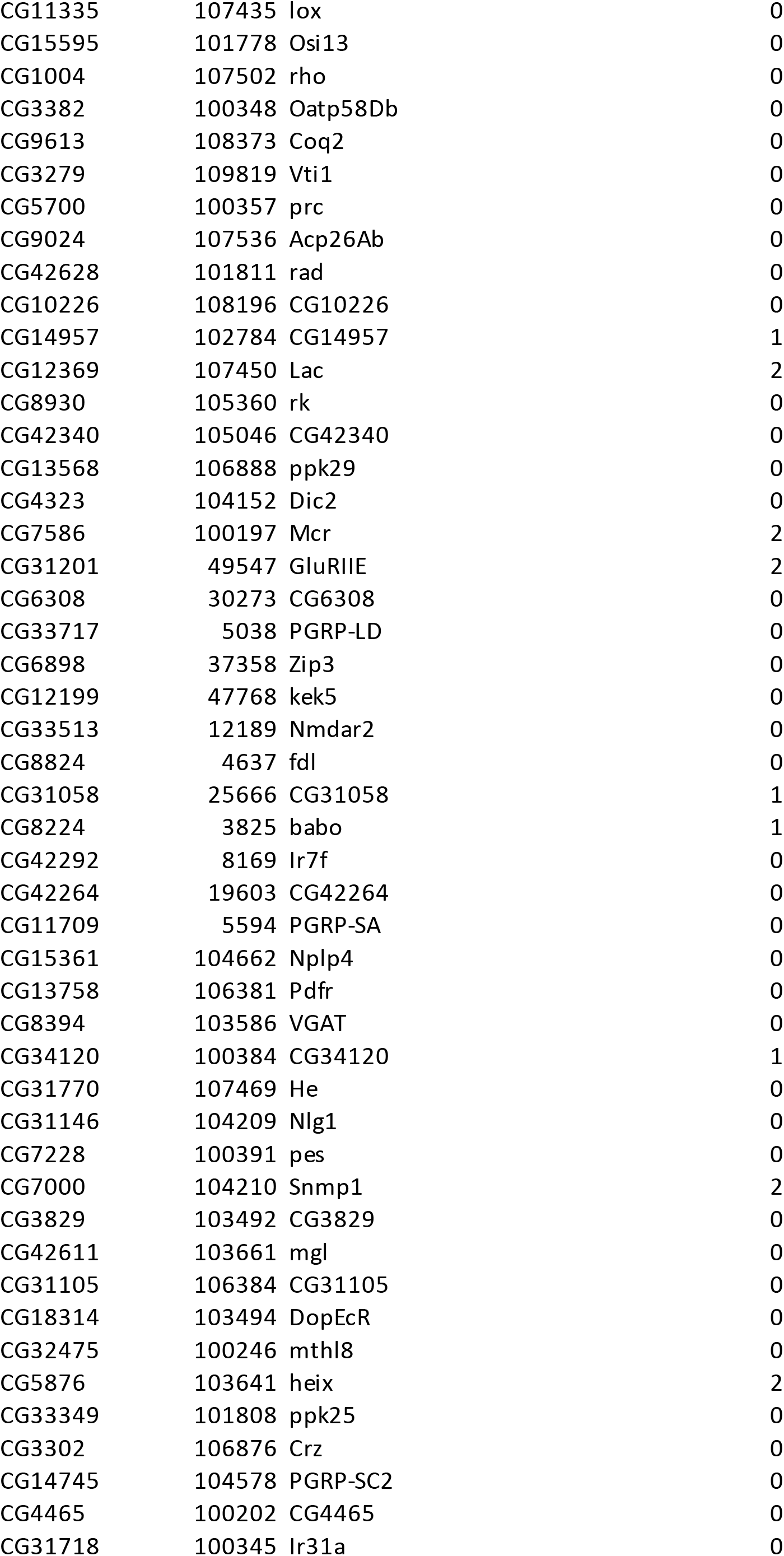
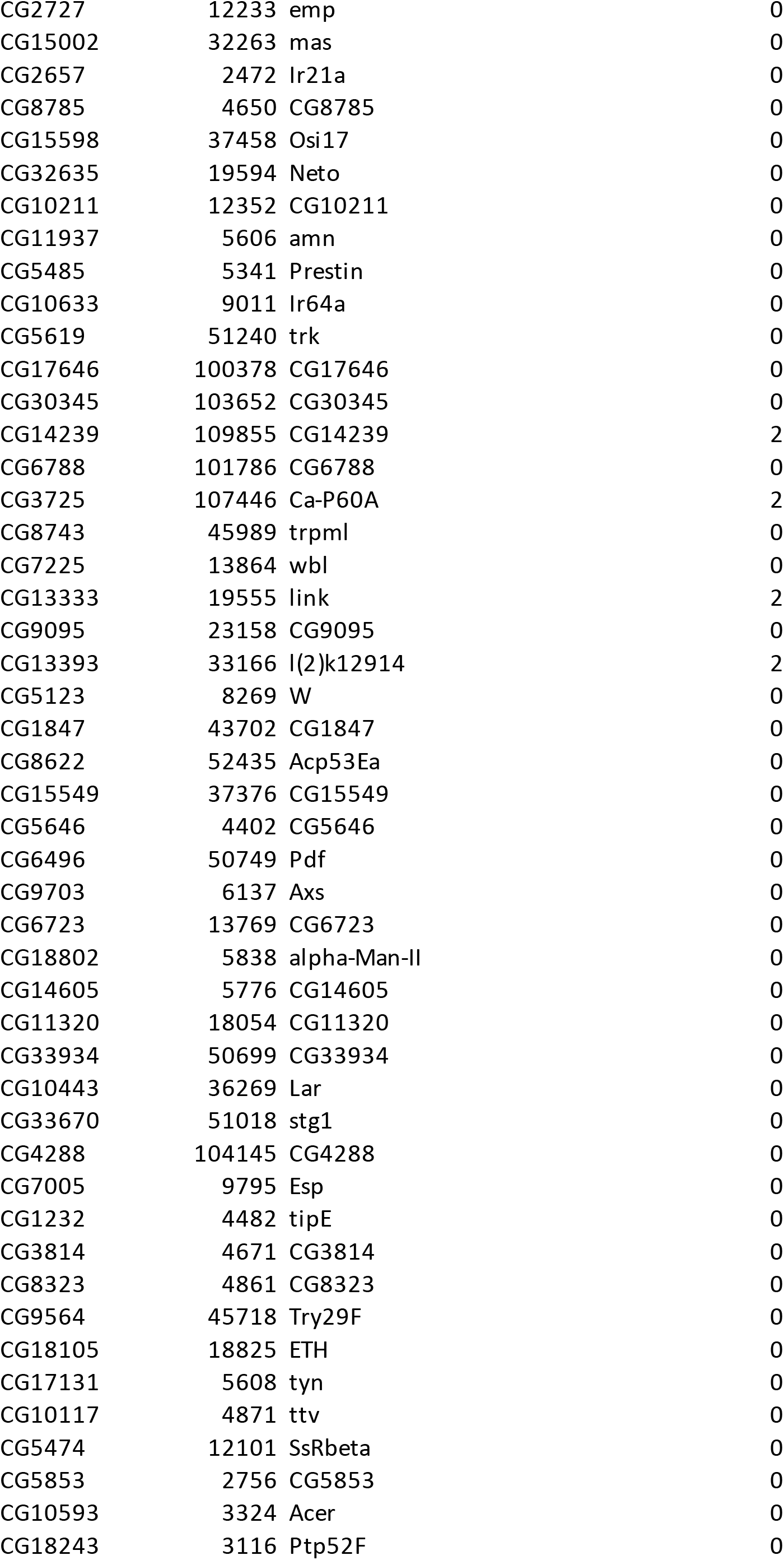
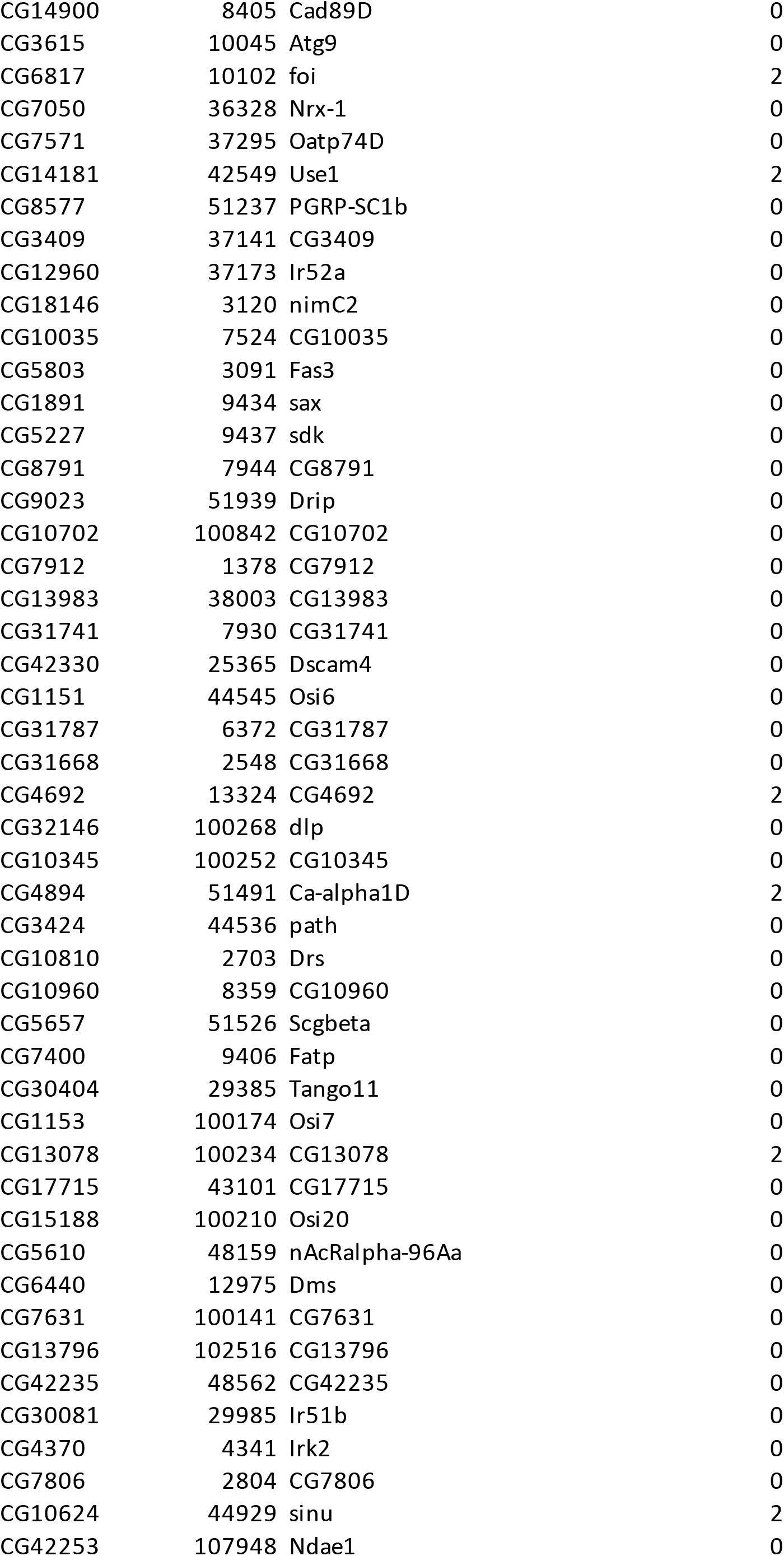
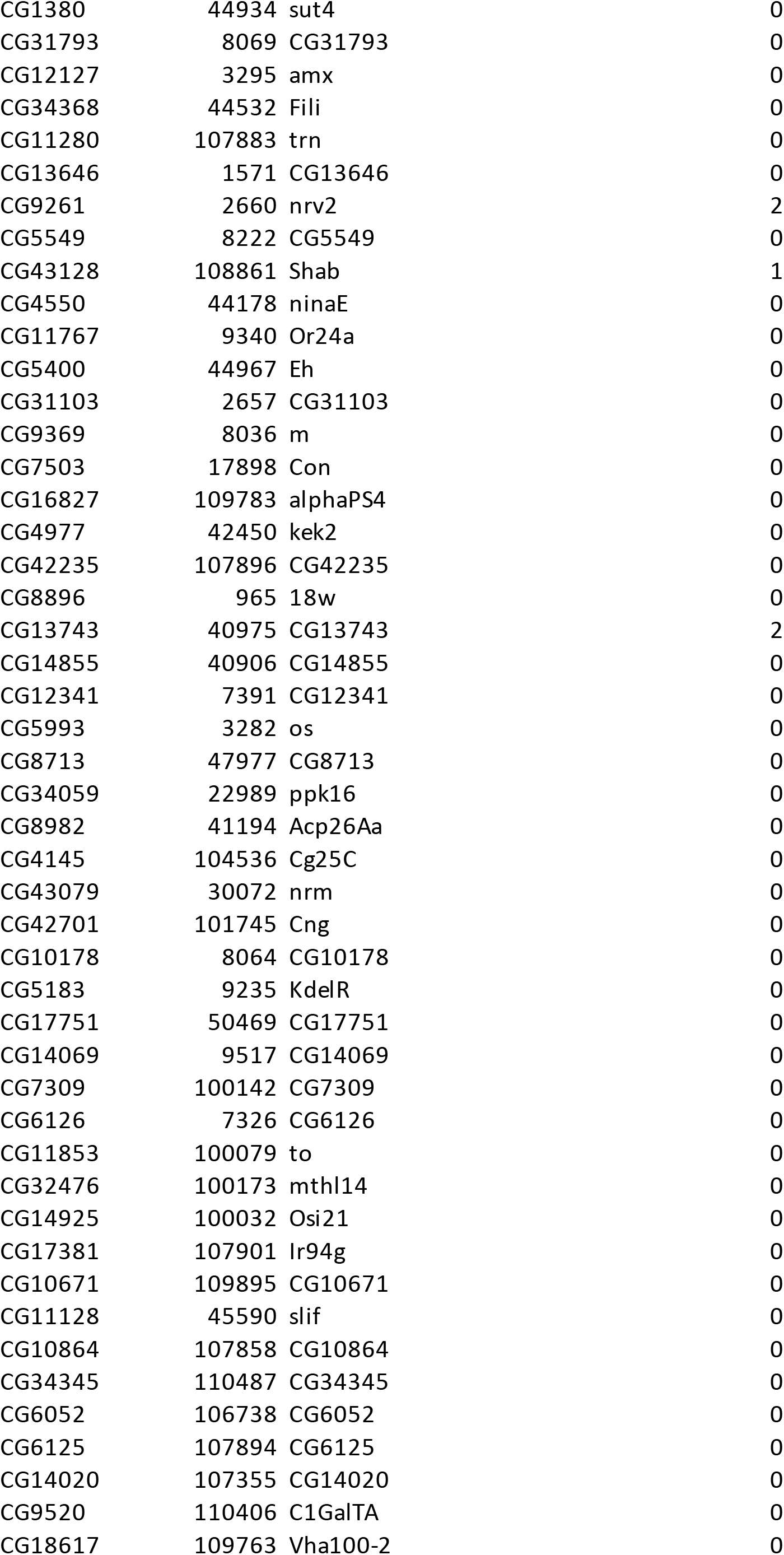
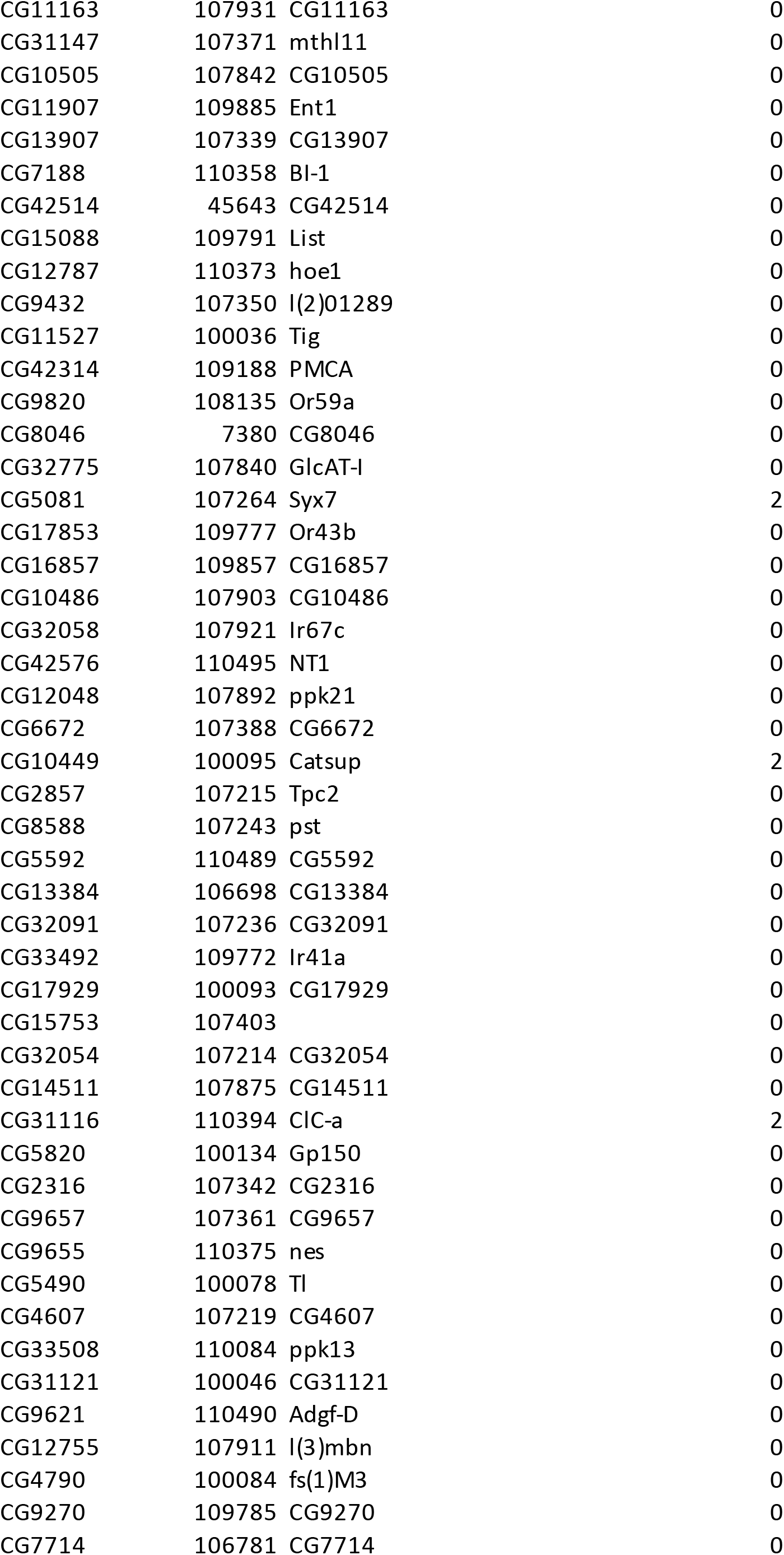
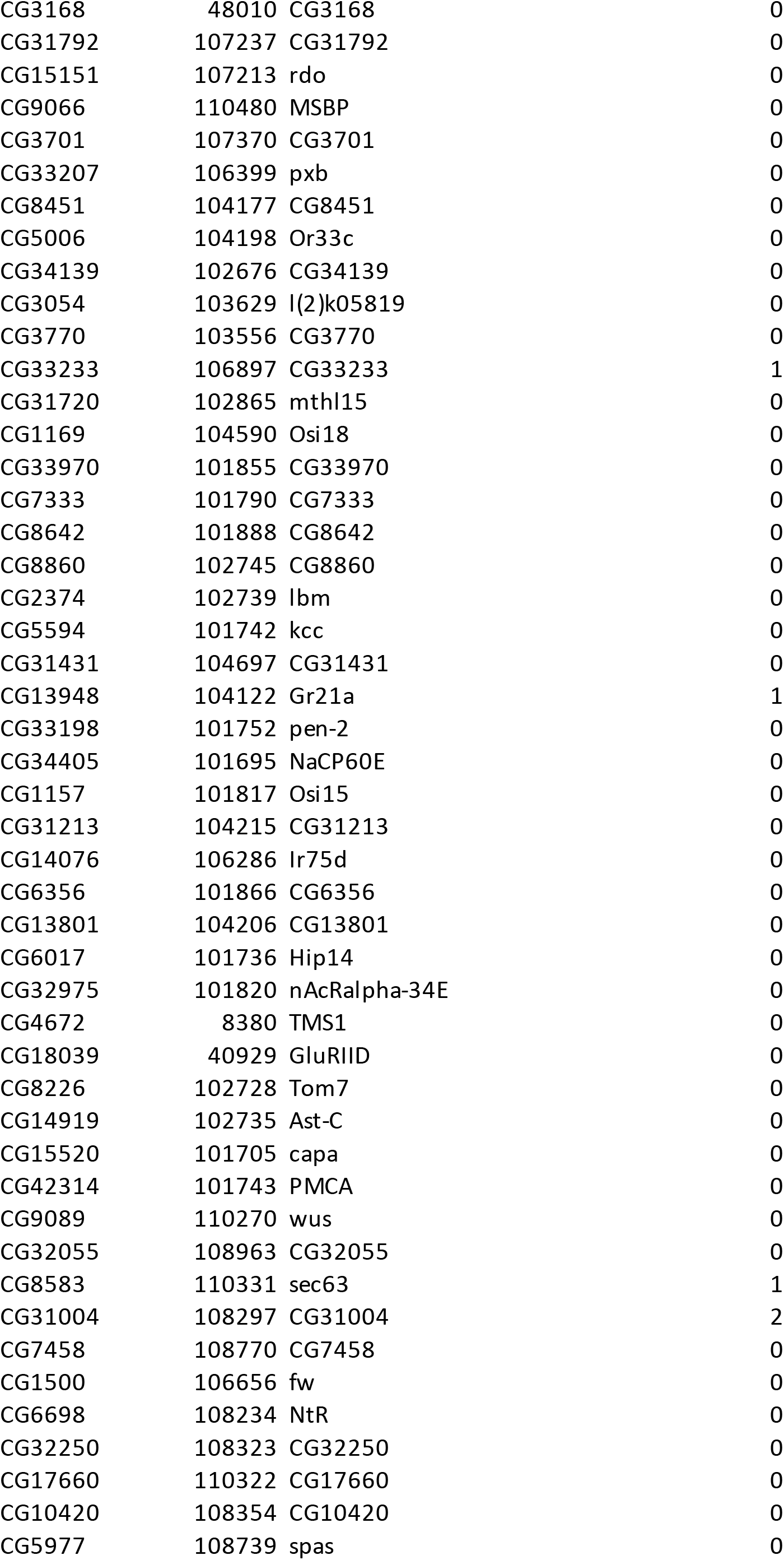
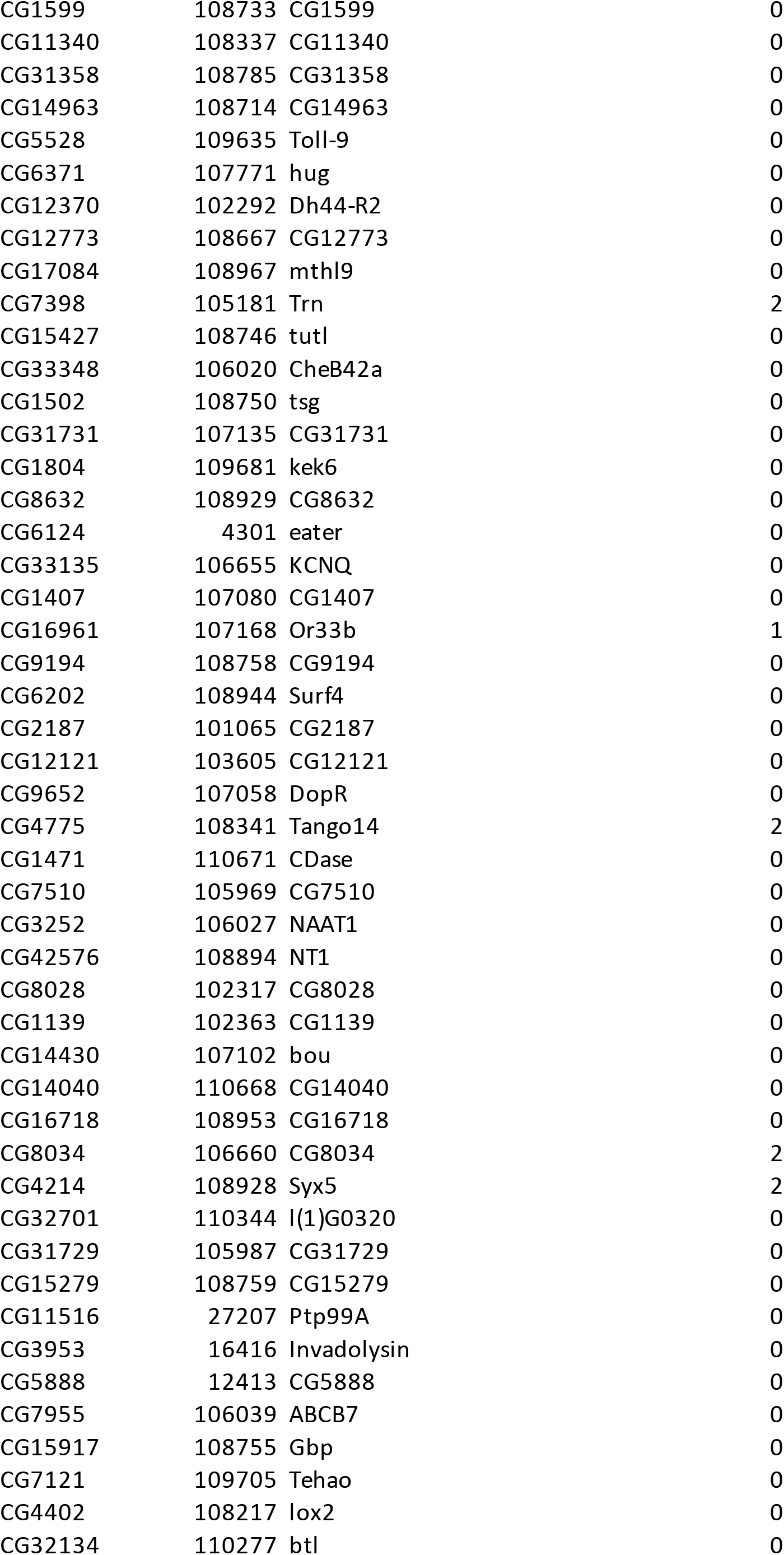
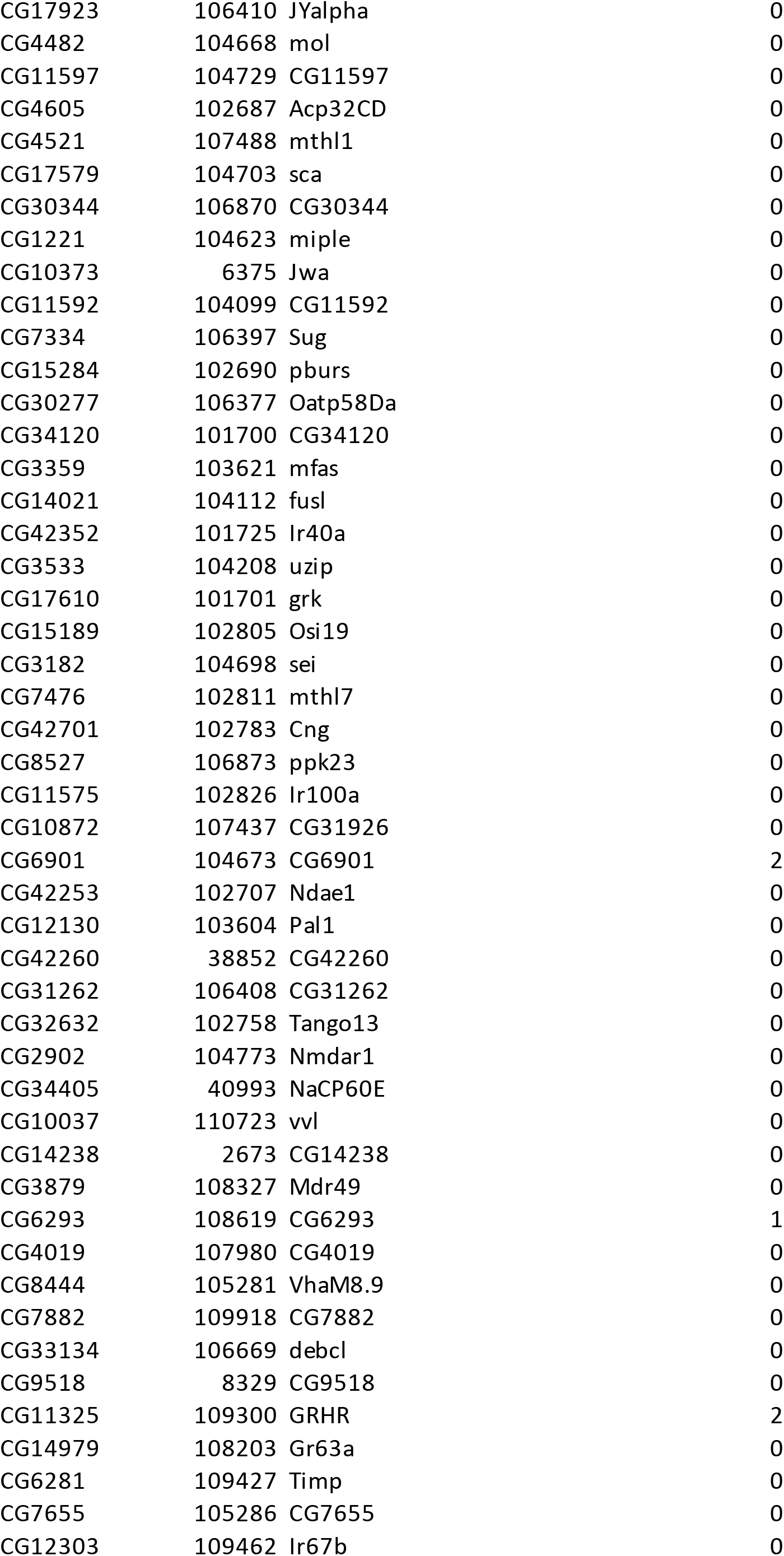
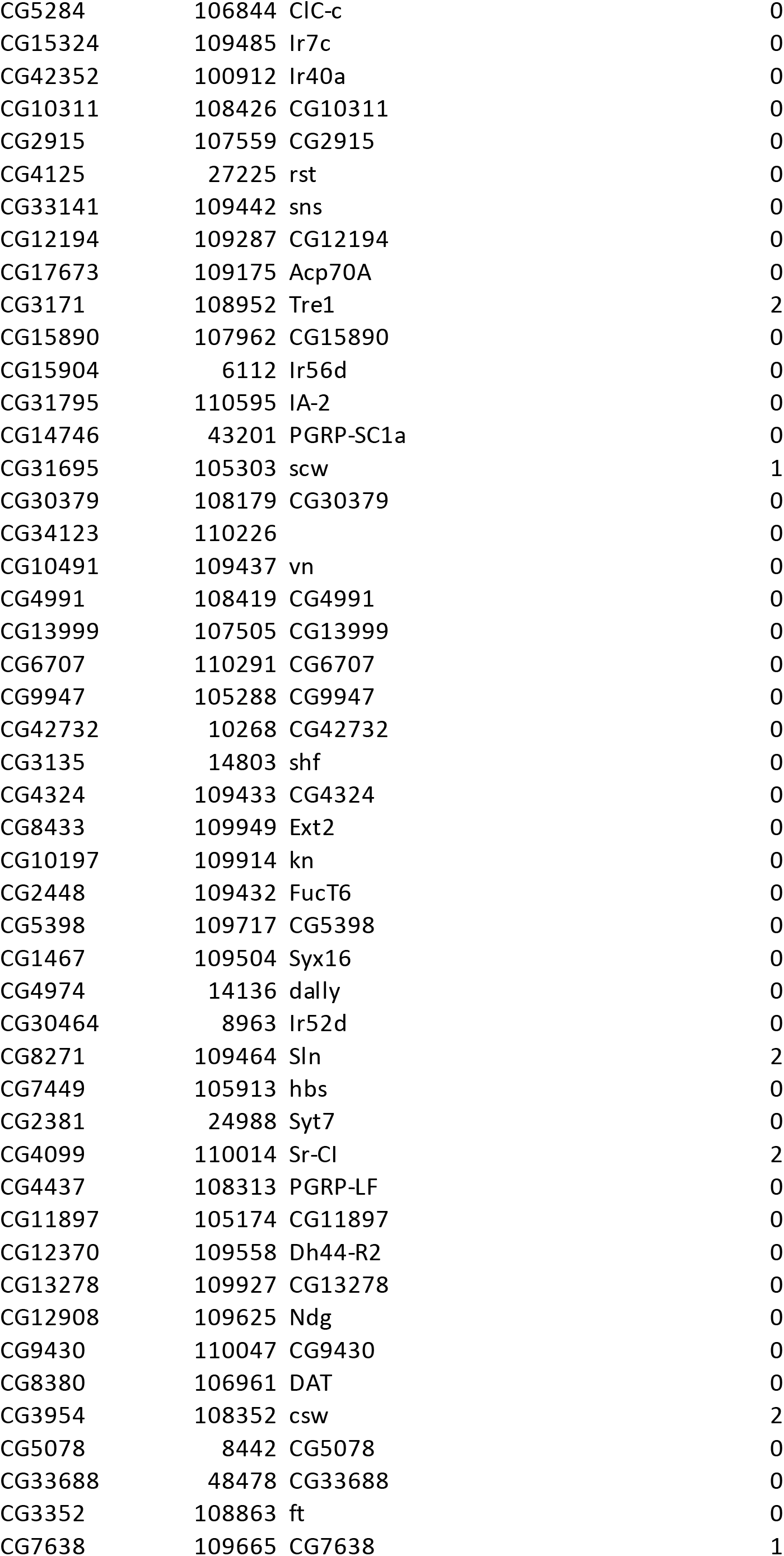
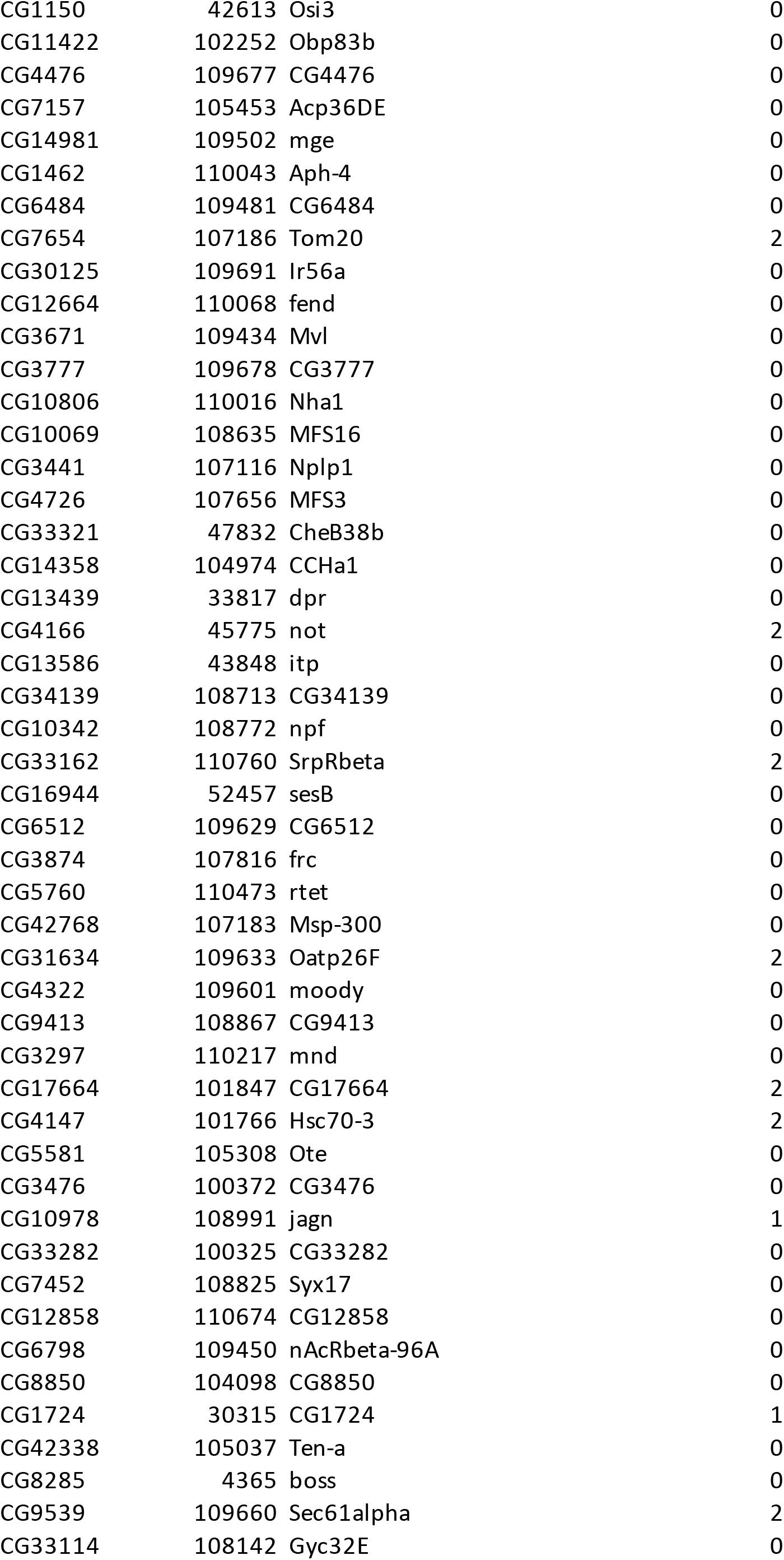
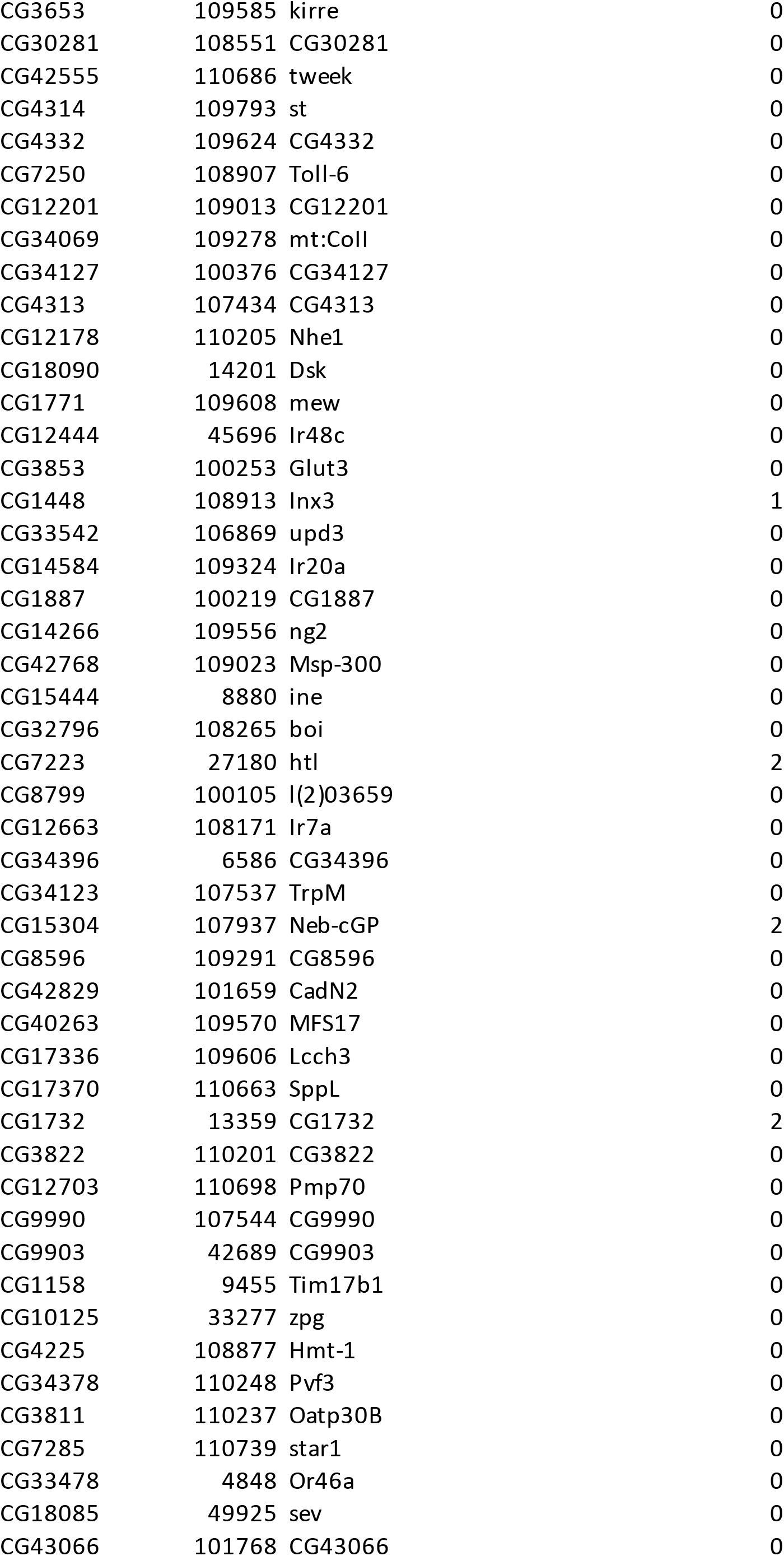
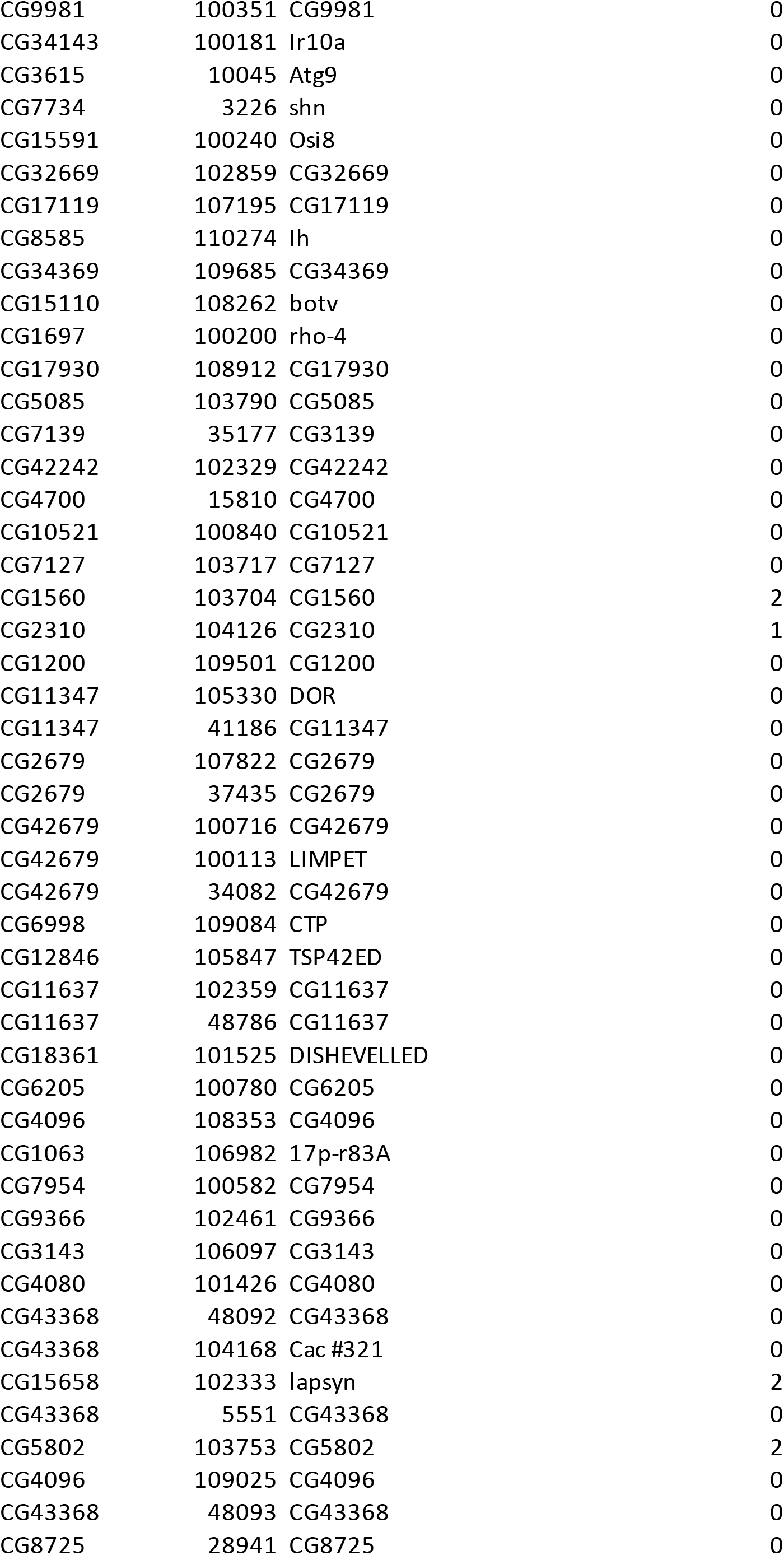
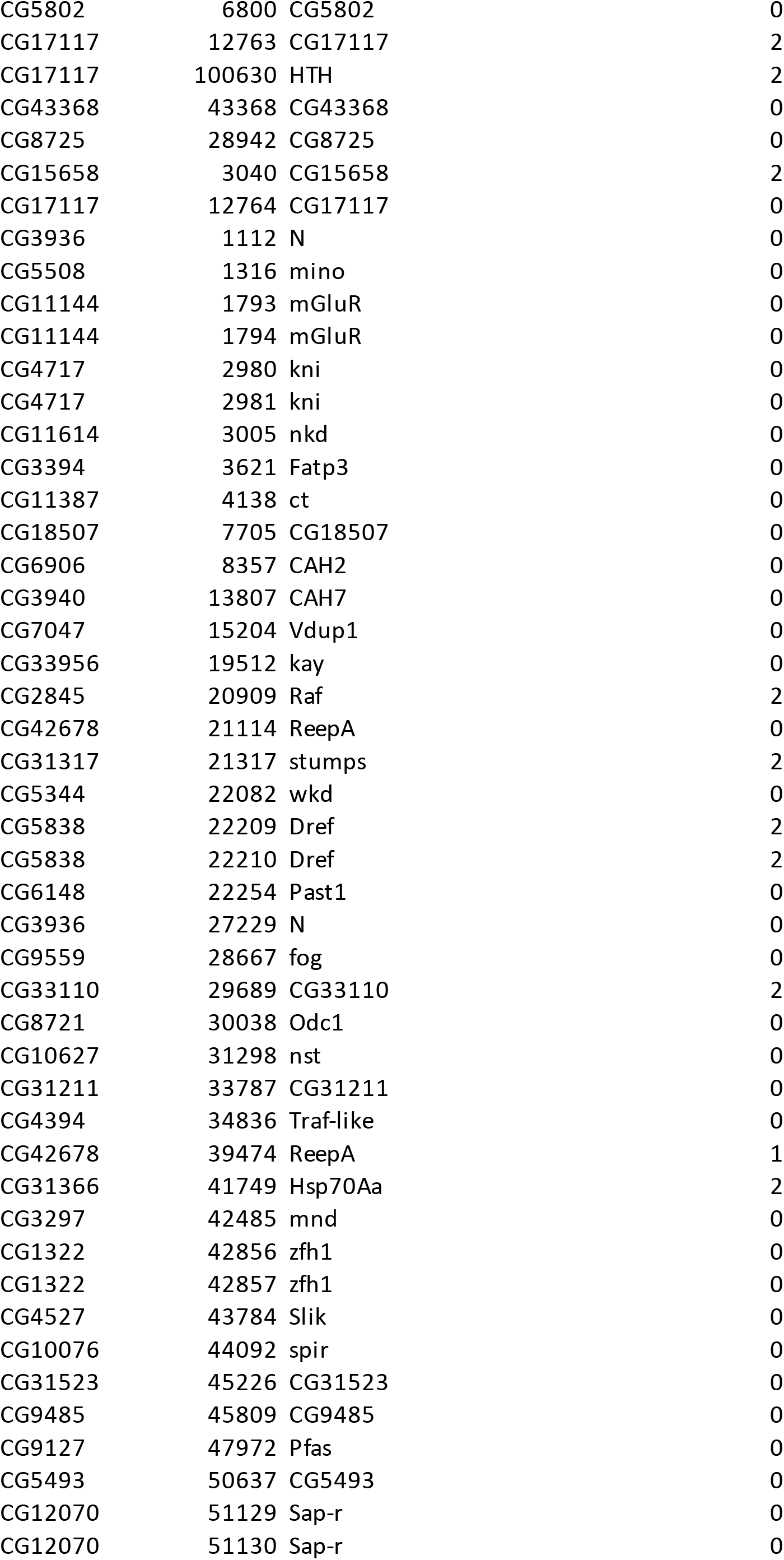
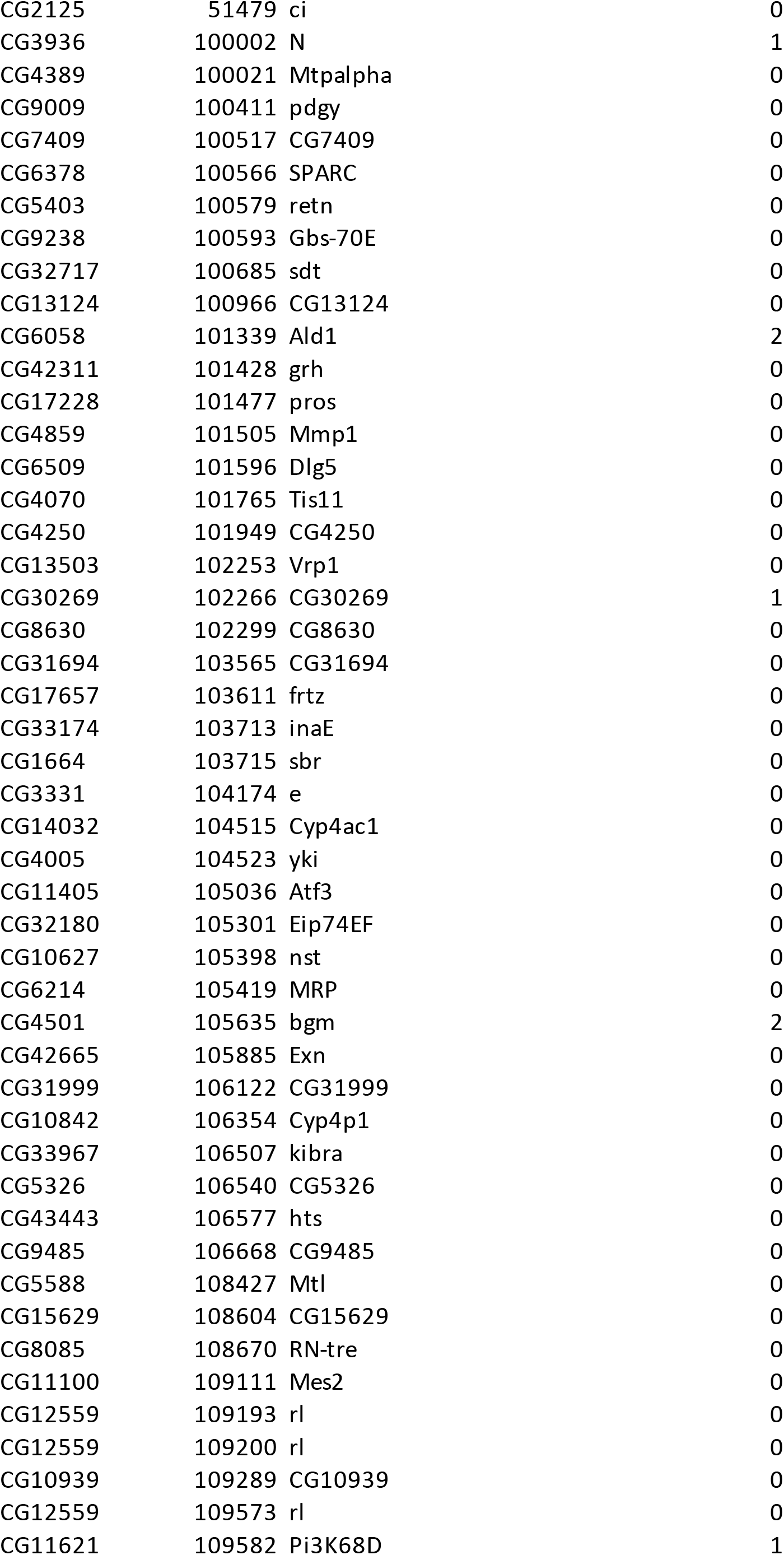
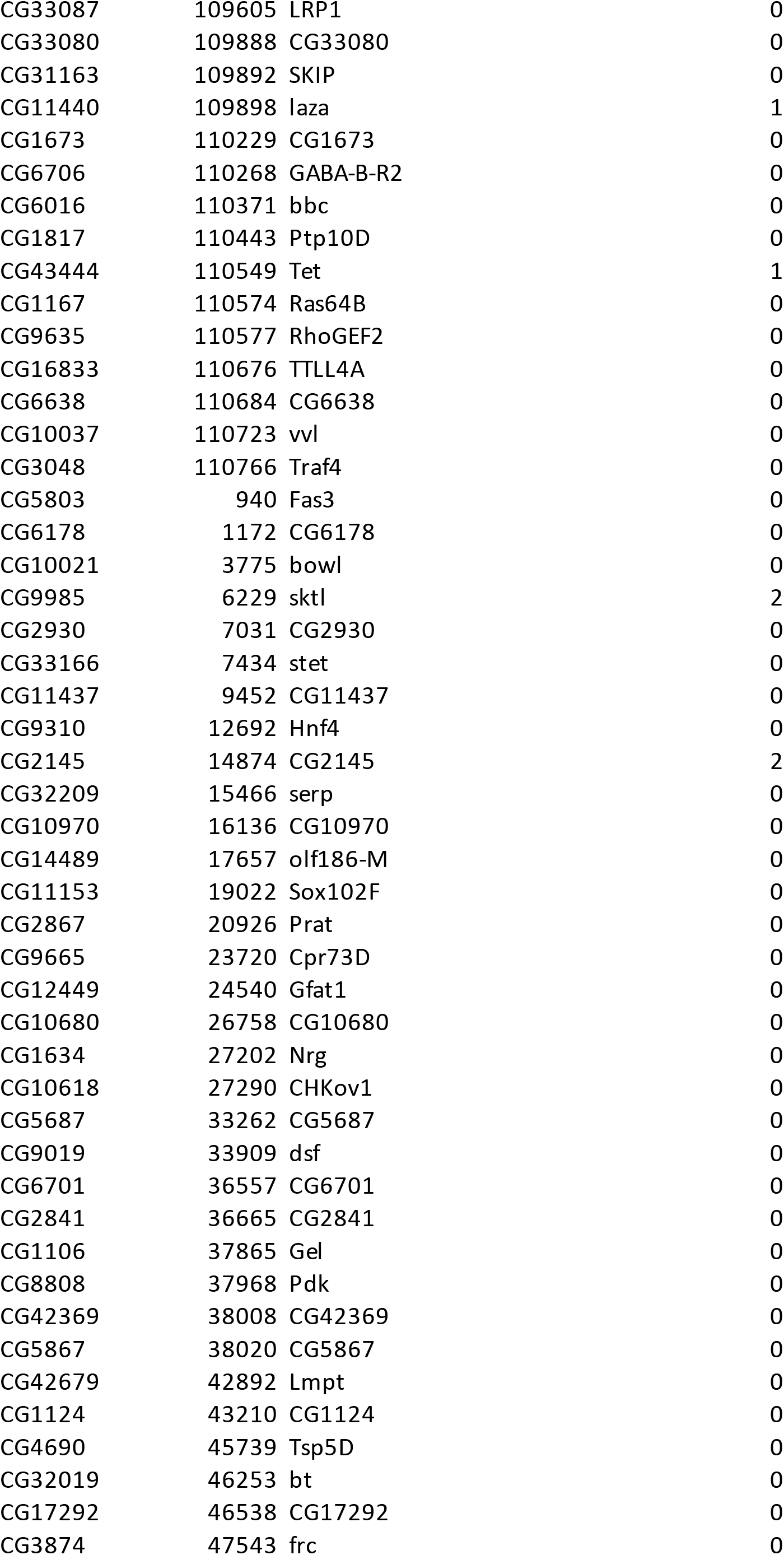
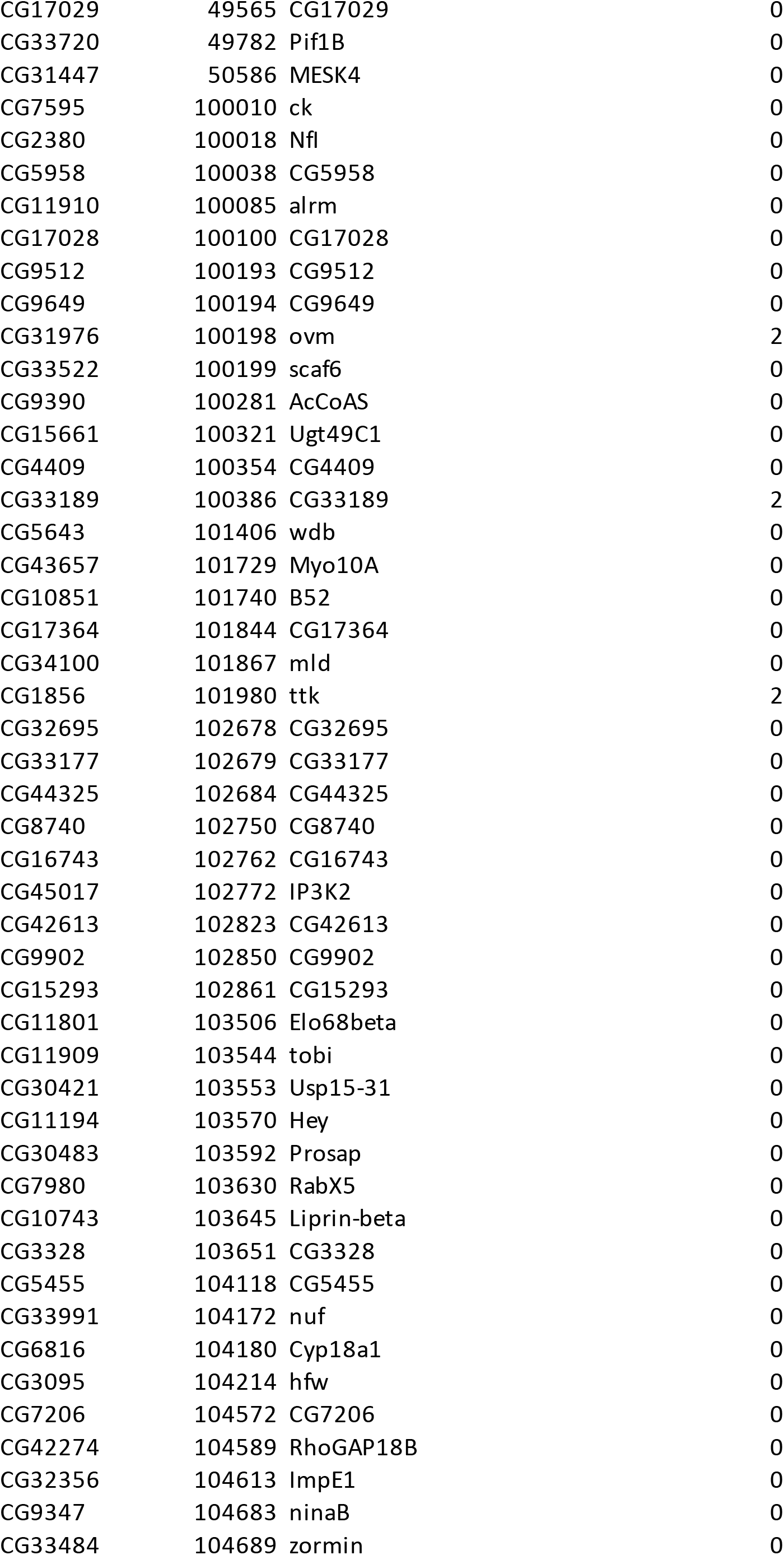
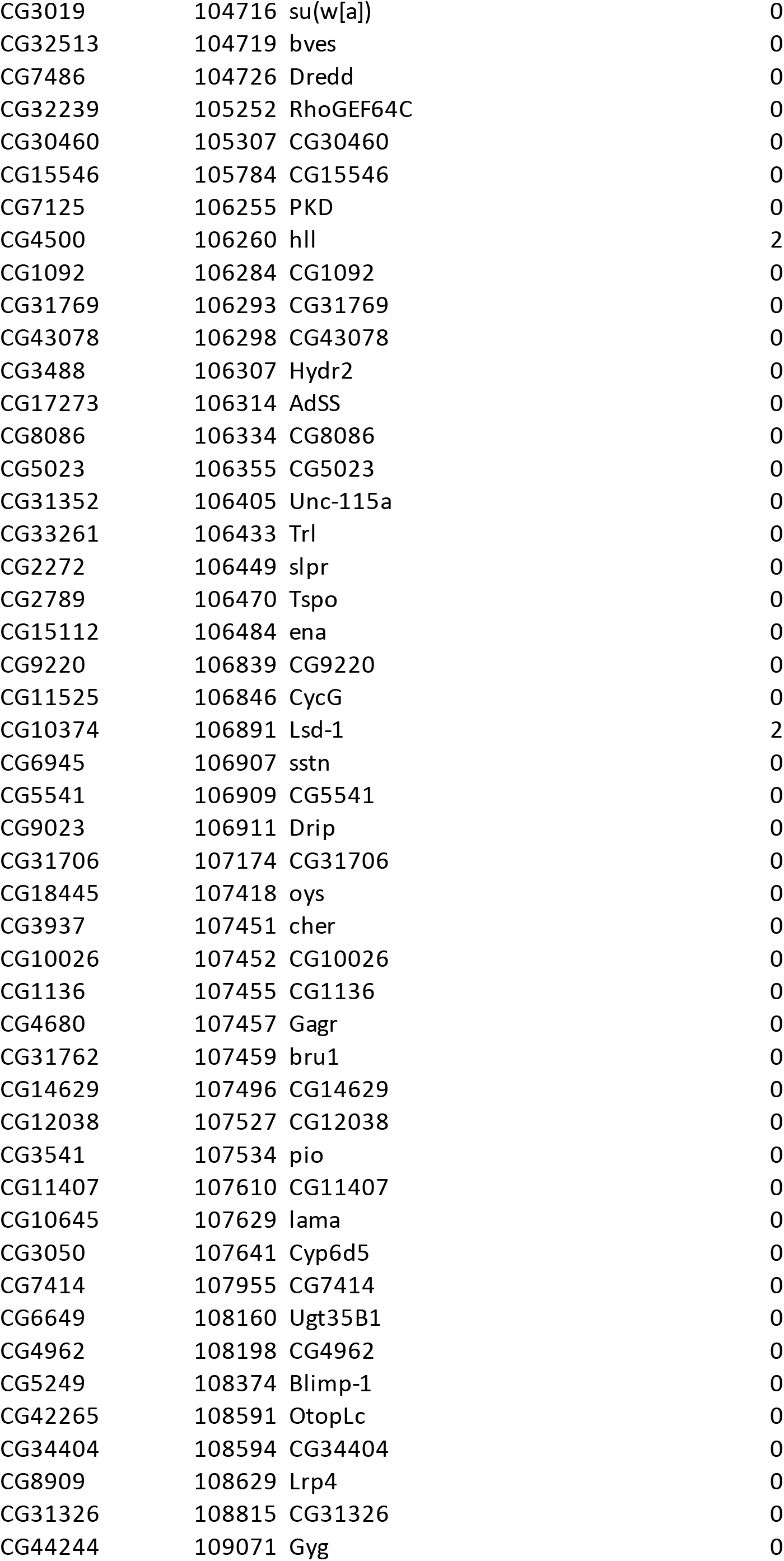
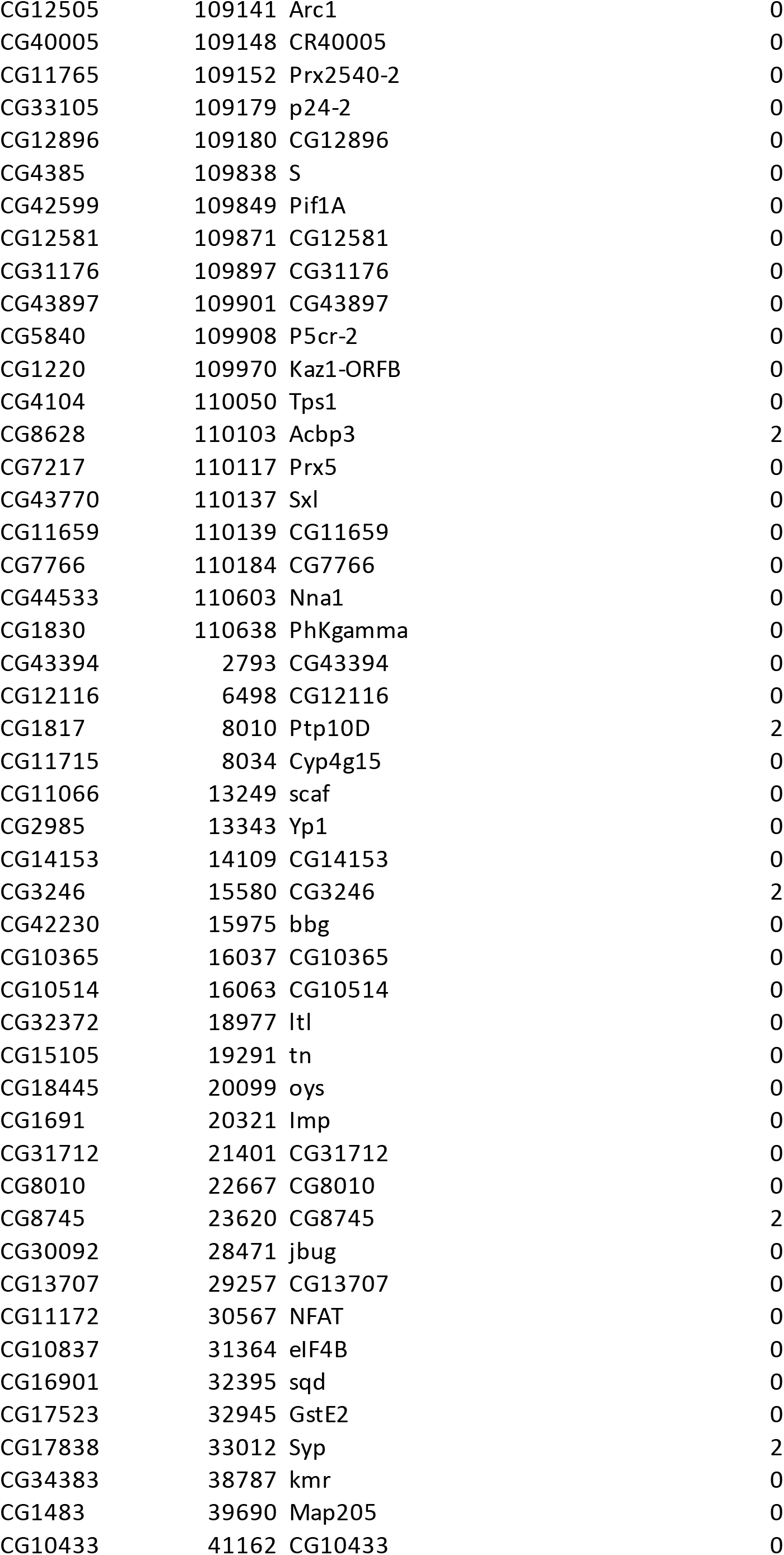
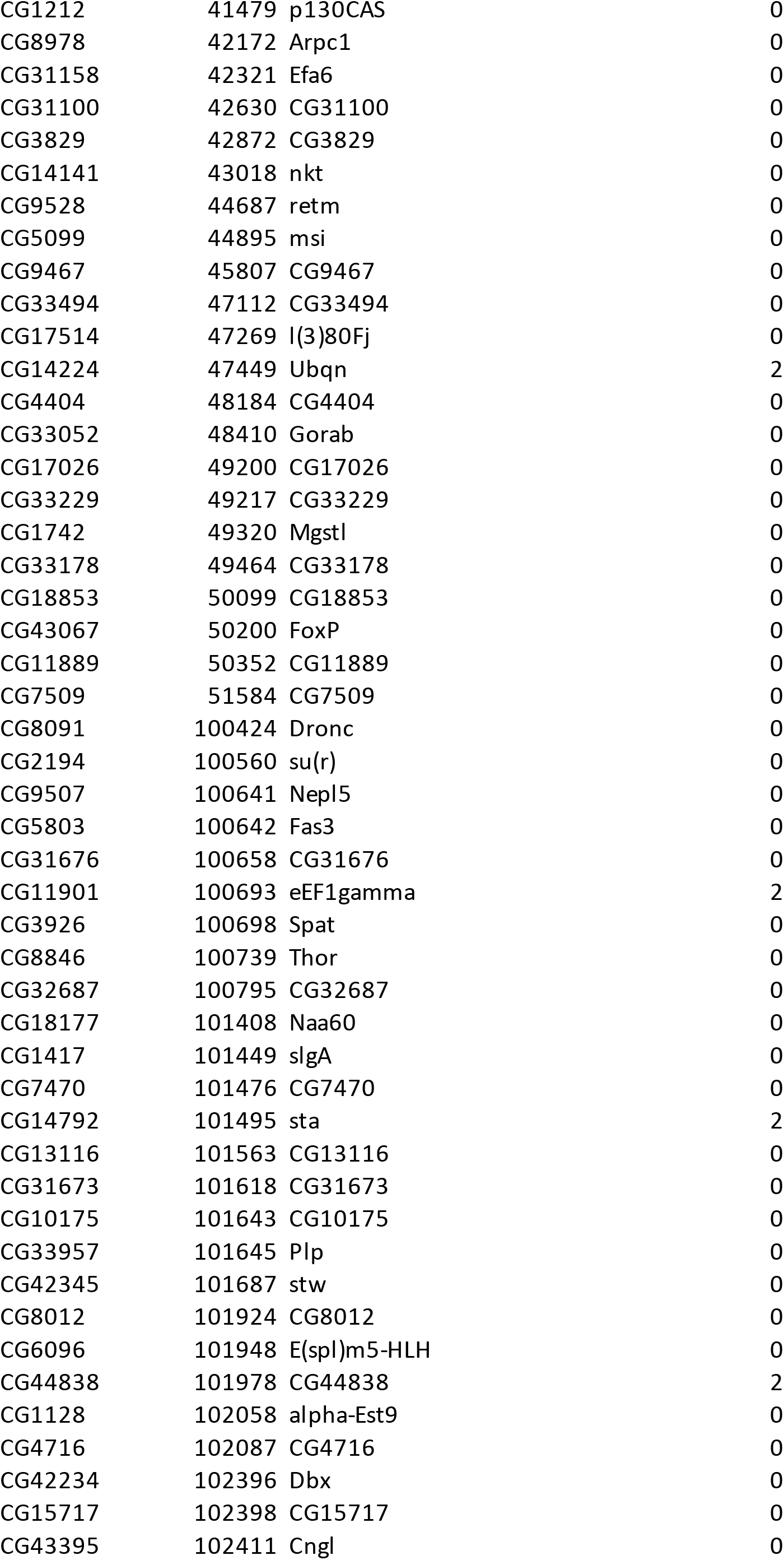
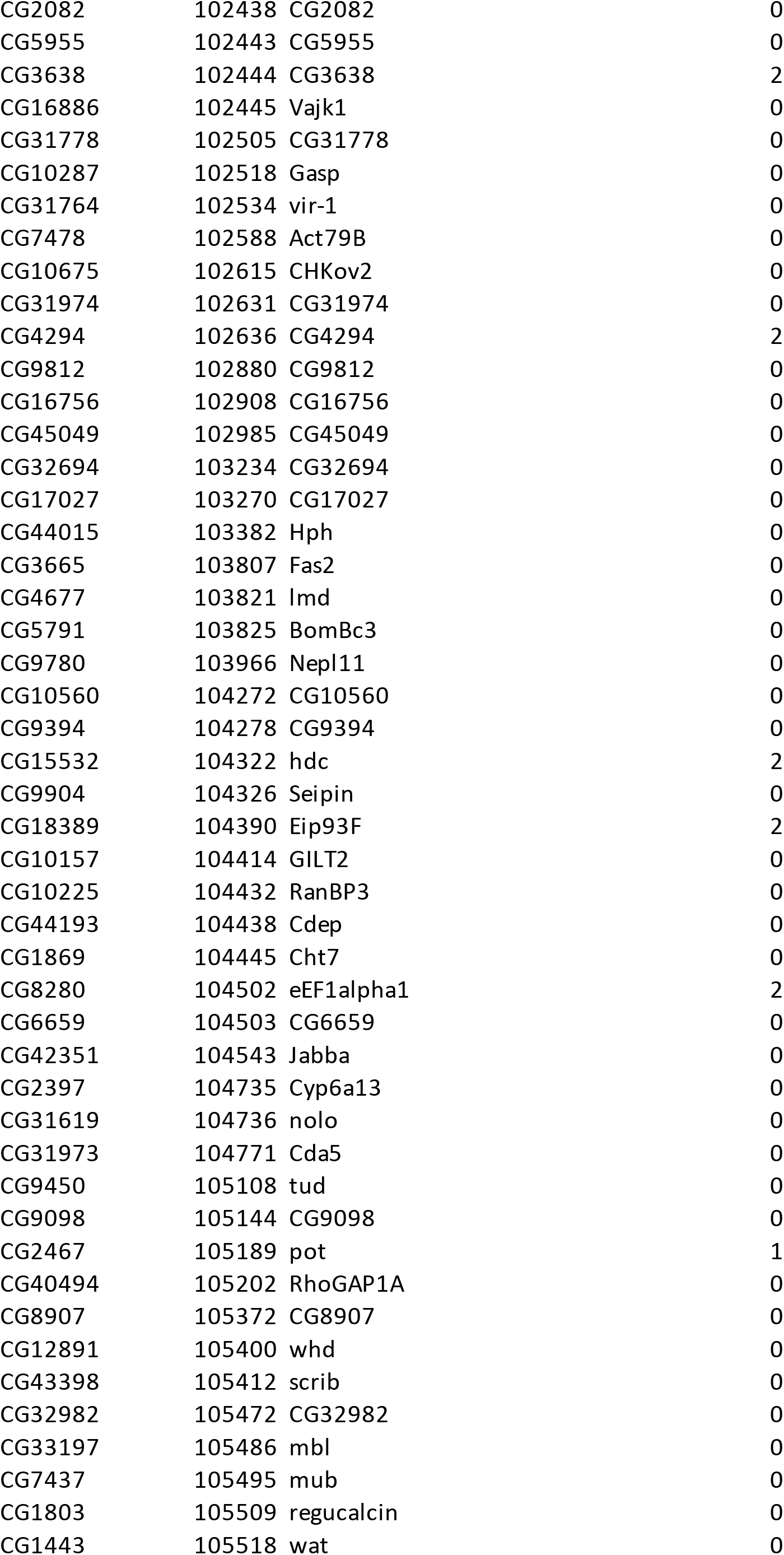
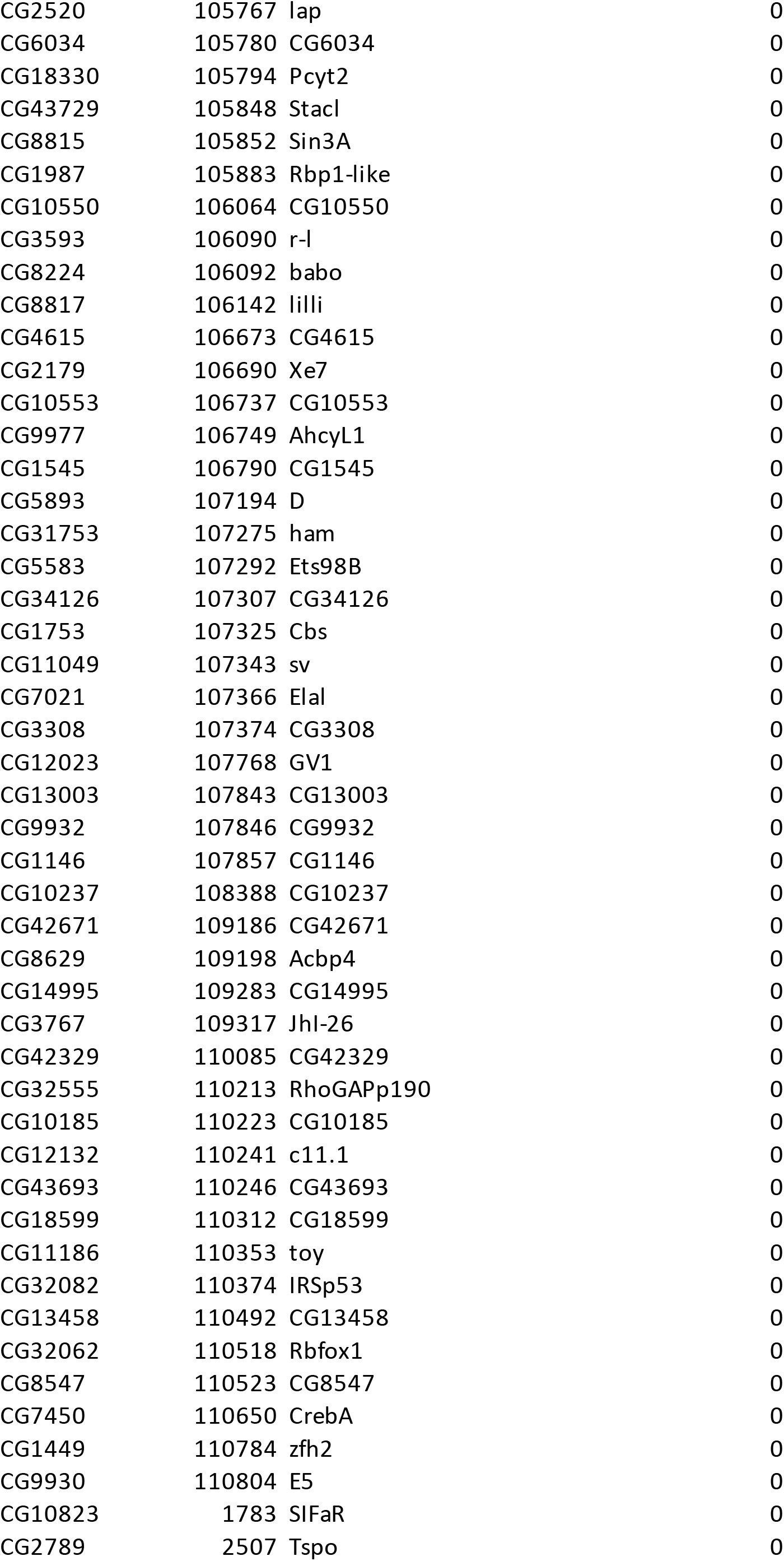
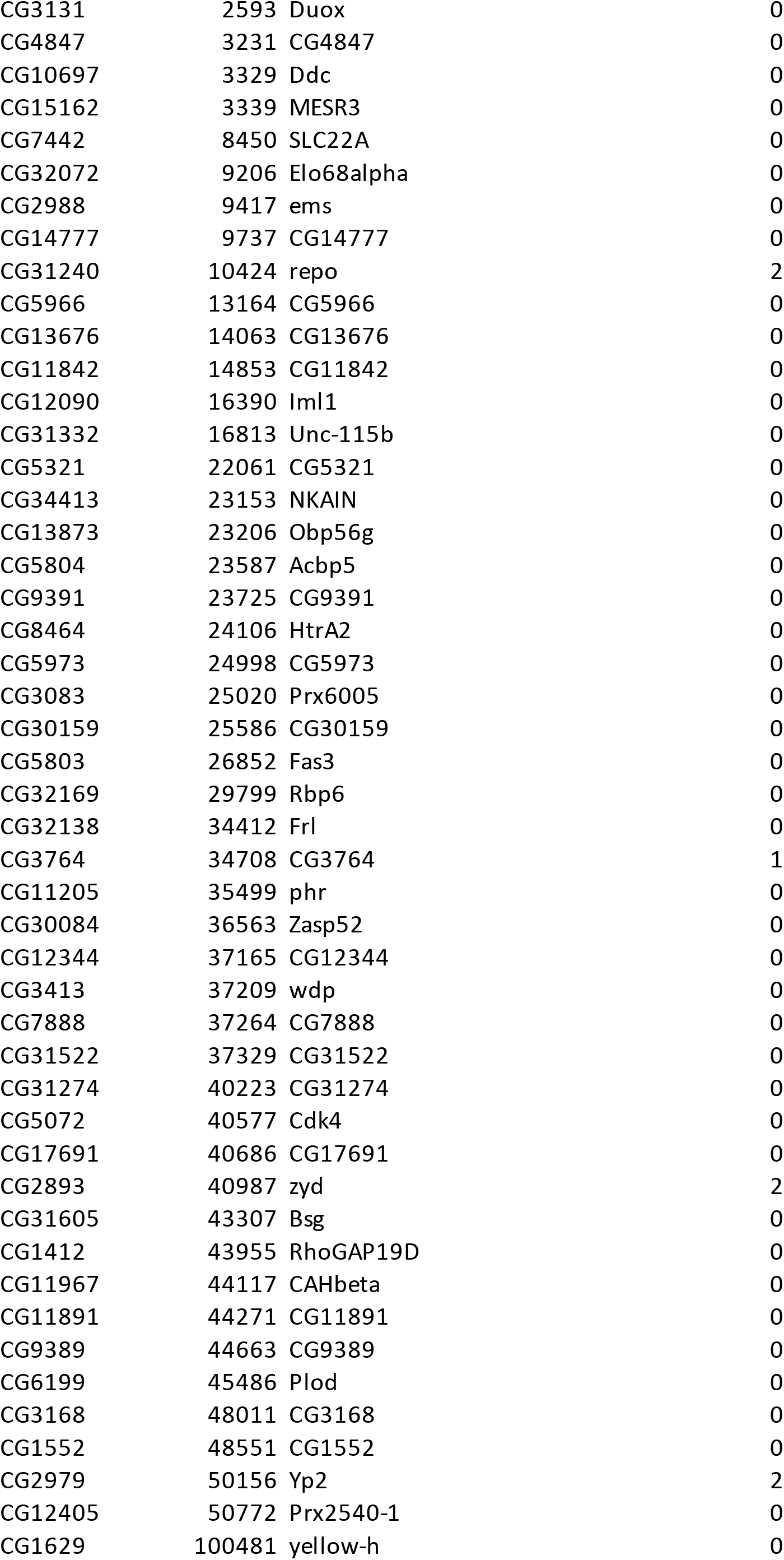
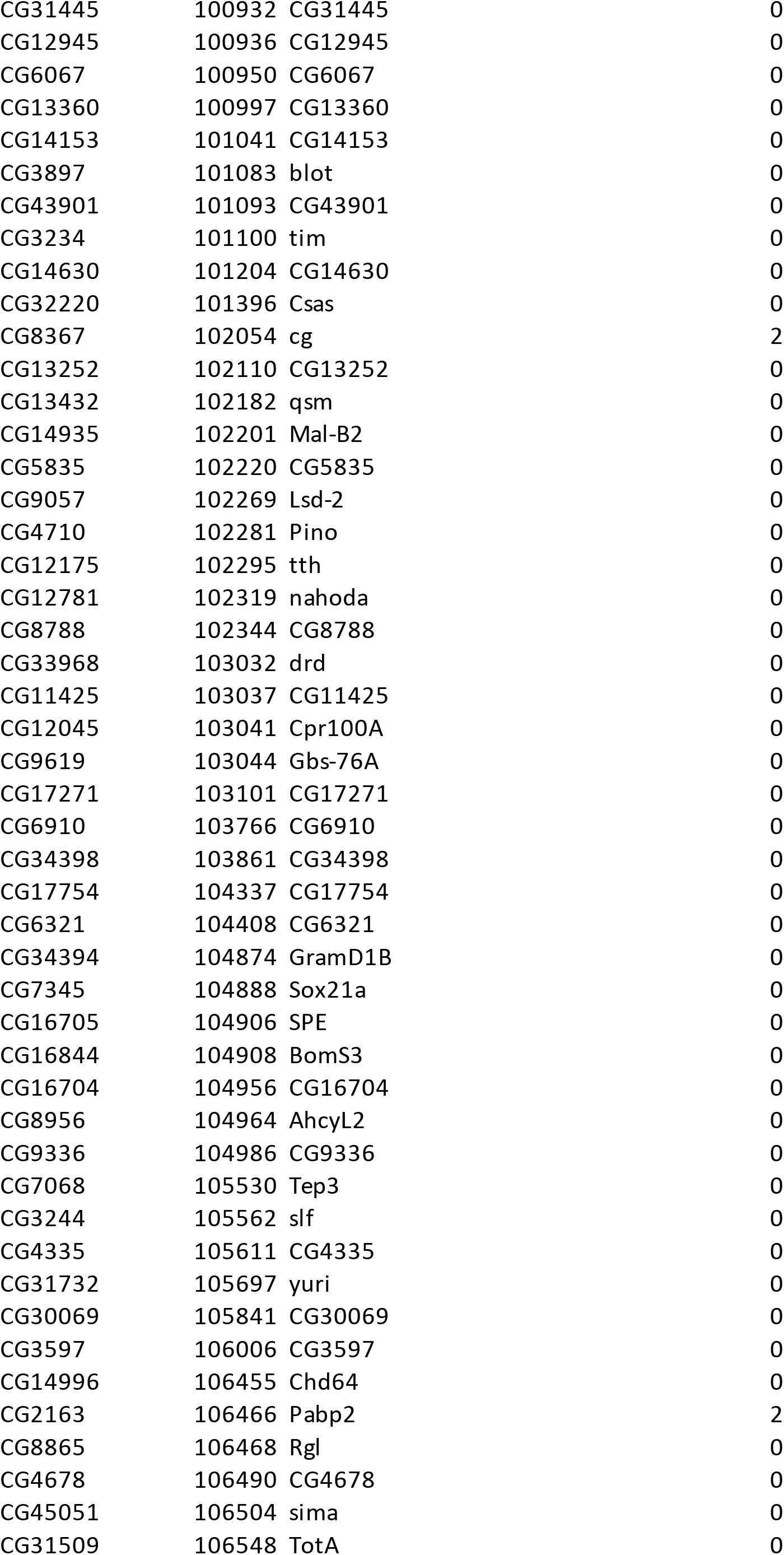
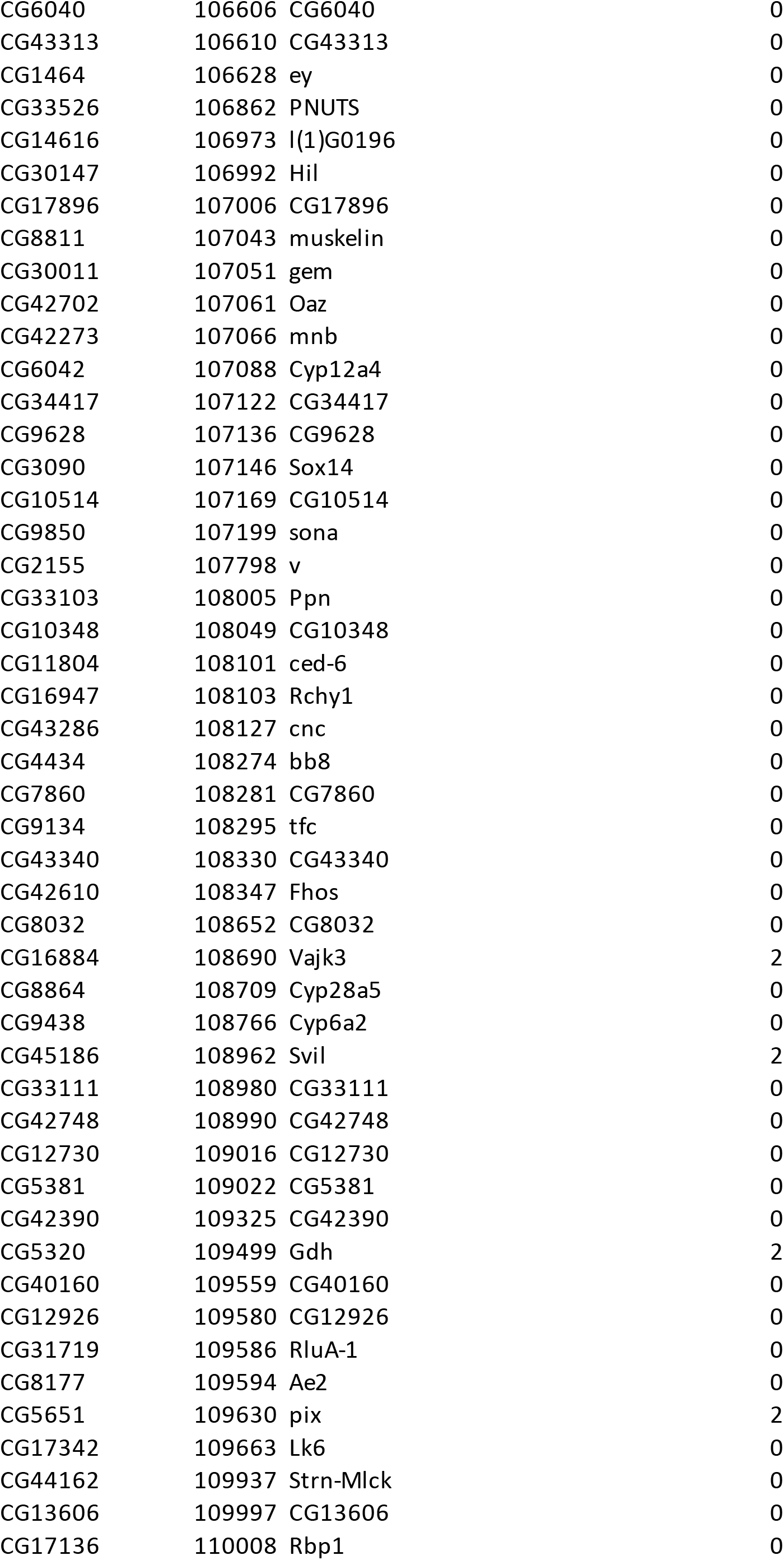
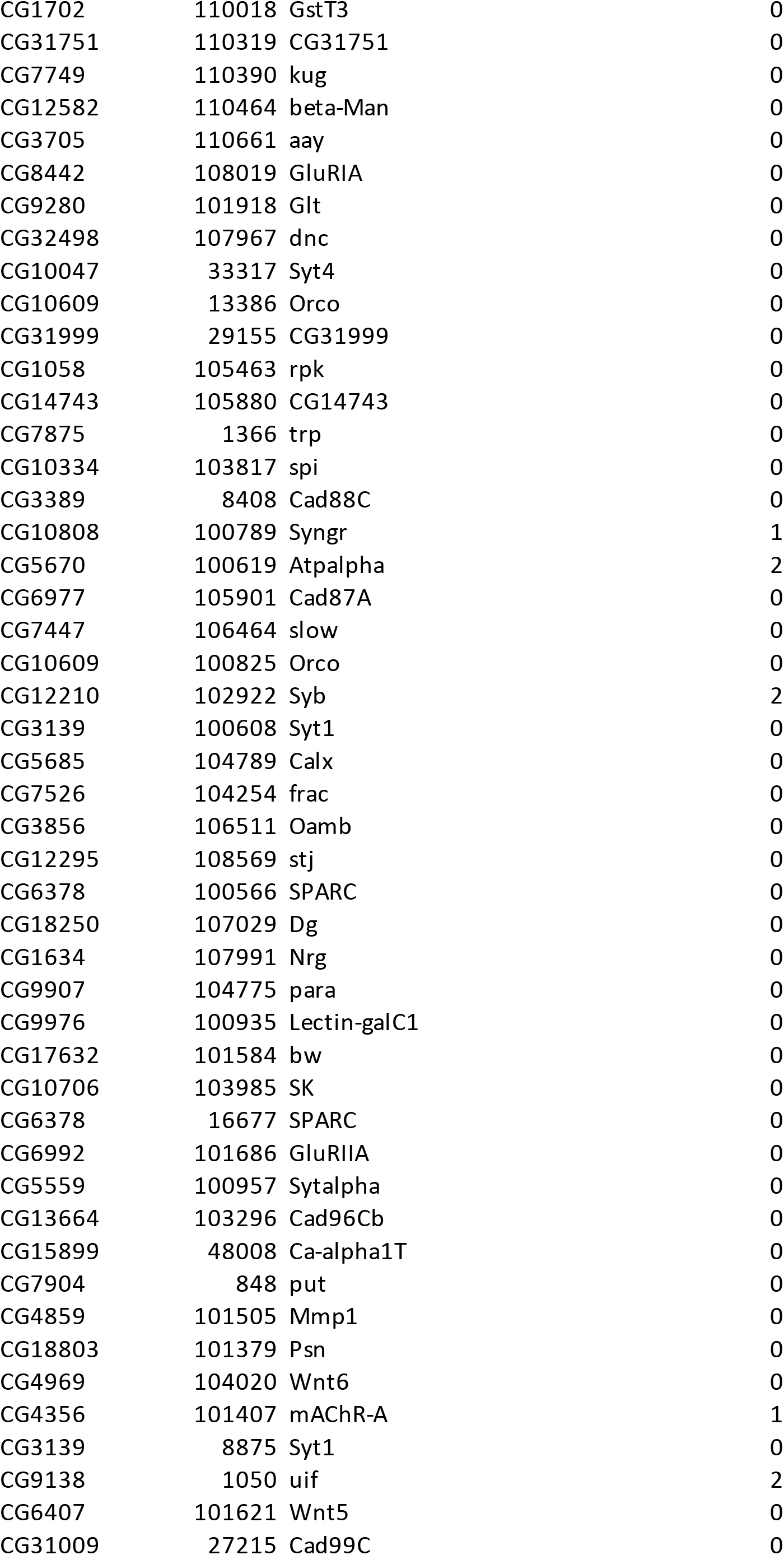
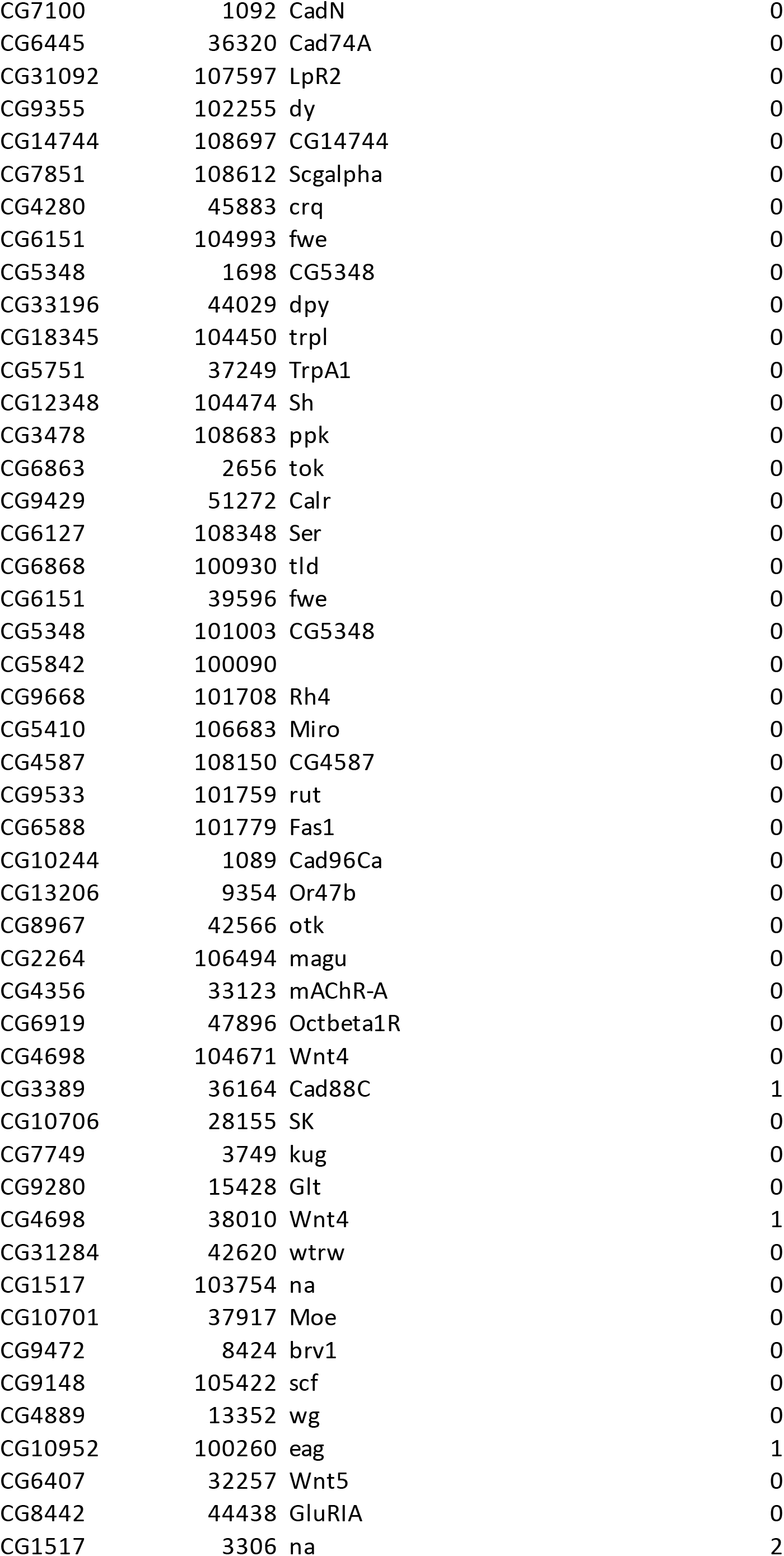
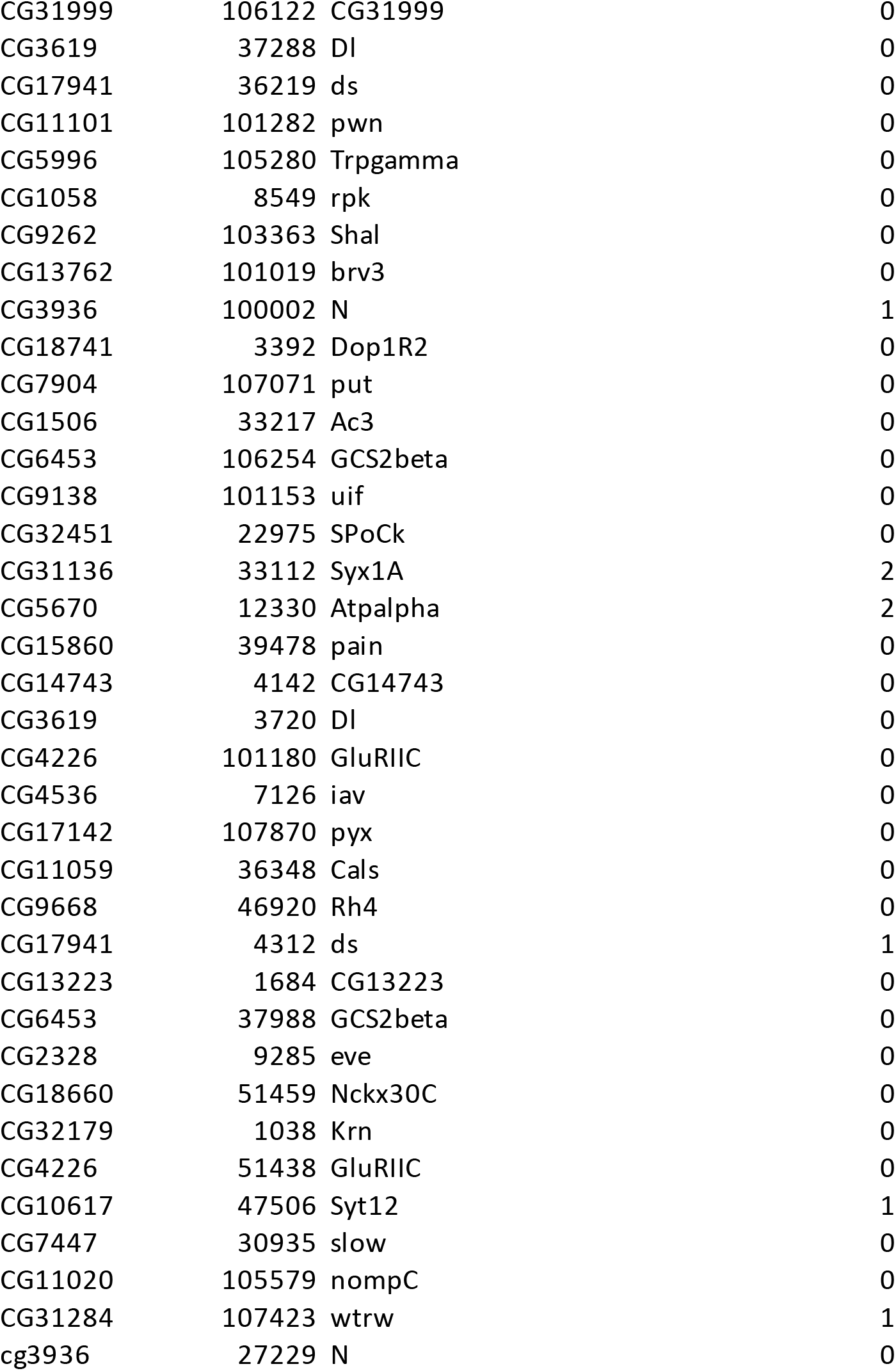

